# Nuclear pore-like structures in a compartmentalized bacterium

**DOI:** 10.1101/076430

**Authors:** Evgeny Sagulenko, Amanda Nouwens, Richard I. Webb, Kathryn Green, Benjamin Yee, Garry Morgan, Andrew Leis, Kuo-Chang Lee, Margaret K. Butler, Nicholas Chia, Uyen Thi Phuong Pham, Stinus Lindgreen, Ryan Catchpole, Anthony M. Poole, John A Fuerst

## Abstract

Planctomycetes are distinguished from other Bacteria by compartmentalization of cells via internal membranes, interpretation of which has been subject to recent debate regarding potential relations to Gram-negative cell structure. In our interpretation of the available data, the planctomycete *Gemmata obscuriglobus* contains a nuclear body compartment, and thus possesses a type of cell organization with parallels to the eukaryote nucleus. Here we show that pore-like structures occur in internal membranes of *G.obscuriglobus* and that they have elements structurally similar to eukaryote nuclear pores, including a basket, ring-spoke structure, and eight-fold rotational symmetry. Bioinformatic analysis of proteomic data reveals that some of the *G. obscuriglobus* proteins associated with pore-containing membranes possess structural domains found in eukaryote nuclear pore complexes. Moreover, immuno-gold labelling demonstrates localization of one such protein, containing a β-propeller domain, specifically to the *G. obscuriglobus* pore-like structures. Finding bacterial pores within internal cell membranes and with structural similarities to eukaryote nuclear pore complexes raises the dual possibilities of either hitherto undetected homology or stunning evolutionary convergence.

## INTRODUCTION

A nucleus surrounded by a double membrane envelope is a universal characteristic of eukaryote cells [1] and is thought to be universally absent from the prokaryote domains Bacteria and Archaea. The nucleus is accompanied by a complex apparatus for transport of macromolecules, including a multi-protein nuclear pore complex embedded in the nuclear envelope, and a soluble transport system [2]. The nuclear pore complex and many of its component proteins appear universal among eukaryotes, spanning yeast, trypanosomes and vertebrates [3], and the Last Eukaryotic Common Ancestor already possessed a complex version of the nuclear pore complex, nuclear envelope and connected endomembrane system [4,5,6]. *Gemmata obscuriglobus,* a member of the bacterial phylum *Planctomycetes*, possesses compartments including the nuclear body containing DNA and ribosomes, riboplasm containing ribosomes but no DNA, and the paryphoplasm, a ribosome-free compartment [7,8]. By whole-cell tomography and conventional transmission electron microscopy (TEM), we have previously established that the riboplasm and nucleoid compartments of *G. obscuriglobus* cells are bounded by membranes [7,8,9]. Confocal fluorescence micrographs of cells where the nuclear region has been stained with DiOC6 membrane stain and DAPI DNA stain are also consistent with the membrane-bounded nature of the DNA in this organism [10]. An earlier study of *G. obscuriglobus* internal membranes differs in its conclusions from those of ours, proposing that only one invaginated membrane exists in such cells and that there is no membrane enclosure of the *Gemmata* chromosome [11], and instead a tubulovescular model for internal membranes has been proposed [12]. Our tomography analysis of *G.obscuriglobus* cells demonstrated that the internal membranes do not display continuity with the cytoplasmic membrane apposed to the cell wall [7]. Such a cell plan implies specialized internal membrane(s) distinct from the cytoplasmic membrane and would also require some form of transport system (e.g. pore structures) for macromolecules passing between the internal compartments and the rest of the cytoplasm. This hypothesis is consistent with the recent finding of confinement of translation to non-nucleoid regions of *G. obscuriglobus* cells [13]. A corollary of our study of *G. obscuriglobus* internal compartments was that several different types of membranes might be isolatable from lysed cells, and we have confirmed this concept here. There has been extensive debate regarding the evolutionary significance of compartmentation in *G. obscuriglobus* [14,15,16]. However, such discussions have been limited by lack of knowledge about *Gemmata* membrane composition and the structure of internal membranes in particular. Recently, components of cell walls characteristic for Gram-negative bacteria such as peptidoglycan and lipopolysaccharide have been found in *Gemmata obscuriglobus* [17,18] correlating with other data on occurrence of peptidoglycan in *Planctomyces limnophilus* [17]and an anammox planctomycete species[19]. The exact location of these components within planctomycete is yet unknown, but the results suggest a potential for planctomycete cell plan to relate more closely to Gram-negative cell wall and structure than previously thought as outlined in published hypotheses[20,21]. The implications of these results for interpretation of planctomycete internal membranes and their evolutionary significance are not yet clear. Here we present evidence that some of the internal membranes of *G. obscuriglobus* possess pores with complex structure. Moreover we identify proteins specific to these membranes, some of which possess structural domains also found in eukaryote nucleoporins. The evolutionary implications of these results are considered, both from the perspective of common ancestry with the eukaryote nuclear pore complex, and from the viewpoint of convergent evolution.

## RESULTS AND DISCUSSION

### Planctomycetes possess pores in internal membranes

Pore-like structures (termed ‘pores’ throughout the remainder of the text) in the internal membranes of the planctomycete bacterium *Gemmata obscuriglobus* can be observed in transmission electron microscopy images of thin or thick sections of whole cells (Fig 1A, 1B, S1 Fig). When thin sections of cells prepared either via high-pressure freezing or cryosubstitution without high-pressure are examined, favourable planes of section reveal pore-like structures along sectioned nuclear envelopes. In Fig 1B a circular complex (arrowhead) appears in a gap between folded regions of the nuclear envelope membranes (arrows). Appearance of the pore is comparable to that of sectioned eukaryote yeast pores prepared by high-pressure freezing [22]. The diameter *ca.* 35 nm of the circular complex in Fig 1B is consistent with the diameter of pore structures in negatively-stained preparations (see below). One ring of the pore complex analogous to that of the nuclear and cytoplasmic rings of the eukaryote nuclear pore may be visible *en face* by a favourable tangential plane of section (Fig 1B). The position of such a pore within the membrane in a whole cell can be seen in S1A Fig, and the details of ring and central plug and spoke-like structures within such pores seen in face are also evident in S1B Fig.

**Fig. 1.**
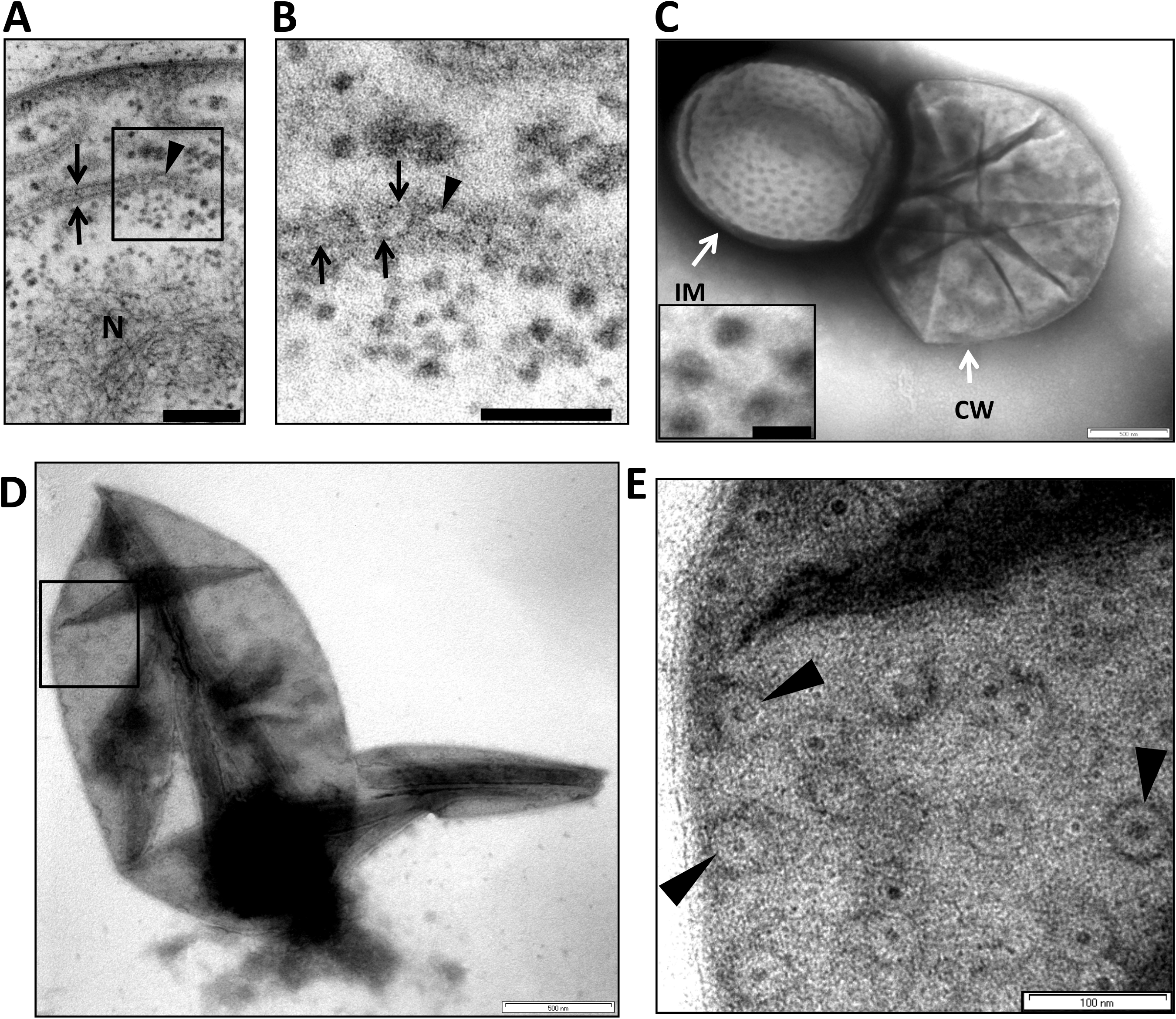
Pores are inserted into the internal membranes of *Gemmata obscuriglobus* cells. (***A***) Transmission electron micrograph of a thin-section of a cryosubstituted cell of *G. obscuriglobus*, showing a portion of the nuclear body envelope, apparently consisting of two closely apposed membranes enclosing the fibrillar nucleoid DNA (N) (for evidence of DNA fibrillar nature in *G. obscuriglobus* see [9]. The membranes (arrows) are interrupted by a disc-like structure (indicated by arrowhead within the boxed region) consistent with a pore complex inserted between the membranes on either side. Bar, 50 nm. **(*B*)** Enlargement of the sectioned cell of *G. obscuriglobus* seen in Fig 1A, showing a disc structure (arrowhead) seen *en face*, situated between the folded double membranes of the nuclear body envelope on either side (arrows). Bar, 50 nm. **(*C*)** Transmission electron micrograph of cell lysed by grinding in liquid N_2_, followed by negative staining of thawed cells with uranyl acetate. An internal membrane fragment (IM) possibly representing the nuclear body envelope or other internal compartment membranes appears to have been released from a lysed cell, and the mostly intact cell wall (CW) can also be seen. The membrane displays numerous evenly distributed pore structures on its surface, enlarged views of which can be seen in the inset. Bar, 500 nm. Inset shows enlargement of pore structures, which display a dense core surrounded by a light ring further surrounded by a dense ring. Bar, 50 nm. (***D***) Transmission electron micrograph of negatively stained preparation of a completely released internal compartment from cells lysed as in C. Pore structures are widely distributed over the membrane surface including within the boxed region. The ‘canoe’ shape is typical for pore-containing membranes. Bar, 500 nm. **(*E*)** An enlarged view of the boxed region in Fig 1D showing the large pore structures (arrows), each displaying dark pore centre regions, and lighter inner and outer ring structures, distributed densely on the membrane surface. Bar, 100 nm.

When cells of *G. obscuriglobus* are gently lysed, internal membrane can be released through the broken cell walls (Fig 1C), the overall shape of which is consistent with a sphere or compressed sphere (Fig 1C and 1D). The released membranes are covered with circular ring-like structures, possessing a dense circular centre (Fig 1E).

We have also demonstrated the presence of pores on the internal membranes of intact *G. obscuriglobus* cells using the freeze-fracture technique (Fig 2A and 2B). In freeze-fracture replicas, it is possible to unambiguously identify the membrane surfaces displaying pores on a single major internal membrane-bounded organelle, since the position of this membrane-bounded structure is clearly visible within the cross-fractured cells. The pores, circular structures, appear on one outer surface or fracture surface of the organelle envelope, which clearly overlies another fractured membrane of this envelope, indicated by the splits visible in the envelope. The complexity of the organelle membranes has been observed in *Gemmata* in tomographic and thin-sectioning studies [7,8,10]. In enlarged view (Fig 2A insets, Fig 2C) each pore represents a circular complex in which an outer ring surrounds a centre which usually appears darker than the outer ring to a degree depending on the metal shadow angle relative to the pore aspect. The size of pores in the freeze-fracture replica in Fig 2C are *ca*. 32 – 45 nm, which is consistent with dimensions obtained from negative staining but such dimensions would be expected to be more variable due to metal shadow used to prepare the replicas. It is of interest that we obtained similar freeze-fracture replica images of pore-like structures with central core and ring structure on internal membranes in another *Gemmata* strain, CJuql4 (ACM5157) (S2 Fig), closely related to the general *Gemmata* cluster phylogenetically, and a member of the same roughly genus-level phylogenetic group with high bootstrap support in a phylogenetic tree of planctomycetes [23]. These pore-like structures have a central core and ring structure and are 23-38 nm in diameter. Cells of this additional *Gemmata* group strain have been shown to possess an internal membrane-bounded nucleoid-containing compartment in thin sections, similar to that of *Gemmata obscuriglobus* [23]. These results therefore reinforce the conclusions from *Gemmata obscuriglobus* regarding the significance of pores as structures associated with the *Gemmata* group-characteristic internal membrane envelopes often surrounding the nucleoid region. A membrane-bounded region surrounding the nucleoid represents a type of structure known only from the *Gemmata* group strains so far among either planctomycetes or among any other domain Bacteria.

**Fig. 2.**
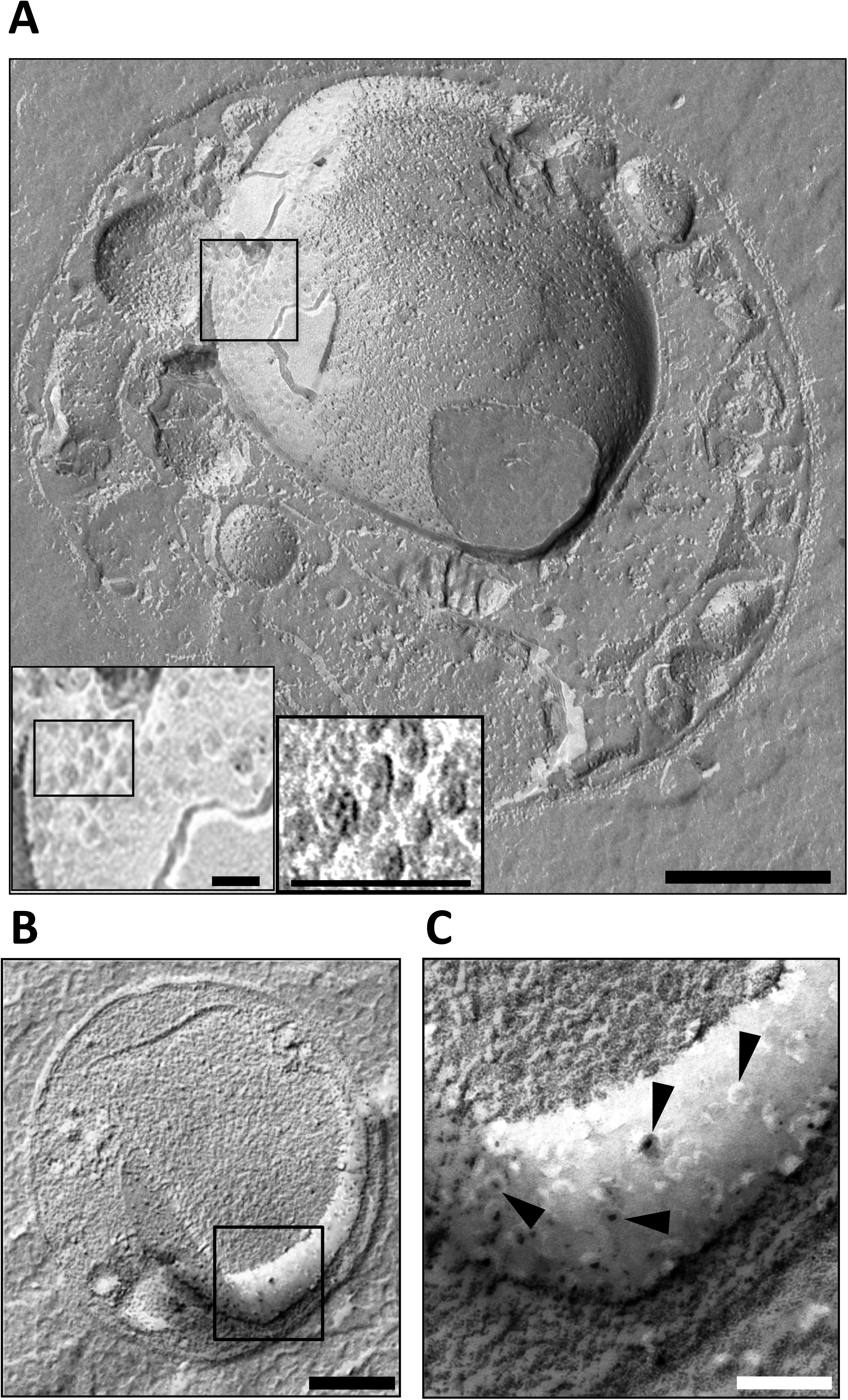
*Gemmata obscuriglobus* internal membrane pores as seen in freeze-fractured cells. **(*A*)** Transmission electron micrograph of a platinum/carbon (Pt/C)-shadowed replica of a whole cell of *G. obscuriglobus* which has been prepared via the freeze-fracture technique. Bar, 100 nm. Inside the cell, a large spherical internal organelle consistent with the nuclear body organelle surrounding the nucleoid has been fractured (split) along and through the surface membranes of its envelope. Pores with a central core and at least one surrounding ring are visible on one region of one of the membranes of this organelle surface. Insets represent successive enlarged views of the boxed region in the main image displaying the pores at higher magnification. Bars, 100nm. At the highest enlargement the substructure of each of several pores can be resolved including central core and surrounding inner dark and outer light rings (right inset). (***B***) This micrograph of the whole cell reveals an apparently cross-fractured major internal organelle compartment and a membrane surface (boxed) representing a fracture through the membrane surrounding the organelle. Bar, 200 nm. **(*C*)** An enlarged view of the boxed region of the freeze-fractured cell seen in Fig 2B showing a region of a membrane surface where roughly circular pore structures (arrowheads) are visible, in some cases with two light rings surrounding a dark centre,. Bar, 50 nm.

In negatively-stained membrane fragments released from cells lysed by sonication, the pores consist of a thin inner ring immediately surrounding the electron-dense pore centre, and an outer thicker ring distinguishable from the inner ring (Fig 3A). Assuming uniform distribution, there are ca. 87 such pores per µm^2^ of such a membrane sheet. At higher magnification the pores of *G. obscuriglobus* appear to be composed of subunits with internal “plug” (Fig 3B and 3C). In membrane fragments released by mechanical lysis and negatively stained with uranyl acetate, the outer diameter of the pore based on the outer ring diameter was calculated as 33.5 ± 2 nm, the diameter of the inner ring as 17.5 ± 2 nm, and that of the electron-dense pore centre (equivalent to an ‘inner pore’ or ‘central plug’) as 9.5 ± 0.8 nm (S3 Fig).

**Fig 3.**
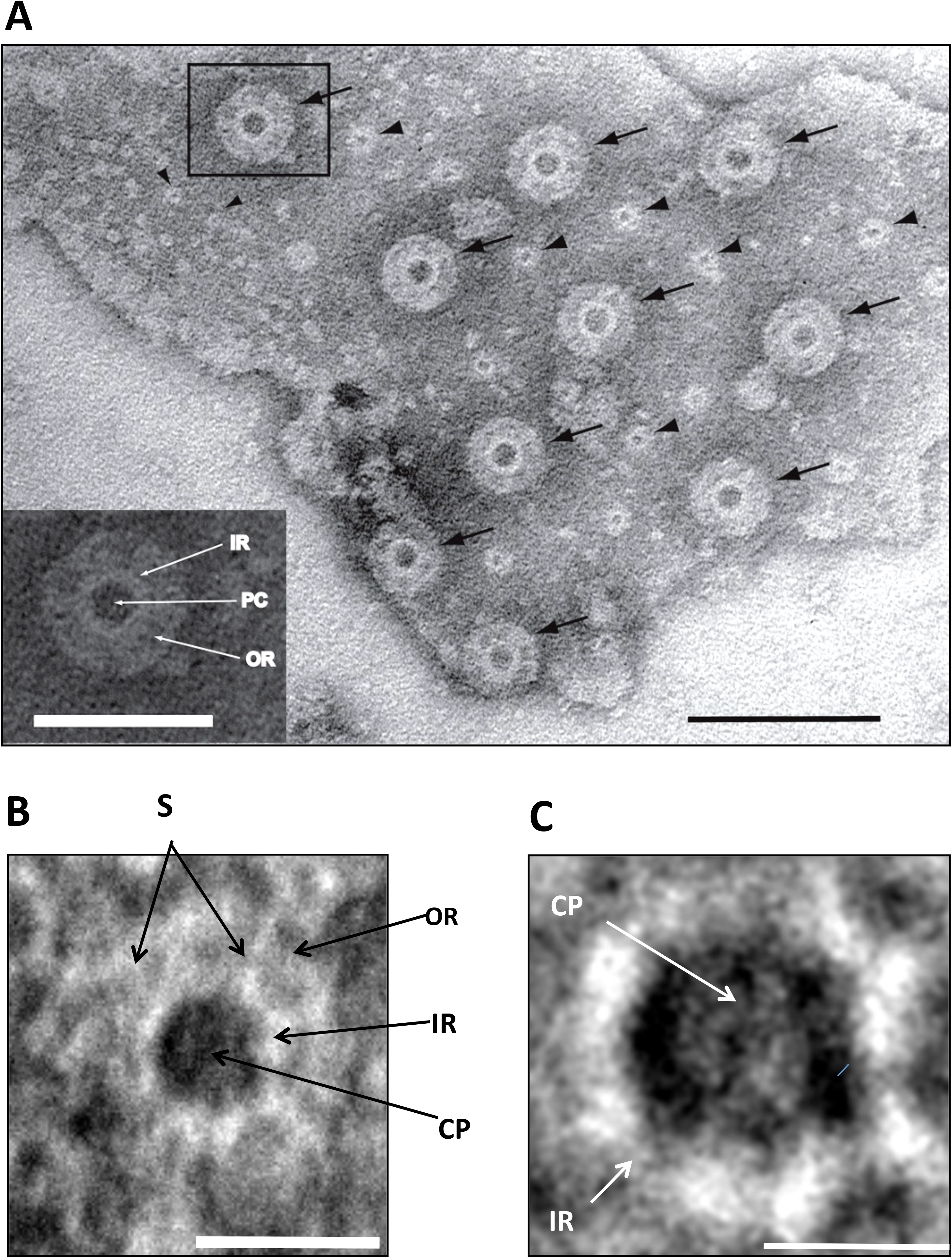
Pores in the membranes of *Gemmata obscuriglobus* released via sonication. **(*A***) Transmission electron micrograph of a membrane fragment released from a lysed cell via sonication and negatively stained with ammonium molybdate. Large pores (arrows) with relatively electron-dense pore centers surrounded by a thin lighter inner ring and a thicker outer ring are seen. Smaller pore structures (arrowheads) are also visible and may represent either another class of pores or a result of a reverse view of the same large pores resulting from overlapping folds in the membrane (evidence for such structures is not derived from other microscopy methods). Bar, 100 nm. **Inset:** enlargement of boxed large pore in main Fig where a pore centre (PC), an inner ring (IR) and an outer ring (OR) can be distinguished. Bar, 50 nm. (***B***) TEM of a pore seen in negatively stained membrane fraction isolated from sonication-lysed cells, showing pore complex structure including outer ring (OR), inner ring (IR), spokes connecting inner and outer rings (S) and central plug (CP). Bar, 30 nm. **(*C*)** Enlarged view of the inner ring (IR) and central plug (CP) of the boxed pores in Fig 3A, the octagonal shape of the rings (especially visible if the outer edge of the outer ring is traced) is consistent with an eight-fold symmetry. Bar, 15 nm.

After treating of the membranes released from lysed cells and isolated (as ‘fraction 3’, see below) via density gradient fractionation with detergent, aggregates of individual pores could be detected (S4 Fig) with only degraded membrane between them. Individual pores in such preparations show a central dense core surrounded by a light ring and in some cases material projecting from the outer rim of the ring possibly representing spokes normally connecting inner to outer ring in intact pore complexes.

In addition to these pores, two smaller classes of pores are also found. They may be seen most clearly in membrane fragments released via sonication (Fig 3A). The larger of these two classes consists of rings that are 14.5 ± 2 nm in diameter with an inner dense centre of 5 ± 1 nm wide, while the smaller class consists of pores 6± 0.9 nm wide and which also possess a dense centre and can appear in clusters. Neither of these smaller pore types seem comparable in structure to eukaryote nuclear pore structures in the sense of possession of both inner and outer rings as well as an inner dense centre. It is also possible that at least the 14.5 nm diameter class is the result of a reverse view (‘basket’ side view) of only part of the larger pore structures seen because of a folded membrane, since they appear most clearly in examinations of negatively stained membrane sheets rather than via other EM preparation methods. If true additional pore types, they suggest either specialization in pore function in these membranes or some stages in assembly for the largest pores. It is the largest pore type we have seen which is in any case most relevant to the argument of this paper since it is the class displaying most complex structure and is the class comparable with that of eukaryotic nuclear pores.

Thus, we have determined here from studies of whole cells and membranes released from lysed whole cells that at least one type of internal membrane contains pores. This has been demonstrated by three distinct electron microscope preparative techniques – thin-sectioning and freeze-fracture replica technique of the whole cryosubstituted cells, and negative staining of membranes released from lysed cells. If pores are genuine structures, one would predict that several different methods should be consistent e.g although negatively stained internal membranes released from lysis display the clearest examples of pores, methods such as freeze-fracture and thin-sectioning should also reveal entities with comparable structure on internal membranes in whole cells, and this is indeed the case. If such structures are significant functionally, one would expect they would be associated with a specific type of internal membrane and be able to find differences in protein composition of that distinct type of internal membrane.

To confirm the association of pores with a specific type of internal membrane and to investigate the composition of such membranes further we fractionated the internal membranes and applied proteomics and EM methods including immunoelectron microscopy to localize a specific protein of potential relevance to pore structure.

### There are three types of membranes in *G.obscuriglobus* cells, one of which contains pores

To purify membranes for structural and proteome studies we applied a two-step density gradient fractionation technique (S5 Fig). The discontinuous and subsequent continuous gradient fractionations aimed to separate the membranes at the highest possible level. This procedure resulted in appearance of three distinct membrane types (S6 – S8 Fig). Only one of the fractions (fraction 3, consisting of characteristic ‘canoes’), displayed pores on the membrane surfaces when examined via TEM after negative staining (S7 Fig). The large pores are present at high density (ca. 200 / µm^2^ of membrane sheet) and similar in size and structure to the complex pores seen in unfractionated membrane released from lysed cells (e.g. Fig. 3A). We were able to apply clear markers for two of these fractions via an antibody against a beta-propeller-containing protein (a protein exclusive to fraction 3, see below and S9A Fig), and via a specific antibody against *G. obscuriglobus* clathrin-like membrane coat (MC) protein gp4978 (shown in a previous study to react specifically with membrane vesicles associated with endocytosis-like protein uptake in *G. obscuriglobus* [24]. The anti-beta-propeller-containing protein reacted only with fraction 3 but not with fractions 2 or 6, and the anti-MC protein antibody demonstrated reactivity only against fraction 2 (S9B Fig).

The gradient fractionation technique demonstrates that membranes related to distinct internal structures of *G.obscuriglobus* can be separated, and the probing with antibodies clearly shows that the fractions do not contain significant amount of cross-contamination.

### Structural analyses of the pores reveal their similarity to the nuclear pores of eukaryotes

Electron tomography of the fraction 3 membranes shows that, topologically, these pores are membrane insertions (Fig 4A and 4B, S1 Video and S2 Video). By analysing digital slices through the tomogram, we see continuous membrane envelope followed by a pore structure and then followed again by continuous membrane (Fig 4B). Such pores display a projection on one side and in some slices the plug could be seen (Fig 4A and 4B). Cryo-EM analyses (S10 Fig) and a modified Markham rotation analysis of the electron micrographs from frozen-hydrated pore-containing membranes (Fig 4C and 4D) suggests an eight-fold symmetry for the organization of these pores, indicating that these complexes are likely to be modular constructions. It should be noted that we have used the same technique here as applied in the first demonstration of eight-fold symmetry in nuclear pores from invertebrate and vertebrate animal species[25]. Eight-fold rotation gave strongest reinforcement e.g. relative to 7-fold rotation. Markham rotation cannot by itself prove 8-fold symmetry of pore structure, but taking both the Markham analysis and the clear radial and octagonal symmetry visible in the original micrographs into account, on current evidence, 8-fold symmetry is the most likely possibility.

**Fig 4.**
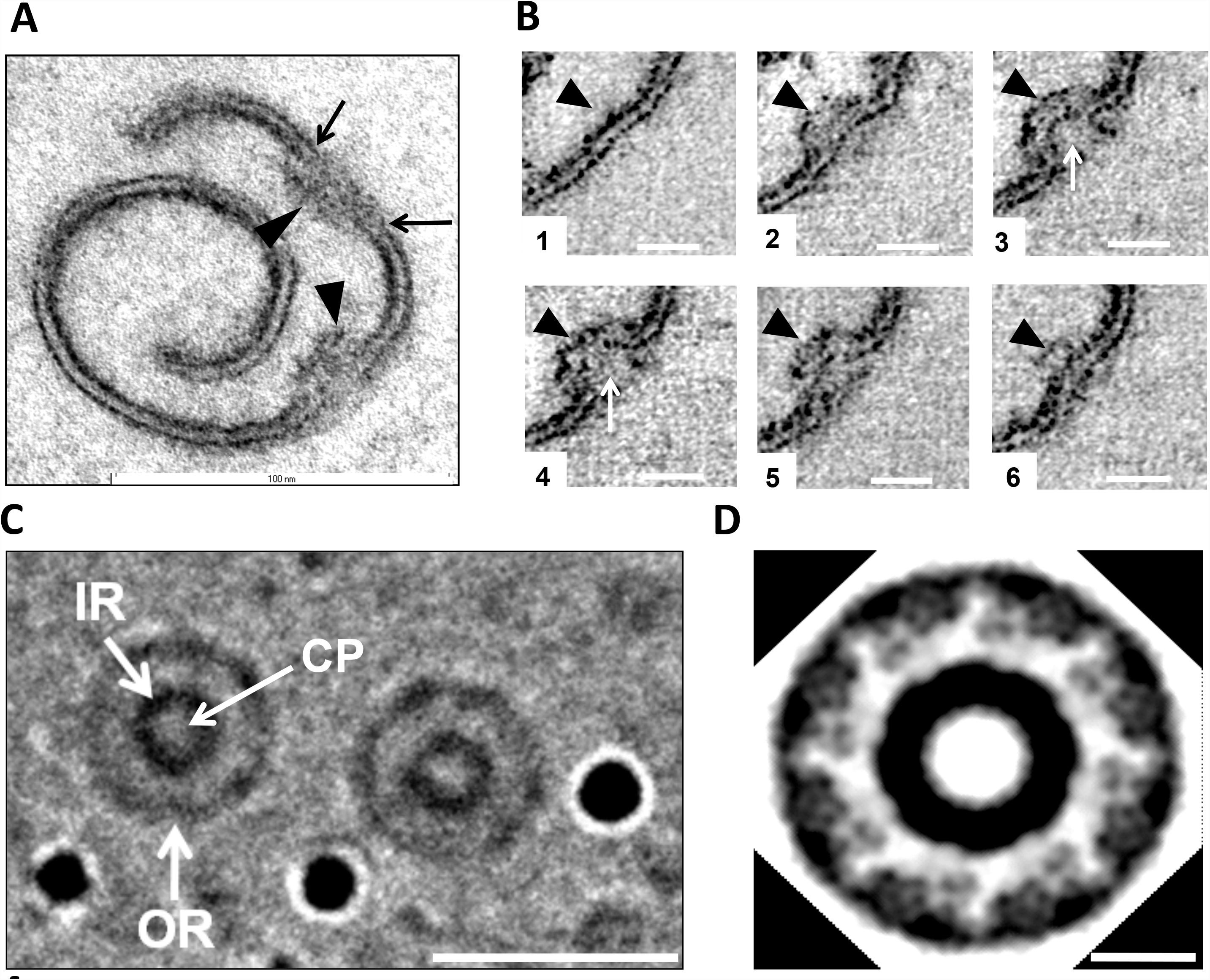
Architecture of the *Gemmata obscuriglobus* pore. (***A***) Structure of pores embedded into membranes from fraction 3 purified via density gradient centrifugations and visualized via TEM of thin-sections. The spiral seen consists of a membrane (arrows) in which pores are embedded interrupting dense-light-dense layers of the trilaminar membrane. Basket structures (arrowheads) of each pore complex project only from one side of the membrane. The inner and outer dense leaflets of the membrane are seen to be connected forming a continuous folded membrane (on each side of the pore) of extreme membrane curvature (arrows). Bar, 100 nm.(See also S1 Video and S2 Video for 3D reconstructions of the membranes and S3 Video for 3D reconstruction of the pore). **(*B*)** Transmission electron micrographs from a tilt-series of one pore. In panel 1 intact membrane without pore is seen, while in panels 2-6 passing through progressive slices generated via the tilt-series, the pore appears, interrupting the trilaminar membrane on either side, and most clearly indicated by a basket structure projecting below the plane of the membrane (arrowhead). In panels 3 and 4, the central plug region of the pore can be seen (arrows). In panel 6 the trilaminar membrane is again continuous, but some parts of the basket structure are still visible (arrow). This series is consistent with the interruption of membrane by embedded pore structures, the basket component of which projects beyond the membrane plane. Bar, 20 nm. (***C***) Micrograph from cryo-EMof a frozen-hydrated preparation of the isolated and sucrose-purified fraction 3 membranes. Two randomly selected pores (see Fig S10 supplement 1for cryo-EM of fraction 3 membrane sheets from which these pores were selected) clearly display inner ring (IR), outer ring (OR) and central plug (CP).This image has been processed via uniform application of a conservative bandpass filter (respective low-and high-frequency cut-offs of 40 and 3 pixels). Bar, 30 nm. (***D***) Modified Markham rotation analysis of one of the pores from Figure 4C showing reinforcement of 8-fold symmetry of pore structure. Bar, 10 nm.

3-D reconstructions of the pores inserted into the membranes confirms a basket-like structure, with struts connecting the pore to a distal ring, and projecting from the part of the pore inserted in membrane (Fig 5 and S3 Video).

**Fig 5.**
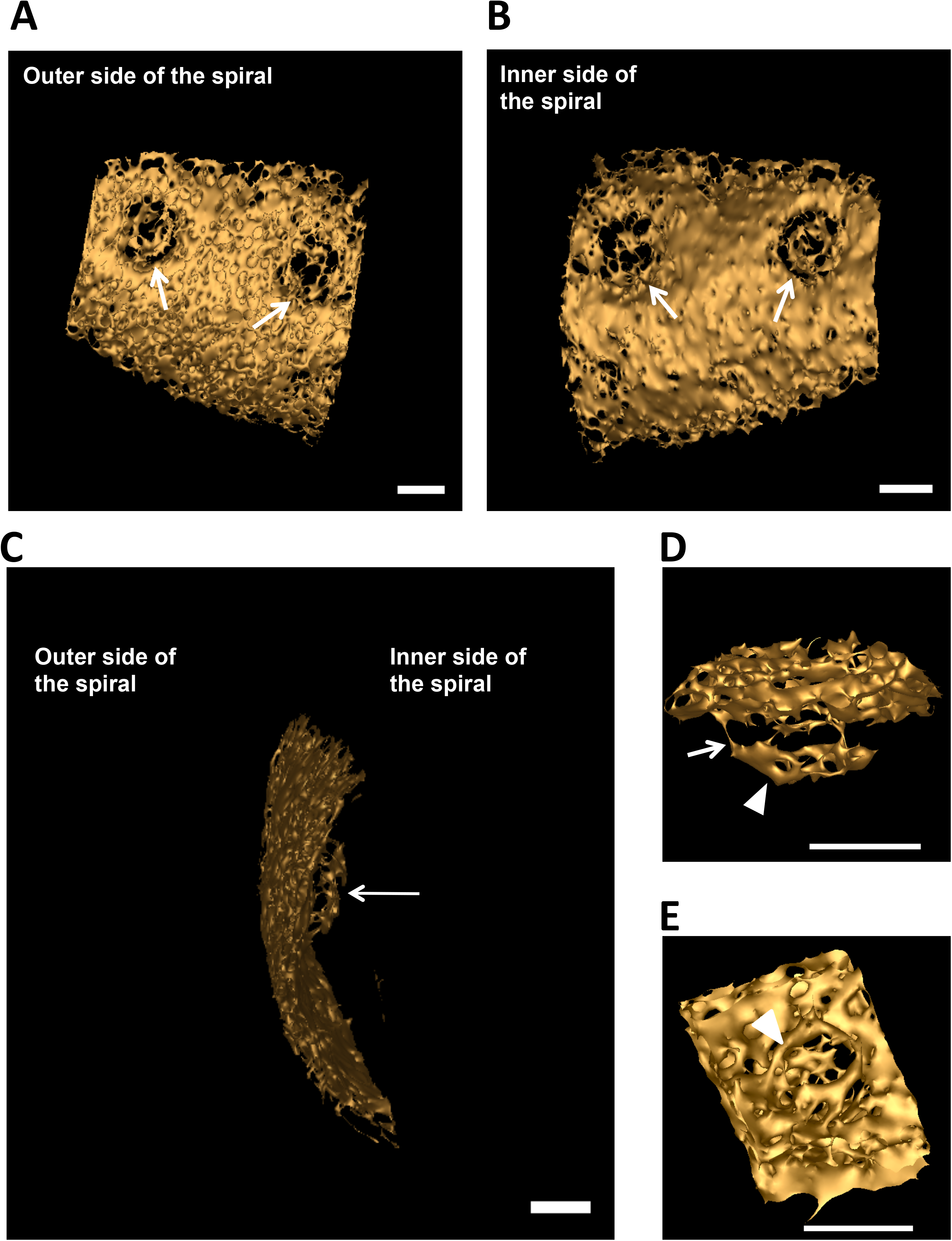
3-D reconstructions of the pore complex. **(*A* and *B*)** Views of the 3-D reconstructions based on one spiral membrane from fraction 3 membranes (see Fig 4A). Pore complexes (arrows) are visible as embedded structures in the surface of the envelope, shown as viewed from the inner side of the spiral in Fig **5A** and from the outer side in Fig **5B**. Fig **5C** shows the basket structure of one of these pores projecting from the inner side of the membrane spiral. Bars, 20 nm. **(*D* and *E*)** Reconstruction of architecture of a single pore seen from two different angles. In panel ***D***, a side view of the pore displays the basket structure with its distal ring (arrowhead) and a series of struts (arrow) connecting with the main pore rings. In panel ***E***, a top view shows the ring-like element (arrowhead) of the main part of the pore and a central plug structure is visible within the pore connected to the ring’s inner rim via spokes.

In summary, the pores embedded in the internal membranes of *G.obscuriglobus* display startling similarities to the nuclear pore complex of eukaryotes [26,27,28,29]. For structural comparison of the *Gemmata* pore with a eukaryotic pore model [2] see S11 Fig. The pores are comparable to the appearance and distribution of nuclear pores on negatively stained isolated nuclear membranes of yeast, plant and mammalian liver cells [30,31,32]. Those similarities include apparent eight-fold symmetry of its components, the presence of a basket structure projecting asymmetrically from the membrane plane, and the presence of at least two rings within that plane. These structurally complex pores of *G.obscuriglobus* are considerably smaller than nuclear pores on isolated nuclear membranes of eukaryotic cells. At *ca.* 35 nm in frozen-hydrated membranes, they are only approximately a third the diameter of characterized nuclear pores [28]. The frozen-hydrated NPCs of yeast are 96 nm wide [33], the *Xenopus* or yeast nuclear pore complexes in thin-sectioned cells are *ca.* 120 nm and 103 nm wide respectively [22,34], negatively stained NPCs of *Xenopus* are 107 nm wide [35], and detergent-released negatively stained are 133 nm wide [36], and finally intact *Dictyostelium* slime mould NPCs studied by cryo-electron tomography are 125 nm wide [26]. However, despite the difference in size of the *Gemmata* pores compared to those of eukaryote nuclear envelopes, there are clear analogies in structure, since the eukaryote pores also have a central plug and a central ring-like assembly composed of spokes sandwiched between a cytoplasmic ring and nuclear ring[22], and in negatively stained detergent-treated nuclear envelopes of *Xenopus* oocytes pores are visible as rings containing a central plug connected to the ring by spokes [27].

Our structural studies of the *Gemmata* pores are summarised in a deduced model (Fig 6). From Cryo-EM (Fig 4C and 4D) the pores from the “basket side” have two rings of ca 20 nm (inner ring) and 35 nm (outer ring) in diameter. The pores from this side display an octagonal symmetry, consistent with rotational folding analysis of cryo-electron micrographs from frozen-hydrated TEM of pore-containing membranes (Fig 4C and 4D). Within the central core region is a central ‘plug’ of *ca.* 10 nm in diameter (Fig 4C and 4D). At the higher magnification the central plug is seen as a structure connected to the inner side of the pore (Fig 3C). From the opposite side the pores also have two rings. As from transmission electron micrographs of the whole compartments lysed by grinding in liquid N_2_ (Fig 1C and 1E), Pt/C-shadowed replicas of whole cells (Fig 2C), and 3D reconstructions of the pores (Fig 5A and 5E) the outer ring is *ca.* 25 nm and the inner ring is *ca.* 20 nm in diameter. As calculated from the isolated membranes used for tomography studies (Fig 4B) the maximum distance from the top side of the pore to the end of the basket is 25 nm.

**Fig 6.**
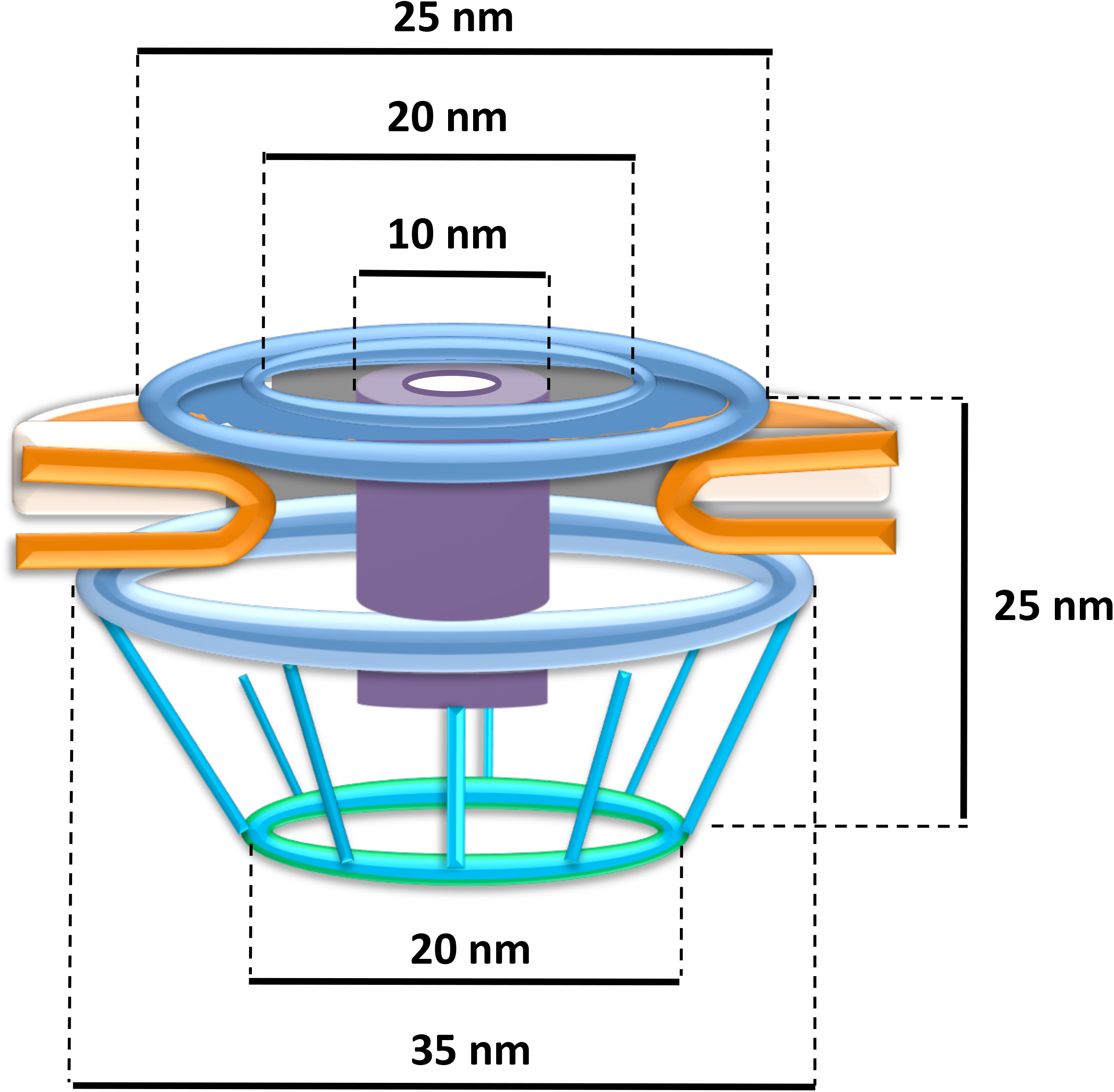
Model of the pore complex of *Gemmata obscuriglobus*. The pore complex is composed of at least two concentric upper rings (blue), and a lower ring (light blue) connected by struts to a distal ring (green) to form a basket structure. The central plug (purple) rests within the inner ring and spans the length of the pore. The whole pore complex rests within membrane (orange). The structure and dimensions are based on available data from all EM methods applied, from both whole cells and fraction 3 isolated membranes, and with minimal extrapolation, so that although the pore is probably not a hollow structure the space within the pore has not been filled in.

### Whole membrane proteome analyses reveal distinct protein content for three types of internal membranes and the cell wall

Proteomics was applied to cross-compare proteins from the fractions obtained via gradient centrifugations (see above) in order to distinguish proteins unique to the pore-containing membranes. Protein composition appeared different for pore-containing membranes (fraction 3) compared to those with no visible pores (fractions 2 and 6) (Fig 7A). 342 membrane and membrane-associated proteins were identified from all fractions examined (Fig 7B, S1 Table), 46% of which are annotated as hypothetical proteins. In non-pore-containing membranes (fractions 2 and 6), many constituents of the respiratory chain, ABC transporters and secretion system components were identified. The recently described MC-like vesicle-associated protein [37] was found exclusively as a constituent of fraction 2 (S1 Table). Fraction 2 appears enriched with vesicles, which are present at high number within the paryphoplasm. Some vesicles may be formed during endocytosis and are derived from the cytoplasmic membrane [24,38]. Proteins such as glycosyl transferases, a number of dehydrogenases, including the NADH-dependent dehydrogenase, and the periplasmic solute-binding protein, were restricted to fraction 6, which suggests that this fraction is enriched for the cytoplasmic membranes, and the other two fractions do not contain detectable amount of cytoplasmic membrane debris. ATP synthase was found *only* in this cytoplasmic membrane fraction (fraction 6) and in the vesicle-enriched membrane fraction (fraction 2), and *not* in the pore-containing membrane fraction (fraction 3). This indicates that the function of the pore-containing membranes is probably not that of electron transport or energy generation. In the pore-containing membranes (fraction 3), we identified 128 proteins, 39 (30.5%) of which were unique to this fraction (S2 Table). Common proteins found by proteomics analysis in all three fractions (S1 Table) are predicted pilins and predicted ribosomal proteins, which we believe are the result of contamination since actual pili structures were detected in small number by electron microscopy in all three fractions, and heavy ribosomal subunits which might be distributed along the entire gradient. In our proteomic analysis regarding significant proteins correlated with presence of pore structures, we only consider proteins exclusive to fraction 3.

**Fig 7.**
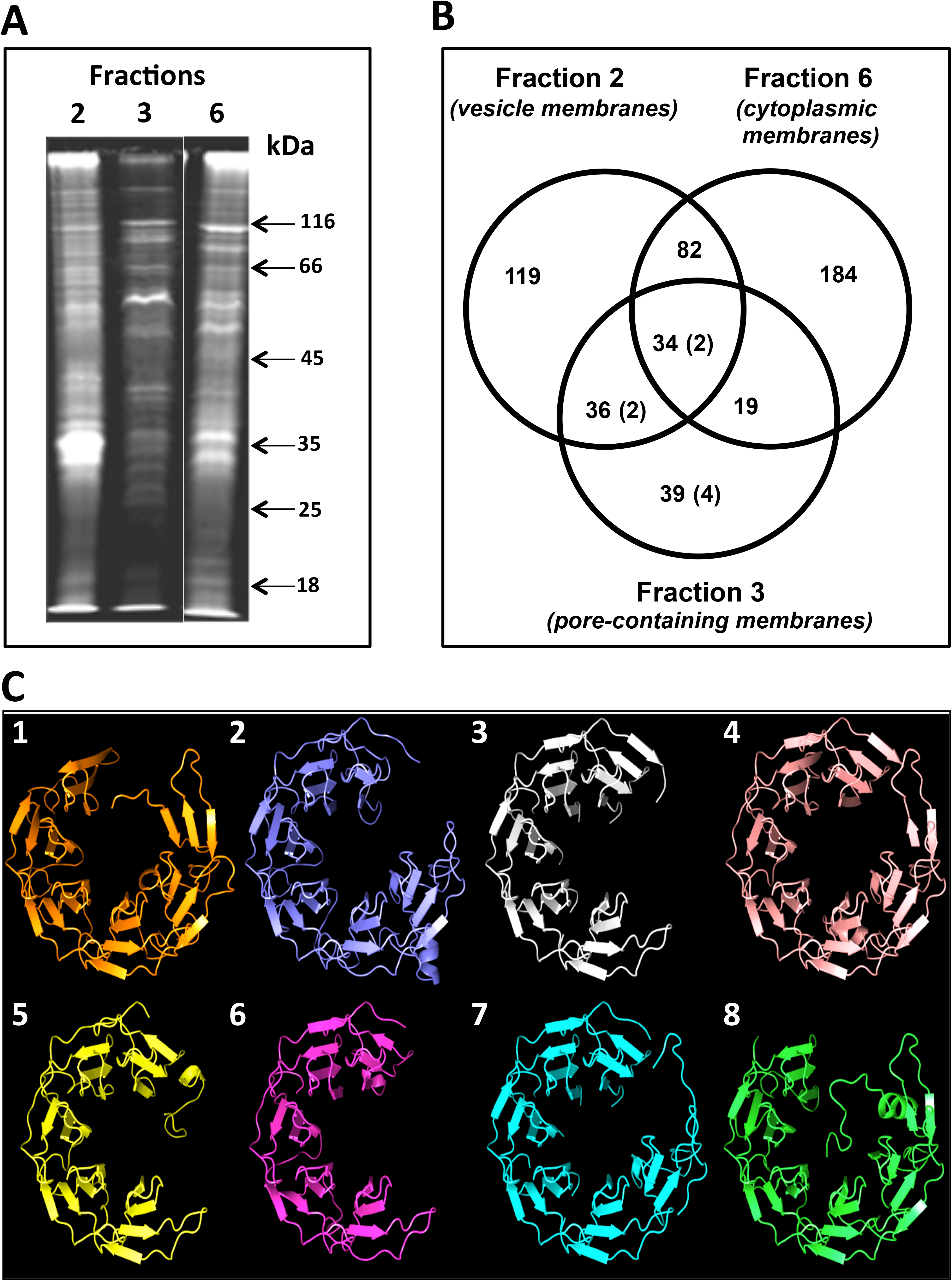
Protein composition of *Gemmata obscuriglobus* pore-containing membrane. **(*A*)**SDS-PAGE gel showing that *G.obscuriglobus* cells have three different types of membranes. Exclusively pore-containing membranes (fraction 3) display a characteristic protein profile distinct from that of membrane fractions which do not possess pore structures. **(*B*)** Venn diagram showing the number and distribution of proteins among the fractions and (in brackets) the number of proteins with the beta-propeller folds. The members of the beta-propeller cluster belong either exclusively to fraction 3 (4 proteins), or to fractions 3 and 2 (2 proteins), and to fractions 2, 3 and 6 (2 proteins). No beta-propeller containing proteins were found exclusively in fractions 2 or 6. **(*C*)** A beta-propeller family found in fraction 3 (pore-containing membranes), including some exclusive to fraction 3. Cluster analyses revealed a set of proteins with conserved C-terminal regions (Figs S13 – S16) that model beta-propeller folds with high (>95%) confidence. Models 3 (for protein ZP_02737072), 4 (ZP_02736670), 5 (ZP_02734776) and 6 (ZP_ZP_02734577) were deduced from proteins found exclusively in fraction 3 (pore-containing fraction); models 2 (for ZP_02737073) and 7 (for ZP_02733245) were deduced from proteins found in fractions 3 and 2 only; models 1 (for ZP_02737797) and 8 (for ZP_02731113) – for proteins found in fractions 3, 2, and 6 (Table S4).

It should be noted that the pores described here structurally resemble crateriform structures, a characteristic signature structure of planctomycete surfaces [39] and isolated cell walls[40] including those of *G. obscuriglobus*[41]. These crateriform structures could be seen on the surface of whole cells as circular regions and in the cell slices used for TEM as pits protruding the cell wall and cytoplasmic membrane, confirming published data on *G. obscuriglobus* [42]. It might be proposed that the pores described here represent in fact the crateriform structures as seen in purified cell walls (Fig S12). However, these crateriform structures when negatively stained display a dense centre but only one surrounding ring, so apparently differ from the more complex internal membrane pores. The membrane fractions we used as starting material for the gradient fractionations should not have contained significant amounts of wall, as they are effectively eliminated during preparations of the membranes for initiating the gradient fractionations; the walls are denser than any of the membranes and are pelleted and thus separated at relatively low centrifugation speed. Any walls remaining after this step are pelleted during fractionation in any case. The walls also cannot be lysed in the buffers usually used for dissolving the membranes for Western blot analysis, so their proteins would not be detected in isolated gradient-fractionated membranes. Compared to the membranes, the walls are highly resistant to boiling in 10% SDS, and such boiling is used for purification of the walls from the membranes during wall isolation. In a separate experiment we have isolated the walls via boiling the *G.obscuriglobus* cells with 10% SDS and analysed their protein content. The cell walls revealed some proteins homologous to proteins which have in previous studies been shown to be characteristic of the wall of planctomycetes [43,44]. Via proteomics we have identified the major constituents of the walls, the so-called YTV-proteins (Gobs_U38067, GobsU_28375, and GobsU_21360). The bacterial cell wall marker peptidoglycan has now been reported to comprise at least part of the wall composition in *G. obscuriglobus* and some other planctomycetes [17,19], and lipopolysaccharide has been reported in *Gemmata obscuriglobus* [18] but proteins appear to comprise significant proportions of wall in *G. obscuriglobus* as well as other planctomycetes [40,41,45]. However, no planctomycete cell wall/surface protein homologs were found via proteomics in any of the three membrane fractions isolated from fractionation of lysed cells, reinforcing evidence from analysis of enzyme markers that all of these three fractions are from membranes other than cell wall/cell surface structures [42]. In addition, sectioned whole cells immuno-gold-labelled with a fraction 3 specific antibody (see below) were shown to label only internal membranes, and there was no labelling of walls or other surface structures. Thus we conclude that the pores in fraction 3 membranes are not wall/surface crateriform structures either on the criterion of location within the cell or on a criterion of protein composition. The pores represent structures embedded into internal membranes and do not represent crateriform structures or any other wall components. This implies potential performance by such internal membranes of functions such as transport of material between internal membrane-bounded cell compartments.

Thus, we have demonstrated here that the three membrane types separated by density gradient centrifugation are distinct from each other in their composition of specific proteins. Continuity of cytoplasmic membrane with internal *Gemmata* membranes has been proposed as part of the concept of the planctomycete cell plan as essentially one of a classical Gram-negative cell [11,46]. However, our data concerning protein composition of isolated membrane fractions does not support this concept, but rather is consistent with a concept that genuine internal compartmentalisation exists within the *Gemmata* planctomycete cell, where the compartments are separated from each other by different types of membranes. Our data is also consistent with distinction of cytoplaamic membrane from two different types of internal membranes.

### Bioinformatic analyses of the membrane proteome

We characterised the identified set of proteins using a range of bioinformatics-based analyses (S Text and S3 – S6 Tables). Initial blast analyses indicated that many of the proteins showed little or no similarity to proteins outside *G. obscuriglobus*. We therefore performed Blast cluster analysis (VisBLAST) to establish whether any proteins in the pore-containing membranes exhibit sequence similarities to one another. We also performed profile-based (phmmer) screens to search for more distant similarities, and ran structural predictions using Phyre2 [47] to assess similarity of proteins to known folds. Sequence and structural analyses of our membrane proteomics data revealed the presence of a number of bacterial transmembrane proteins, including outer membrane efflux proteins, translocons and porins (S Text), underscoring the bacterial nature of these membranes. However, none of these were unique to pore-containing fraction 3, so their origin as contaminants from cell structures or components other than membranes of fraction 3 is possible, and they cannot be implied as characterizing any specific fraction 3 membranes. In addition, published structural data are not consistent with any of these transmembrane or ‘outer membrane’ protein homologs forming pores with dimensions and attributes similar to the pores we observe in *G.obscuriglobus*.

Cluster analyses performed on all proteins identified through proteomics revealed two groups of proteins from fraction 3 with substantial sequence similarity (S13 and S14 Fig). Phyre2 models generated for one of these clusters identified a conserved C-terminal beta-propeller fold for 8 of the 11 proteins making up this cluster (Fig 7C; S15 – S18 Fig, S4 Table). Beta-propeller folds are found in protein constituents of eukaryotic nuclear pore complexes and coated vesicles, and it has been proposed that the eukaryotic nuclear pore and coatomer complexes evolved from a suite of membrane-curving proteins with common structural elements [48]. We therefore searched for evidence of other folds associated with eukaryotic nuclear pore complexes. Most notably, we identified two fraction 3 proteins (ZP_02735673 and ZP_02736511) that model well (>95% confidence, Phyre2) against clathrin adaptor core proteins, exhibiting an alpha-solenoid architecture (S Text, S18 Fig, S3 Table).

The presence of protein folds characteristic of the eukaryote nuclear pore complex [49] is intriguing in light of our deduced pore model, since in the eukaryote nuclear pore complex, beta-propeller-and alpha solenoid (stacked alpha-helices)-containing proteins act as scaffold proteins [50,51]. Beta-propeller proteins form vertices in a lattice-like model of the NPC and have special sequence-independent protein—protein interaction functions [52] while stacked alpha-helices of other scaffold nucleoporins are central to the lattice model interactions, forming edges of the NPC scaffold lattice [50] and are also structurally related to soluble proteins significant to nucleocytoplasmic transport through nuclear pores [53]. While it is remarkable that structural prediction yields folds known from the eukaryote nuclear pore complex, neither β-propeller folds nor alpha-solenoids are unique to the eukaryote nuclear pore complex or endomembrane system, and examples of both folds are known from both Bacteria and Archaea [54]. For all *G. obscuriglobus* proteins carrying folds also found in eukaryote nuclear pore proteins, we find no evidence of substantive sequence similarity with eukaryote counterparts. Evidence of such similarity might be expected if recent horizontal gene transfer from eukaryotes was their origin. We therefore conclude these genes are not the result of recent transfer from eukaryotes. Our data instead indicate these are genes of bacterial origin.

### Immuno-gold labelling confirms association of the pores with internal membranes

One of the proteins identified in fraction 3 exhibiting a β-propeller fold (ZP_02736670), was selected for antibody generation with the aim of using the antibody to immunolocalize the protein. This antibody (ab6670) showed high specificity as established by Western Blot, and reacted specifically with fraction 3, which comprises pore-containing membranes (S9B Fig). It was used for immunolocalization experiments to assess localization of the pore-containing membranes within the cells. On whole sectioned cells, gold particles were observed exclusively at membranes within the cell cytoplasm and internal to both cytoplasmic membrane and paryphoplasm (ribosome-less cytoplasm). The antibody labelled membranes comprising the nuclear body envelope and membranes associated with riboplasm (ribosome-containing cytoplasm) (Fig 8 and S19 Fig). Consistent with this result, the antibody also recognised pores in the purified membranes from fraction 3 (Fig 9 and S19 Fig) Membranes from other fractions, without such pores, were not labelled when tested. Such labelling is consistent with the presence of the ZP_02736670 β-propeller fold protein within pore complexes found via proteomics exclusively in fraction 3 (the origin of the protein data from which the peptide antibody was prepared).

**Fig 8.**
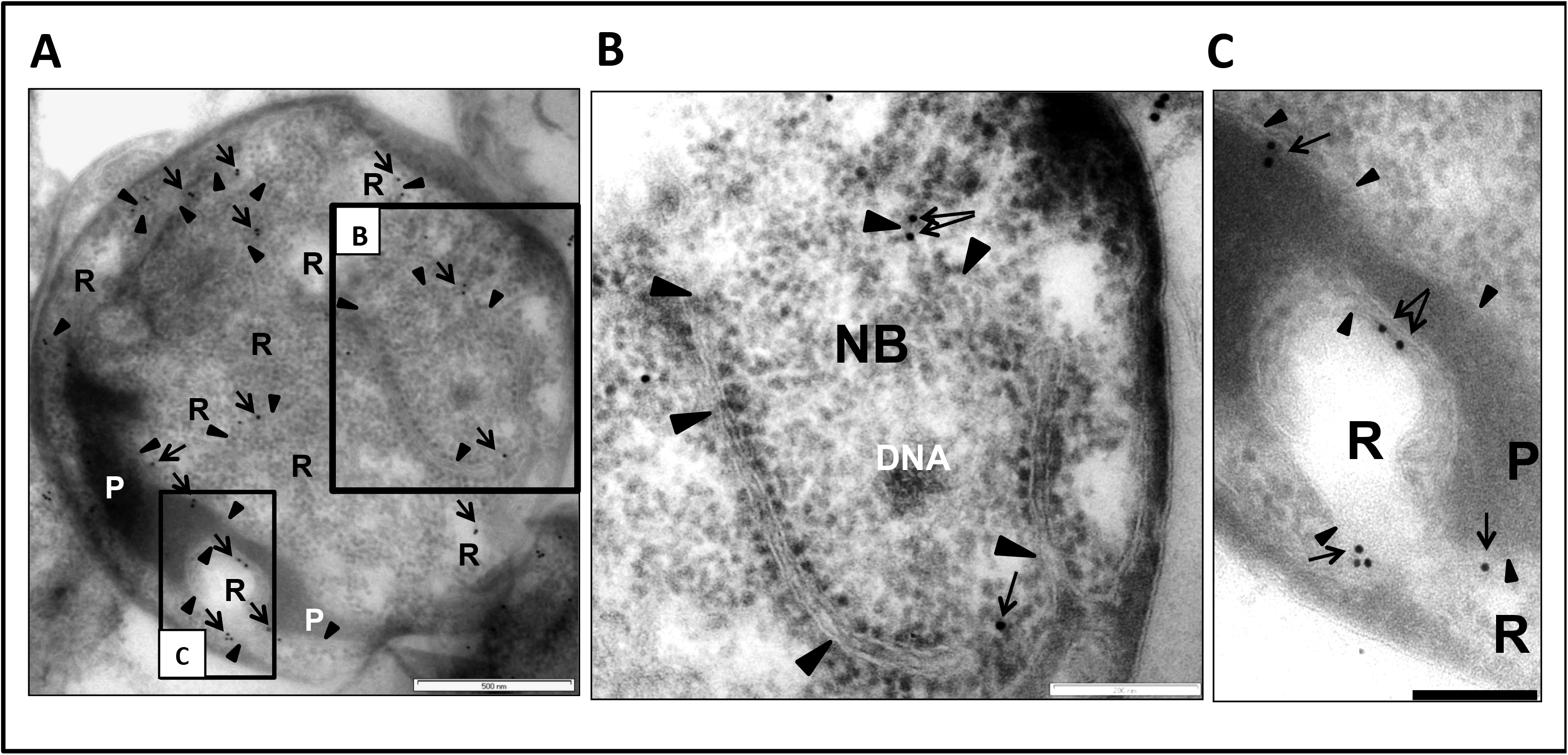
The antibody 6670 recognises internal membranes in *G.obscuriglobus* cells. **(*A*)** TEM of a thin-sectioned cryosubstituted cell labelled with the antibody 6670. The majority of the gold particles (arrows) are seen to be bound to intracytoplasmic membrane (ICM, arrowheads). This membrane separates the electron-dense ribosome-free paryphoplasm (P) from relatively electron-transparent riboplasm (R), as well as riboplasm vesicles from each other. A few particles label the border envelope between NB and riboplasm including double membrane regions. Bar, 500nm. **(*B*)** An enlarged view of the boxed region B in Fig 8A showing the nuclear body (NB) with nucleoid DNA. A few gold particles (arrows) are visible on the envelope membranes (arrowheads), separating NB and riboplasm. Bar, 200 nm. **(*C*)** An enlarged view of the boxed region C in Fig 8A showing an electron-transparent region continuous with riboplasm, surrounded by paryphoplasm which is separated from the riboplasm-continuous region by ICM. Gold particles (arrows) unambiguously label the ICM (arrowheads). Bar, 200 nm.

**Fig 9.**
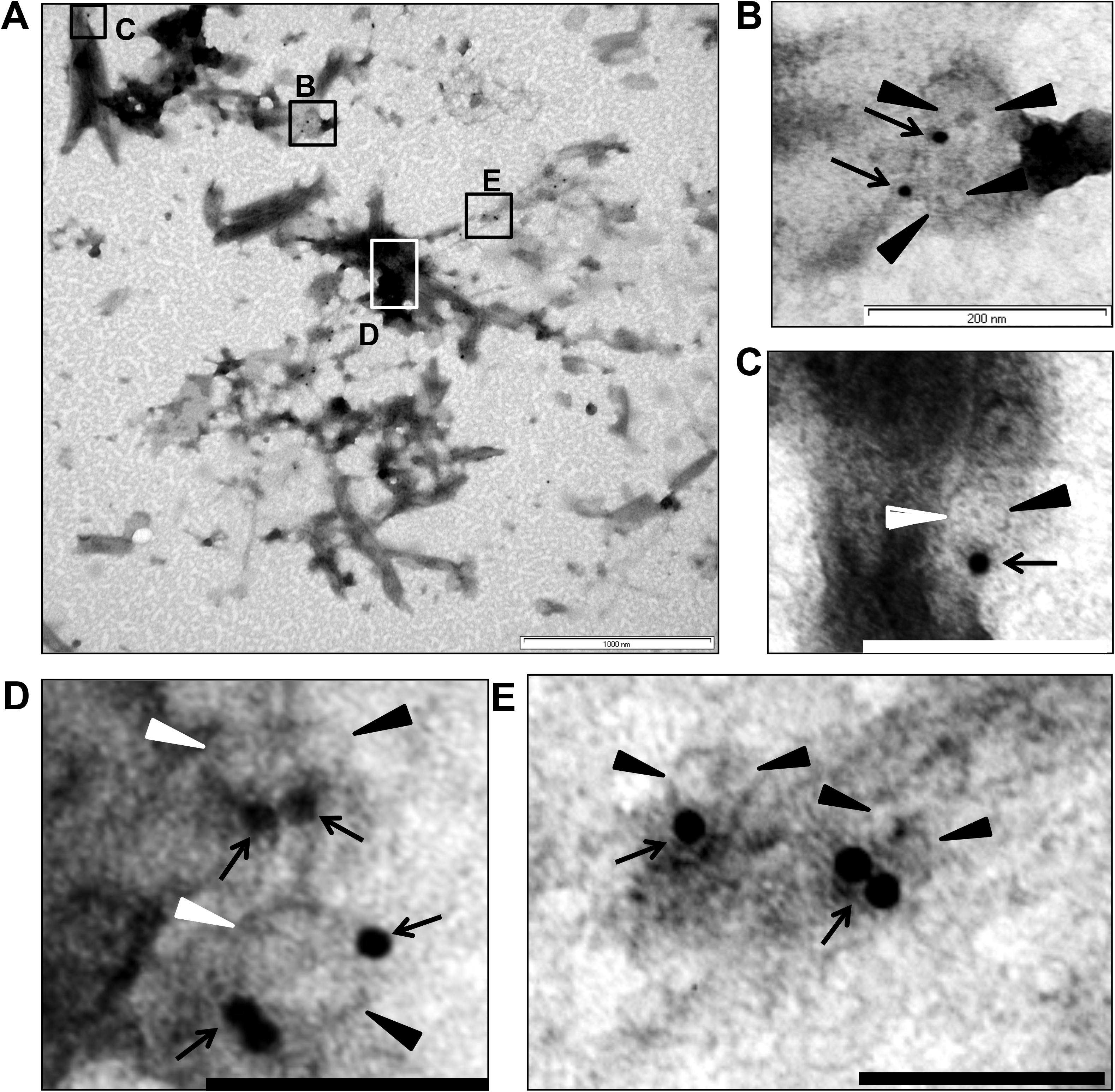
The antibody 6670 recognises pores in the isolated membranes. (***A***) Immuno-gold labelling of membrane sheets from membrane fraction 3 with the antibody 6670. In the majority of cases the gold particles indicating antibody can be seen as associated with the outer ring of the pores. Panels (***B**, **C**, **D***, and ***E***) show enlarged areas of **(*A*)**, which are marked as boxes in (***A***). In all the cases the gold particles can be seen at the edge of the pores. In some cases more than one gold particle is associated with the pores (for example see box **(*D*)**, the bottom pore which is surrounded by three particles). For statistical analyses approximately the same areas were used for counting the particles: 397 particles were observed as associated with pores (distance from a pore does not exceed 20nm) and 45 particles were considered as not associated. Arrows indicate gold particles, black or white arrowheads (depending on background) – pores. Bars, **A** – 1 µm, **B** −200nm, **C**, **D**, and **E** – 100nm.

### Evolutionary implications of the findings

Our results imply two alternative possibilities, both equally profound. One is that the architectural similarity of pores in *Gemmata* bacteria and eukaryotes may be the result of a shared deep evolutionary origin, so that they are homologous and related by vertical inheritance. This would be consistent with past analyses that have placed *Planctomycetes* among the deepest branching phyla within domain Bacteria [55], and more recent analyses support the view that other members of the PVC (Planctomycetes, Verrucomicrobia and Chlamydiae) superphylum also carry compartmentalized cell plans [56,57]. In our view, the lack of detectable sequence similarity precludes homology resultant from recent horizontal gene transfer from a eukaryotic source. More generally, the lack of an unambiguous signal for common descent, together with the notable differences in size, weakens the case for common descent of the *G. obscuriglobus* pores and eukaryotic nuclear pores. An alternative interpretation to deep shared evolutionary origin of pores and pore complex proteins is that the similarities we observe between the *G. obscuriglobus* pore complex and the eukaryote nuclear pore complex are the result of convergence spanning all the way from gross architecture down to the recruitment of individual protein components with shared domain architecture. Under this interpretation the *G. obscuriglobus* pore complex is an analog of the eukaryote nuclear pore complex rather than a homologous structure derived by descent from a common ancestor. In other words, similar solutions to macromolecular transport across internal membranes and between cell compartments may have evolved independently, once in eukaryotes and once in planctomycete bacteria, via the parallel co-option of protein folds present in both bacteria and eukaryotes. The structures we report support the view that these solutions are architecturally similar, and in this regard, further characterization of the *G. obscuriglobus* pore will be critical for understanding the evolution of internal cell structure in this bacterium. The pore structures we have found are not straightforward to explain within the context of recent exciting results regarding Gram-negative bacteria cell wall components in planctomycetes, but there is no evidence from our data consistent with a concept that pore-containing membranes are outer membrane or other cell wall components, or that the pores contain any wall-specific proteins. Such pore structures are however consistent with the known unique internal cell structure features of planctomycetes, and consistent with distinctive planctomycete cell organization and functional properties. Significantly, when considered alongside the recent discoveries of endocytosis [24] and compartmentalized transcription and translation [13], the parallels between the cell biology of *Gemmata obscuriglobus* and that of eukaryotes are nothing short of remarkable, whether due to hidden homology or an analogous reuse of a similar set of protein folds. Detailed analysis of the composition of the internal bacterial pores will no doubt enable the evolutionary origin of these structures to be definitively established.

## ACKNOWLEDGEMENTS

The authors acknowledge the facilities, and the scientific and technical assistance, of the Australian Microscopy & Microanalysis Research Facility (AMMRF) at the Centre for Microscopy and Microanalysis, The University of Queensland. We thank Dr. Vitaliya Sagulenko for help with Western-blot analyses. We thank Tony Romeo of Electron Microscope Unit (EMU), University of Sydney, for help with preparing some of the freeze-fracture replicas. JAF acknowledges assistance in developing some preliminary density gradient protocols and donation of gp4978 antibody from Jody Franke now of Creighton University, Omaha, Nebraska USA.

## MATERIALS AND METHODS

### Cell lysis and electron microscopy of nuclear body envelopes

In the case of mechanical lysis experiments, *Gemmata obscuriglobus* was grown on M1 agar for 6 days at 28 ˚C, and growth harvested by washing into filtered Milli-Q water. For mechanical lysis via grinding with alumina, suspensions were mixed with alumina powder (Sigma Alumina Type A-5) in an eppendorf tube and a plastic pestle manually rotated within the tube for ca. 60 sec to lyse cells. The supernatant from this homogenate after allowing particles to settle was either harvested directly for negative staining or this supernatant purified by centrifugation using a microfuge at 20160 g. Negative staining was performed by mixing a homogenate drop on a pioliform–coated specimen support grid with 1% uranyl acetate containing 0.4% sucrose. In the case of preparations that were sonicated, *G. obscuriglobus* culture was grown on M1 medium for 8 days at 28˚C. Cells were suspended in 1 ml of sterile Milli-Q water, and sonicated in a Branson Sonifier 250 at amplitude output level 1 for ten thirty second intervals, separated by 30 sec rests. The resultant suspension was pelleted in a benchtop centrifuge for five mins at 20160 g, and the pellet was resuspended in 50µL sterile Milli-Q H_2_O. Cells from these suspensions were negatively stained using 2% ammonium molybdate pH6.5 (mixture of 5µl each of suspension and stain was prepared on a carbon-and pioloform-coated specimen support grid followed by removal of excess fluid with filter paper and air-drying).

### Cryo-electron microscopy

4 µL aliquots of density-gradient-purified membrane fractions from sonication-lysed *G. obscuriglobus* cells were deposited onto holey carbon films on hexagonal 200 mesh copper grids (‘C-flat’, Protochips, NC) in the humidified chamber of a commercial thin-film vitrification apparatus (CP3, Gatan, Pleasanton, CA). An additional 4 µL of colloidal gold (nominal size 10-12 nm) was added to the membrane fraction and the grid was blotted from both sides for 3.5 seconds before automatic propulsion into liquefied ethane. Grids were transferred under liquid nitrogen to a cryo-sample holder (Model 914, Gatan) for transfer to the microscope and observation at a stable temperature of approximately −175 Celsius. A JEM-1400 transmission electron microscope (JEOL, Japan) fitted with a high-contrast polepiece and LaB6 cathode, and operating at an accelerating voltage of 120 kV was used for data acquisition. Micrographs were recorded at a nominal defocus value of −10 µm and an electron dose of approximately 4 000 electrons/nm2/micrograph. Detector noise was reduced by means of a median filter with a radius of 1 pixel.

### Electron tomography from thick sections

For electron tomography of isolated membrane samples, 300nm thick sections were cut using Leica EM UC6 ultramicrotome (Leica Microsystems, Austria). The membranes were cryofixed as described for thin-sectioning. Dual-axis tilt-series data was collected at 39000x magnification on an FEI Tecnai F30 (FEG)TEM (FEI Company, the Netherlands) operating at 300kV, over a tilt range of +/−66º at 1º increments for the a-axis and 2º increments for the b-axis, using SerialEM software (The Boulder Lab for 3D Electron Microscopy, USA).

### Membrane fractionations

Cells were collected from two to three M1-agar plates, washed once, and resuspended in 500 μL of bt-DMSO buffer [10mMbis-Tris (pH 7.5), 0.1mMMgCl2, and 20% DMSO] supplemented with 10 μL of protease inhibitor mix (Protease Inhibitor Mixture Set 3; Merck), 10 μg of DNase, and 10 μg of RNase. Cells were then sonicated using a Branson Sonifier 250, and unbroken cells were spun down in a microfuge at 5,000 × g for 10 min. Supernatant was centrifuged at 100,000 × g in a Beckman Coulter tabletop ultracentrifuge (Optima TLX). The pellet was resuspended in 500 μL of bt-DMSO buffer, loaded onto a five-step sucrose gradient, and centrifuged in an SW60 rotor on a Beckman Coulter L8-60M ultracentrifuge at 215,000 × g for 4 h. Visible pink or white membrane fractions were collected via puncturing the side of the tube and to remove sucrose, centrifuged at 100,000 × g in a Beckman Coulter tabletop ultracentrifuge (Optima TLX), and the pellets were resuspended in ≈500 μL of bt-DMSO buffer. The material was loaded onto a 20–60% or 30-70% sucrose/bt-DMSO continuous gradient. After centrifugation using SW60 rotor, for 16 h at 215,000 × g, ≈400 μL fractions were collected from a puncture at the bottom of the tube, To concentrate and remove sucrose, fractions (in all cases containing a visible band) were diluted with bt-DMSO buffer and centrifuged at 100,000 × g in a Beckman Coulter tabletop ultracentrifuge (Optima TLX). The pellets were resuspended in 5mM Tris, pH7.5 and immediately used for electron microscopy experiments or proteomics. Each fractionation was performed twice using separate culture batches of cells, and 2-3 technical replicates from each fractionation were used in proteomics experiments.

### Analysis of structural symmetry

To assess symmetry of structure within pore-like structures within a pore-containing membrane fraction purified from membranes released from lysed *G. obscuriglobus* cells, Markham rotation [58](Markham et al., 1963) was performed according to the modifications suggested by Friedman [59](Friedman, 1970) by assuming any possible symmetry up to and including 8-fold symmetry. Pores in isolated pore-containing membranes from fraction 3 from the membrane fractionation described above were imaged *en face* by cryo-electron microscopy and were extracted from images and rotated in silico by set increments corresponding to the assumed symmetry. For example, reinforcement of possible 8-fold symmetry was assessed by superimposition of the corresponding 45.0 degree rotations, 7-fold by 51.4 degrees and so on. The procedure was conducted on raw data only rather than the bandpass-filtered structures. ImageJ was used for all feature extraction, rotation and summation.

Preparation of material used for Markham rotation: Isolated membranes from fraction 3 of the density gradient fractionation of membranes from lysed *Gemmata* cells were processed for cryo-electron microscopy and symmetry analysis via Markham rotation. Briefly, isolated membrane sheets were vitrified by rapid immersion in liquid ethane prior to mounting the samples in a cryo-sample holder (Model 914, Gatan, Pleasanton, CA) and imaging the frozen-hydrated specimens in a JEM-1400 transmission electron microscope (JEOL Ltd, Japan) equipped with a charge-coupled device detector (Gatan) and low-dose exposure conditions (< 4,000 electrons/nm2) to avoid radiation damage. The accelerating voltage was 120 kV and the nominal magnification of 15,000 corresponded to a pixel size of 0.69 nm at the detector. A minority of pores appeared to be markedly ellipsoid in projection. This was found to be the result of a tilted or folded membrane, which was taken into account by manual tilting of the specimen. Note that this differs from slight deviations in circularity that probably represent the respective functional states of transport-competent pores[60].This tilt was always less than 10 degrees, indicating that the untilted membrane sheets were mounted approximately orthogonal to the beam.

### Freeze-fracture electron microscopy of whole cells

Cells from a 14-day culture of *G. obscuriglobus* ACM 2246 were grown on soil extract agar medium. Cells were harvested directly without chemical fixation into 20% (v/v) aqueous glycerol as cryoprotectant prior to freezing in liquid Freon 22. Fracturing was performed using a Balzers BAF 301 apparatus fitted with a complementary fracturing device, at −115 ˚C and 10^−7^ torr (1 torr = 133 Pa). Replicas were produced using platinum/carbon and stabilized with a layer of carbon.

### Cryosubstitution and thin-sectioning

For cells cryofixed by plunging into liquid propane, cells of *G. obscuriglobus* were cryofixed using a Reichert-Jung KF80 cryofixation system. Cryosubstitution was performed with 2% osmium tetroxide in molecular-sieve-dried acetone at −79 ˚C (dry ice/acetone bath) for 50 hr. The temperature was increased to −20 ˚C over 14 hr. Specimens were brought to room temperature and then washed in acetone. Specimens were embedded in Epon resin, then sectioned and stained on pioloform-covered specimen support grids with aqueous uranyl acetate and lead citrate. For cells cryofixed by high-pressure freezing, cells of *G. obscuriglobus* were cryofixed by first mixing with 2% agarose and placing the suspension between hexadecene-soaked planchettes, then frozen using a BAL-TEC HPM 010 High-Pressure Freezer. Frozen cells at −160 ˚C were warmed to −85 in 1.9 hrs at 4 C/hr, stored at −85 ˚C for 52 hrs and then raised to −20 ˚C over 11 hrs in a Leica EM AFS cryosubstitution apparatus. Cells were then embedded in Epon resin and sectioned and stained as above.

### Electron microscopy

For experiments other than those involving tomography, specimens were examined using a JEOL 1010 transmission electron microscope operated at 80 kV.

### Proteomics

Before proteomics, pellets of membrane fractions in Tris buffer were dissolved using Laemmli buffer, protein concentration was measured using BCA Protein assay (ThermoFisher Scientific) and 20 µg of each suspension was loaded onto PAA gels. **T**he proteins, separated by 1-D SDS-PAGE electrophoresis, were cut out from the 8-12% PAA gels for mass spectroscopy. Gel slices were destained with 50% ACN in 50 mM ammonium bicarbonate (ABC) followed by dehydration in 100% acetonitrile (ACN). Trypsin (80 ng) in 50 mM ABC was added and gel slices rehydrated at 4 °C for 10 min, followed by incubation at 37 °C overnight. Peptides were extracted three times with 50 ul of 50% ACN / 0.1% formic acid, followed by clean up with a ZipTip (Millipore). Peptides were separated using reversed-phase chromatography on a Shimadzu Prominence nanoLC system. Using a flow rate of 30 µl/min, samples were desalted on an Agilent C18 trap (0.3 x 5 mm, 5 µm) for 3 min, followed by separation on a Vydac Everest C18 (300 A, 5 µm, 150 mm x 150 µm) column at a flow rate of 1 µl/min. A gradient of 10-60% buffer B over 30 min where buffer A = 1 % ACN / 0.1% FA and buffer B = 80% ACN / 0.1% FA was used to separate peptides. Eluted peptides were directly analysed on a TripleTof 5600 instrument (ABSciex) using a Nanospray III interface. Gas and voltage settings were adjusted as required. MS TOF scan across m/z 350-1800 was performed for 0.5 sec followed by information dependent acquisition of the top 20 peptides across m/z 40-1800 (0.05 sec per spectra). Data were converted to mgf format and searched in MASCOT accessed via the Australian Proteomics Computational Facility and searched against the LudwigNR database, limited to ‘other bacteria’, using trypsin as enzyme, 2 mis-cleavages, MS tolerance of 0.5 Da and MS/MS tolerance of 0.2 Da. Oxidation (met, variable) and carbamidomethylation (cys, fixed) modifications were also included.

The MS analyses performed were strictly qualitative, not quantitative, and therefore no normalisation of protein amount prior to trypsin digested was performed. A nominal amount (eg 80 ng) trypsin is added per gel slice, as it is not feasible to determine the amount of protein in each gel band processed for MS. This is typical practice in the proteomics field. Samples were ziptipped after digestion, prior to MS, in part to desalt/concentrate the samples, but also to ensure the LCMS system was not overloaded (ziptips have a limited loading capacity (5ug)).

### Antibodies

A polyclonal antibody (designated as Anti-Protein 6670) was raised by GenScript Inc. (Piscataway, NJ, USA). The antigen used for immunization was VPVTDDTRKEPTETC, derived from the translated protein ZP_02736670. Immunogen was a Peptide-KLH conjugate, and host strain was New Zealand rabbit. The antibody was affinity purified and stored in Phosphate Buffered Saline (PBS, pH 7.4) with 0.02% Sodium Azide at −20^°^C. Membrane preparations of *G. obscuriglobus* were resolved by SDS/PAGE (10%) and the specificity of the antibody was tested via western blotting at 1:2000 dilution (S10 Fig). HRP-conjugated goat anti-rabbit antibody (Cell Signalling Technology, Australia, 1:5000 dilution) was used as secondary antibody. Detection was done by using ECL Western Blotting System (GE Healthcare Life Science). The ‘anti-MC protein’ antibody against the Gemmata obscuriglobus protein gp4978, a clathrin heavy chain-like membrane coat (MC) protein already shown to be present on internal membrane vesicles of *Gemmata obscuriglobus* associated with endocytosis-like protein uptake in this species[24]. This antibody to gp4978 was a rabbit polyclonal antibody raised against recombinantly expressed and purified gp4978 protein identified as a eukaryotic MC coatomer protein (National Center for Biotechnology Information reference sequence: ZP_02732338.1; see [37]).

### Immuno-gold labelling

Ultrathin sections of high-pressure frozen and cryosubstituted intact *G. obscuriglobus* cells or density gradient-purified fraction 3 membranes on formvar-carbon-coated copper grids were floated onto drops of Block solution containing 0.2% (wt/vol) fish skin gelatin, 0.2% (wt/vol) BSA, 200 mM glycine, and 1× PBS on a sheet of Parafilm, and treated for 1 min at 150 W in a Biowave microwave oven. The grids were then transferred onto 8 µL of primary antibody, diluted 1:25 in blocking buffer, and treated in the microwave at 150 W, for 2 min with microwave on, 2 min off, and 2 min on. The grids were then washed on drops of Block solution three times and treated in the microwave at 150 W each time for 1 min before being placed on 8 µL of goat anti-mouse IgG Fc (γ)-specific antibody conjugated with 10 nm gold (British Biocell International, catalog no. EM GAM10) diluted 1:50 in Block solution and treated in the microwave at 150 W, for 2 min with microwave on, 2 min off, and 2 min on. Then grids were washed three times in 1× PBS, each time being treated for 1 min each in the microwave at 150 W, and four times in water for 1 min each in the microwave at 150 W, and examined via transmission electron microscopy.

## LEGENDS FOR SUPPLEMENTARY VIDEOS AND FIGURES

**S1 Video: Electron tomography of the fraction 3 membranes containing pores**

Curving sheets comprising isolated membranes from fraction 3 (isolated via density gradient fractionation from lysed *G. obscuriglobus* cells) can be seen in section. As tomogram slices of the spirally coiled sheets are passed through during the movie’s course (in effect passing through successive and different planes of a thick section), they can be seen to be interrupted periodically by non-membranous pore structures some regions of which project from one side of each of the membranes which comprise a coil. If one membrane sheet is examined at different points of the movie, several pores can be identified as slices of the membrane sheet are passed through successively. Bar, 50 nm.

**S2 Video: 3D reconstruction of an internal pore-containing membrane of *Gemmata obscuriglobus***

The electron tomography membrane reconstruction shown here is derived from a representative of the fraction 3 membranes in the thick section tilt-series.

S3 Video: 3D reconstruction of a pore embedded in the internal pore-containing membrane of *Gemmata obscuriglobus*.

The reconstruction is derived from a pore in the membrane shown in S1 Video and S2 Video.

**S1 Fig.**
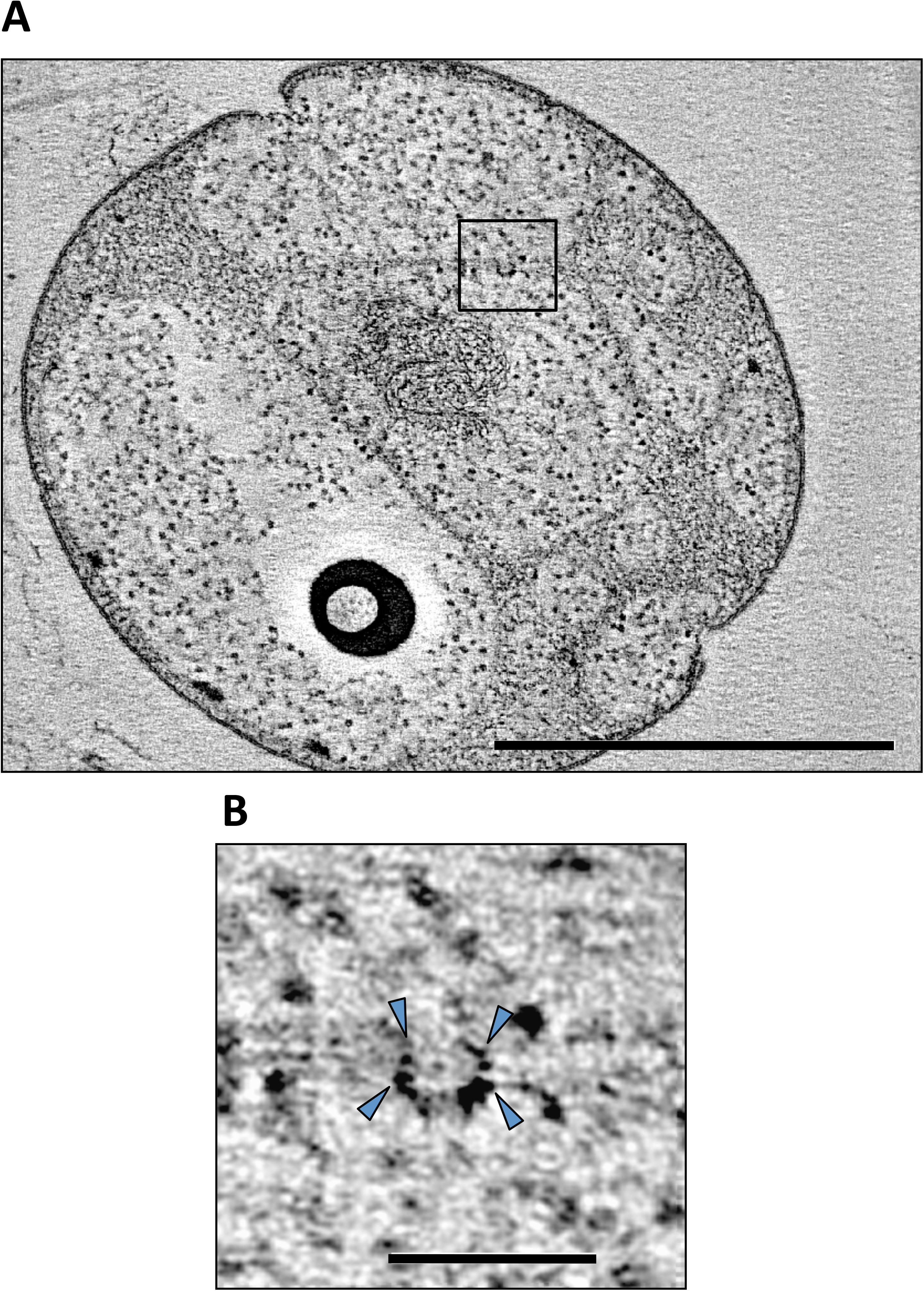
Pores embedded in internal membranes of *Gemmata obscuriglobus*. (***A***)Transmission electron micrograph of a tomographic slice of high-pressure-frozen cryosubstituted thick-sectioned cell showing a pore (boxed region) embedded in internal membranes situated within the cytoplasm and bounding the nuclear body region containing the cell’s nucleoid. Bar, 1 μm. **Inset:** (***B***) Enlarged view of the pore outlined by the box in (A) – the circular pore (arrowheads) is seen tilted *en face* and displays a complex ring structure and a central plug. Bar, 100 nm.

**S2 Fig.**
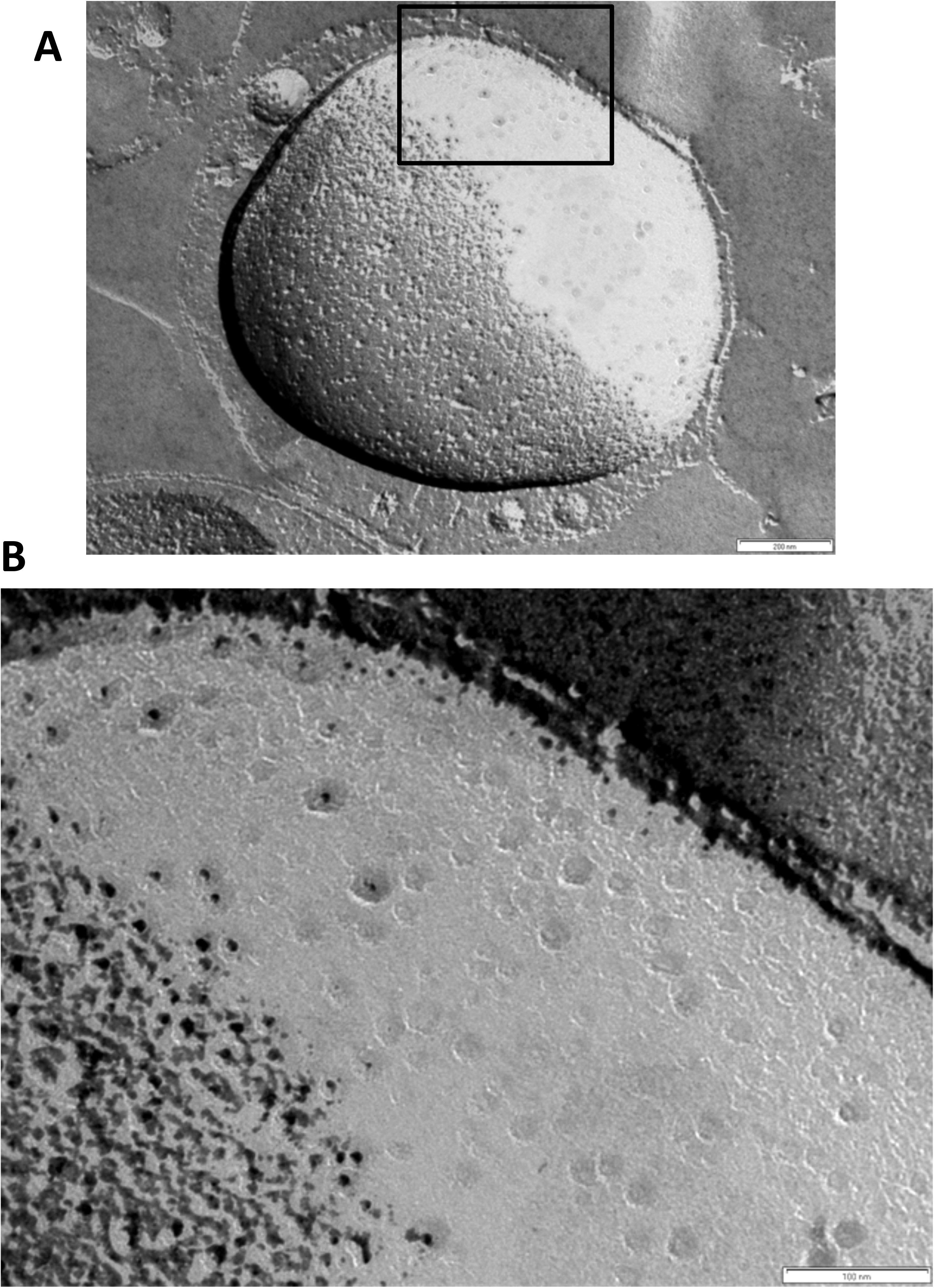
*Gemmata* CJuql4 internal membrane pores as seen in the freeze-fractured cells. **(A)**Transmission electron micrograph of a platinum/carbon (Pt/C)-shadowed replica of a whole cell of *Gemmata* CJuql4 (ACM5157) which has been prepared via the freeze-fracture technique. The fractured whole cell contains a large spherical organelle taking up most of the cell volume, the surface of which has been fractured along a membrane. Pores are visible on the fractured membrane surface of this major cell compartment, interpreted as the nuclear body (e.g. in the boxed region) Bar, 200 nm. (***B***) An enlarged view of the boxed region of the freeze-fractured cell seen in (*A*) showing a region of a membrane surface where pore structures are visible. Several pores display substructure consistent with complex structure including a dark central core and a lighter ring surrounding the core (arrows). Other circular structures in the same size range are also visible but do not present this complex core-ring structure as clearly, presumably reflecting angle at which Pt/C metal shadow has been deposited during formation of the replica after fracture of the frozen cell. Bar, 100 nm.

**S3 Fig.**
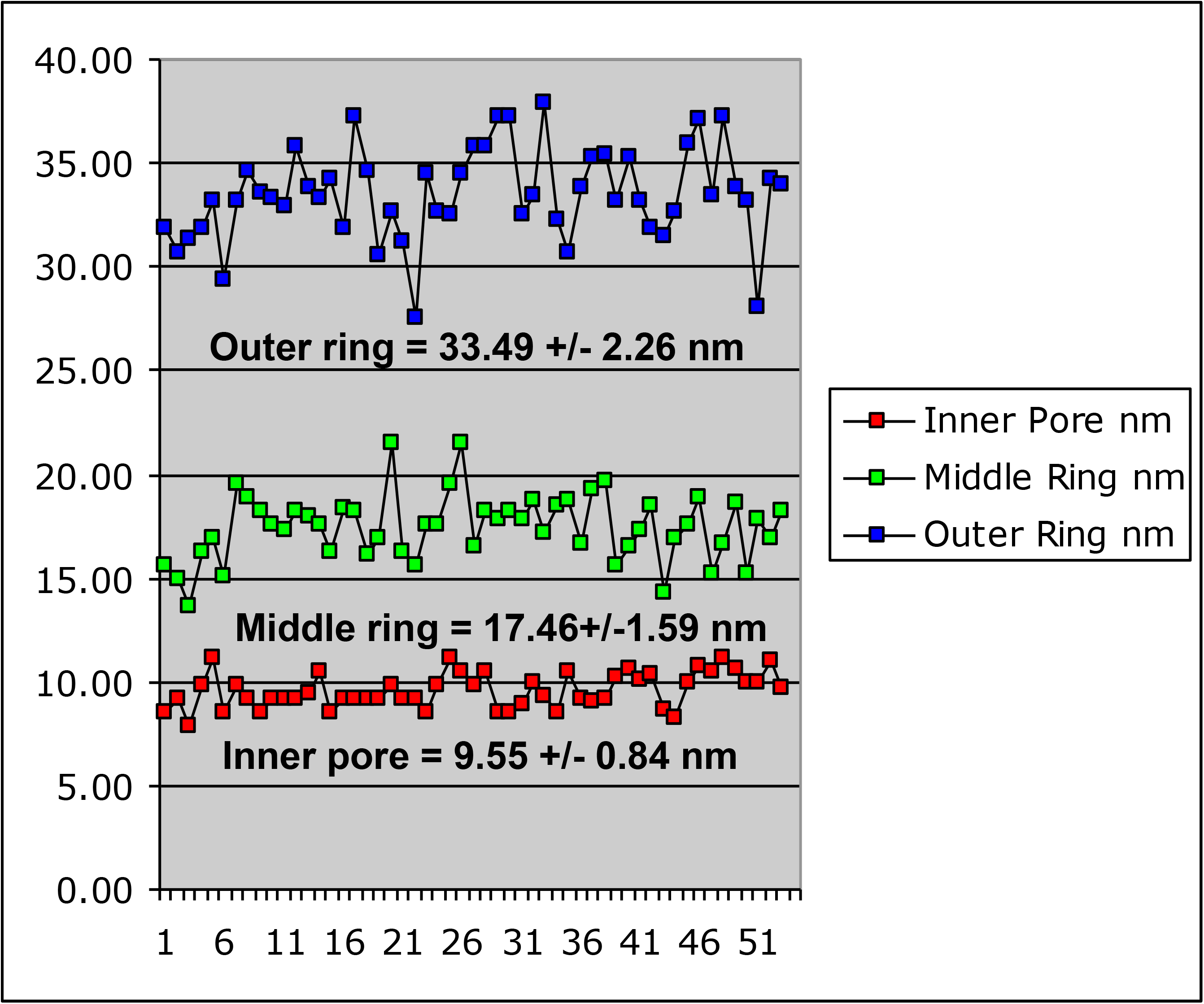
Dimensions of the *Gemmata obscuriglobus* internal pores. The dimensions were calculated from transmission electron micrographs of the membrane fragments released from lysed cells via sonication and negatively stained with ammonium molybdate. The pores usually appear as circular structures with dense pore centers surrounded by a thin lighter inner ring and a thicker outer ring (see Fig 3 for example). The bars are generated automatically and calculated by microscope software.

**S4 Fig.**
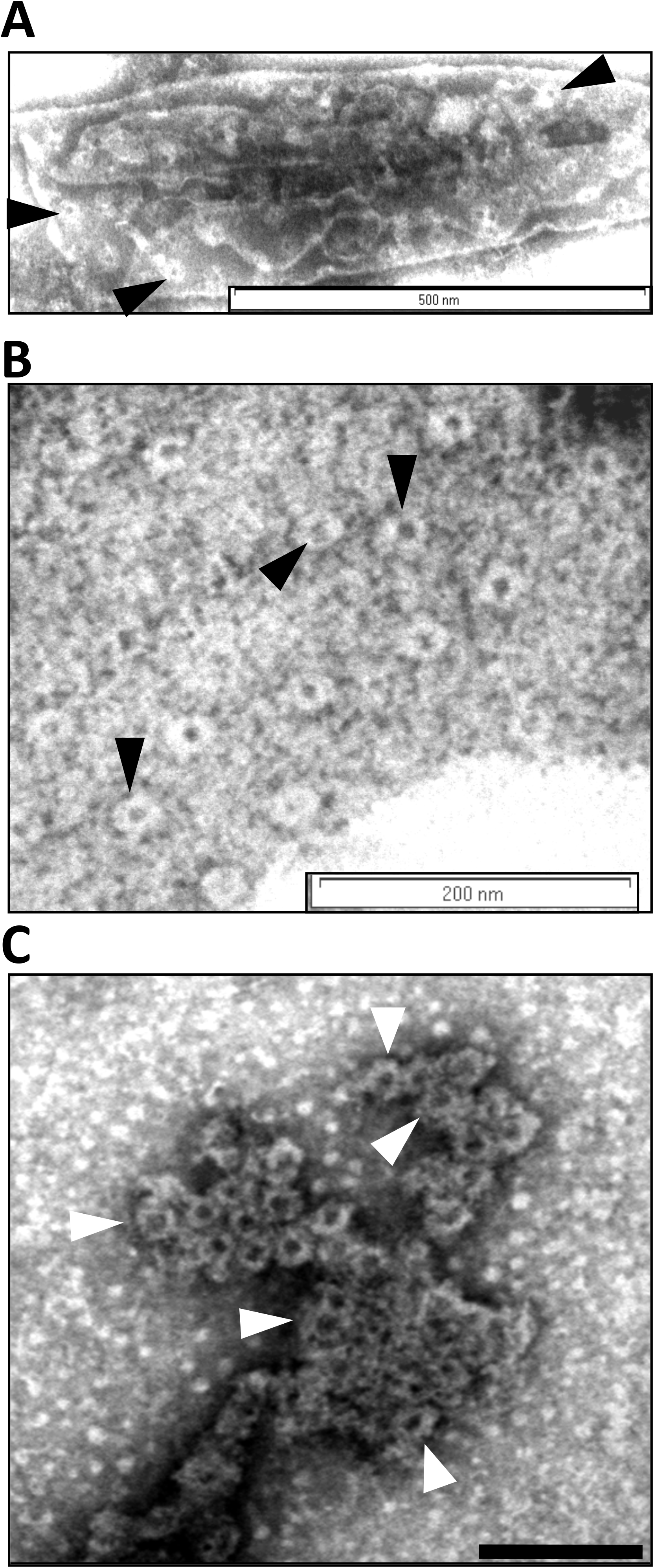
Pore-containing membrane of *Gemmata obscuriglobus* disintegrates and pores aggregate after detergent treatment. **(*A*)** Transmission electron micrograph of negatively stained gradient-fractionated pore-containing membranes purified from sonicated *G. obscuriglobus* acting as control for detergent treatments shown in (B) and (*C*). A “canoe” structure with pores (arrowheads) is visible. Bar, 500 nm. **(*B*)** Transmission electron micrograph of negatively stained gradient-fractionated pore-containing membranes purified from sonicated *G. obscuriglobus* after treatment with 1% Triton X-100 and 1% sodium deoxycholate detergent for 5 min. Pores (arrowheads) are visible within a partially degraded membrane background. Bar, 200 nm. **(*C*)** Transmission electron micrograph of negatively stained aggregated pore complexes seen after treatment of gradient-fractionated pore-containing membranes with 1% Triton X-100 and 1% sodium deoxycholate detergent for 30 min. Individual pores show a central dense core surrounded by a light ring and in some cases material projecting from the outer rim of the ring possibly representing spokes normally connecting inner to outer ring in intact pore complexes (arrowheads). Bar, 50 nm.

**S5 Fig.**
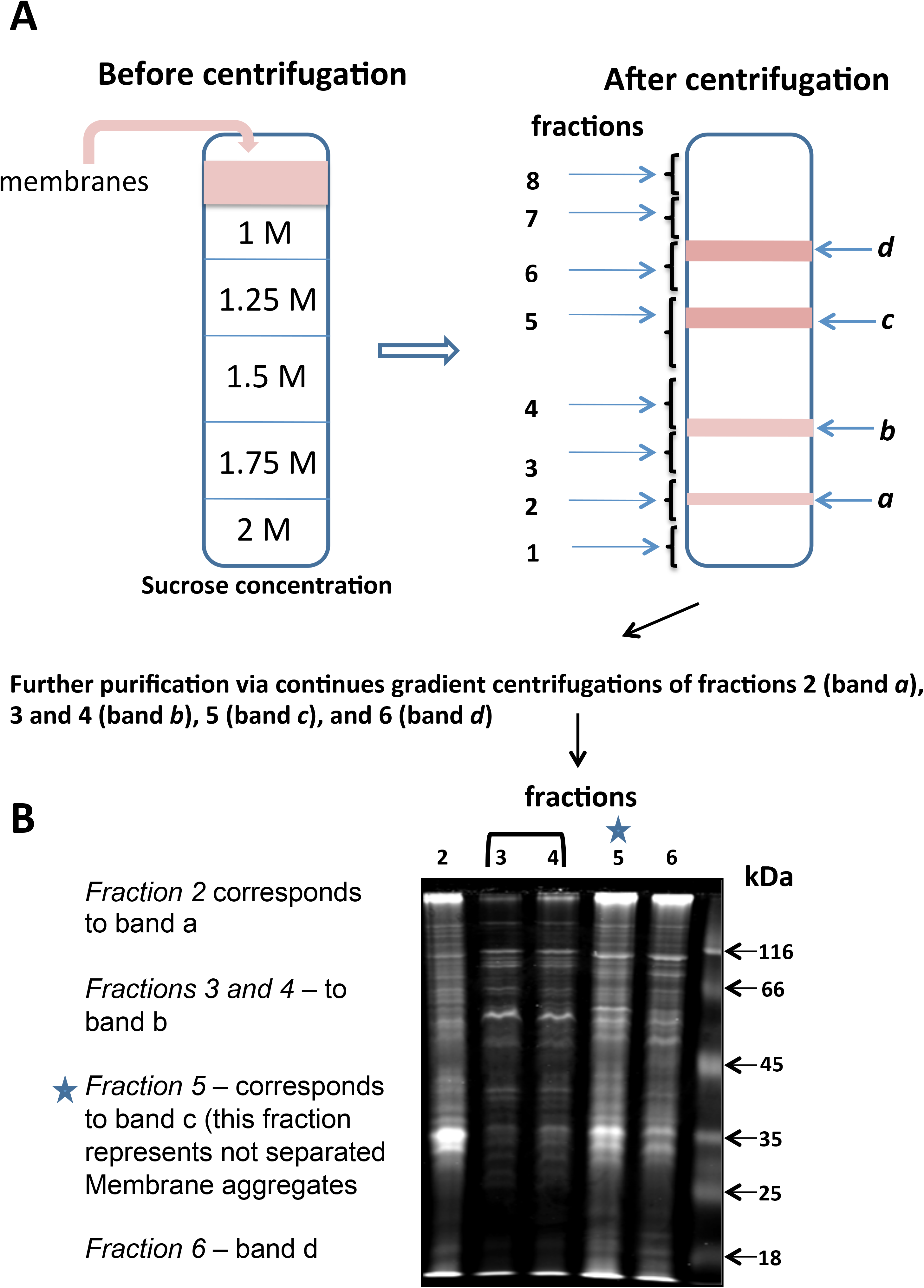
Sucrose gradient fractionation of *Gemmata obscuriglobus* membranes. **(*A*)** Schematic diagram showing bands resulting from density gradient fractionation of membrane fractions released from cells of *G. obscuriglobus* lysed via sonication. On the left is the initial distribution of sucrose concentrations in the gradient before ultracentrifugation and the initial position of the total membrane mixture. On the right are the resulting bands that were visible after ultracentrifugation – fractions collected from the whole length of the gradient are indicated by numbers 1-8 and the resulting protein bands are indicated as *a-d*. **(*B*)** SDS-PAGE gel of continuous-gradient purified fractions corresponding to fractions 2-6 of membrane bands described in (**A**). Bands resulting from electrophoresis of the different membrane fractions show that purified fractions 3 and 4 (band *b*) contain a distinctive pattern of a limited number of proteins relative to fractions 2 (band *a*), 5 (band *c*) and 6 (band *d*). Fractions 1, 7, and 8 did not contain any material and were excluded from further work. Purified fractions 3 and 4 were shown to contain only ‘canoe’ membranes with pore structures via TEM of negatively stained membranes (Fig S8). Protein fraction 5 after continuous gradient fractionation formed a “smear” band which was collected and examined under electron microscope. The collected fraction was found to contain a mixture of membranes morphologically similar to those from fractions 2, 3 (and 4), and 6, and was therefore excluded from proteome analysis. Fraction 4 after preliminary Mass-spec analysis revealed the same protein content as fraction 3, thus for the analysis of the whole protein content of the band *b* we used fraction 3 only.

**S6 Fig.**
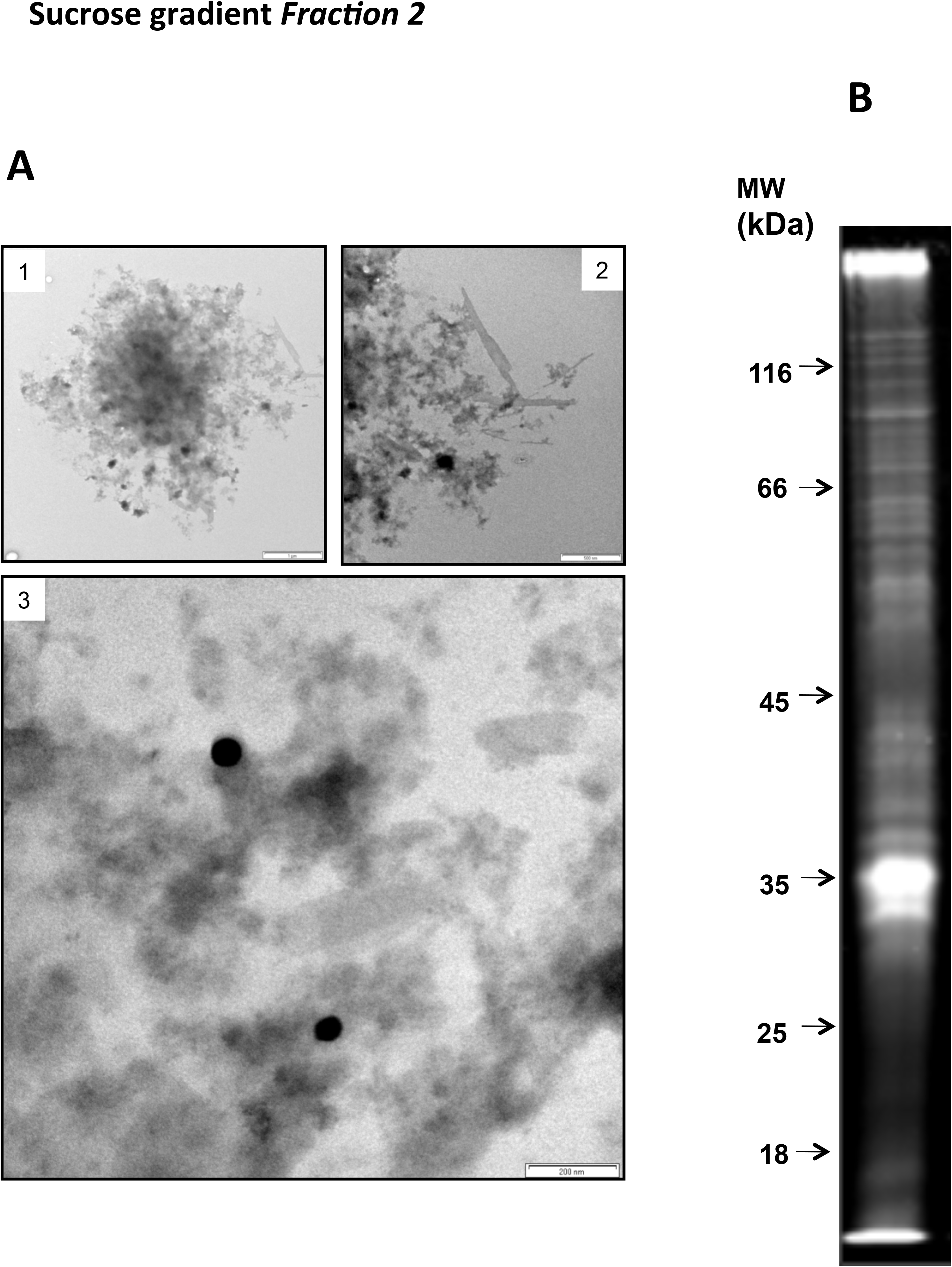
Transmission electron microscopy of the membranes enriched in fraction 2 and SDS PAGE of the proteins obtained from this fraction. **(*A*)** (**1**, **2** and **3)-** transmission electron micrographs of negatively stained membranes of fraction 2 (see Fig S6) containing membranes which do not display pore complexes. Bar A1, 1 µm, Bar A2, 500 nm, Bar A3, 200 nm. **(B)** SDS-PAGE of membrane fraction 2 proteins. All these individual bands were cut out for proteomic analysis (for results see S1 Table).

**S7 Fig.**
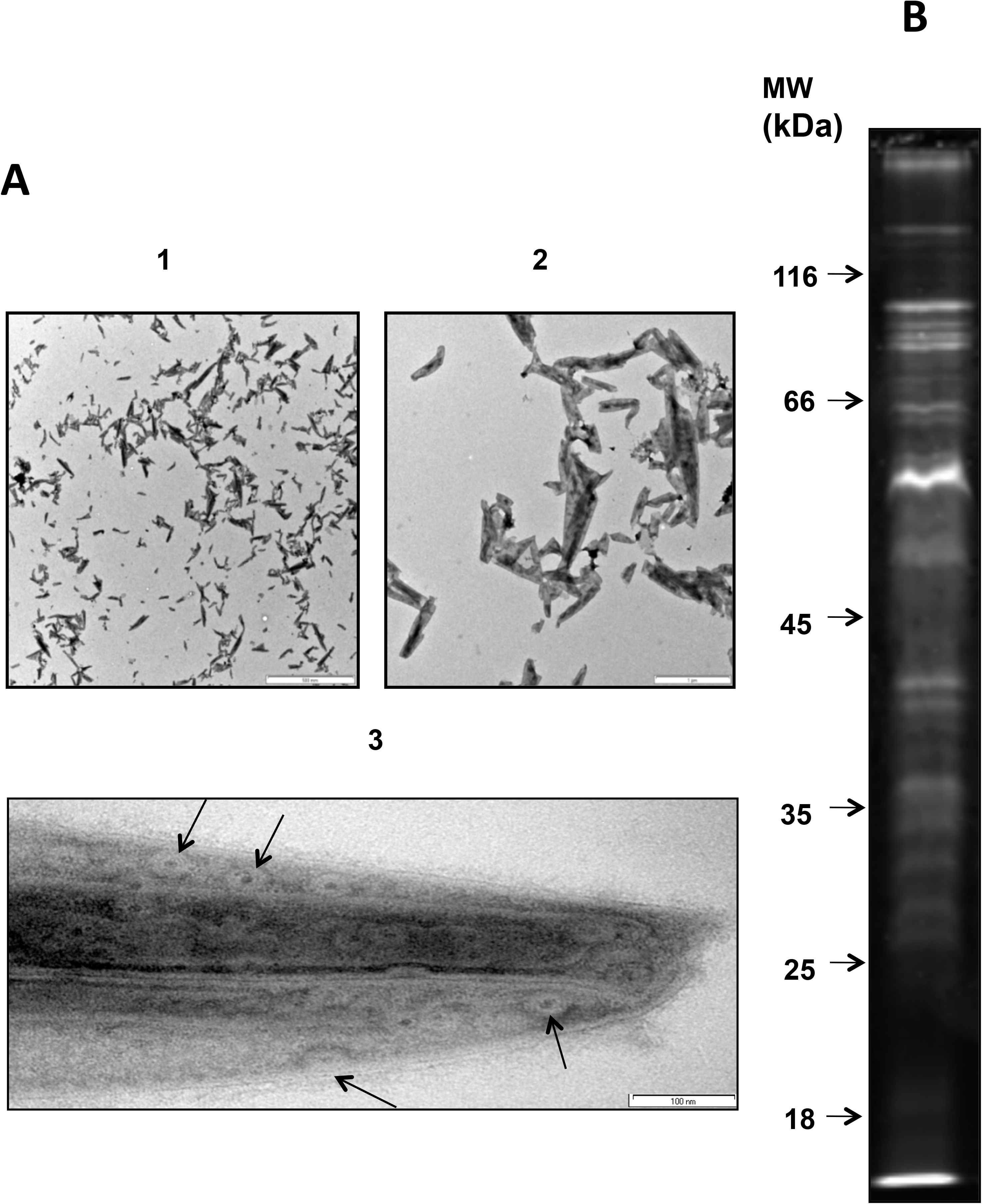
TEM of the membranes enriched in fraction 3 and SDS PAGE of the proteins obtained from this fraction. **(*A*)** TEM of negatively stained membranes of fraction 3 (see Fig S6) containing membranes which display pore complexes. **1** and **2** show appearance of aggregates of membranes at relatively low magnification while **3** shows the characteristic ‘canoe’ shape of pore-containing membranes comprising this fraction. The enlarged view in A3 shows the typical appearance of the large pore ring structures (arrows) on these ‘canoe’-shaped membranes. Bar A1, 5 µm, Bar A2, 1 µm, Bar A3, 200 nm. **(*B*)** SDS-PAGE of membrane fraction 3 proteins. All the individual bands were cut out for proteomic analysis (for results see S1 Table). Fraction 4 contained the same ‘canoe’ shaped membranes and proteomics analysis revealed no difference between those fractions at protein level.

**S8 Fig.**
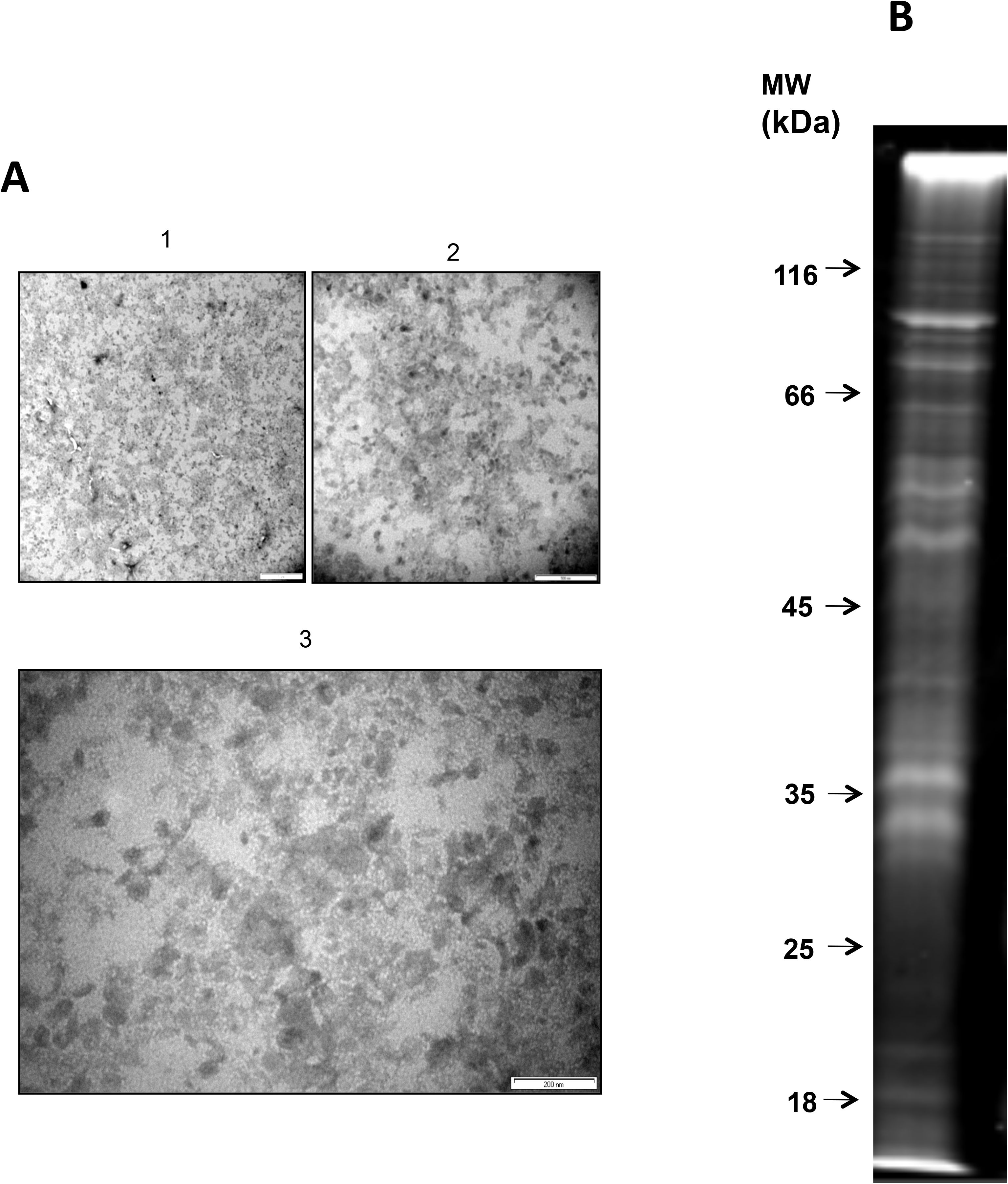
Transmission electron microscopy of the membranes enriched in fraction 6 and SDS PAGE of the proteins obtained from this fraction. **(*A*)** Transmission electron microscopy of negatively stained membranes of fraction 6 (see Fig S6) containing membranes which do not display pore complexes. Bar A1, 10 µm, Bar A2, 500 nm, Bar A3, 200 nm. **(*B*)** SDS-PAGE of membrane fraction 6 proteins. All the individual bands were cut out for preparation for proteomic analysis (for results see S1 Table).

**S9 Fig.**
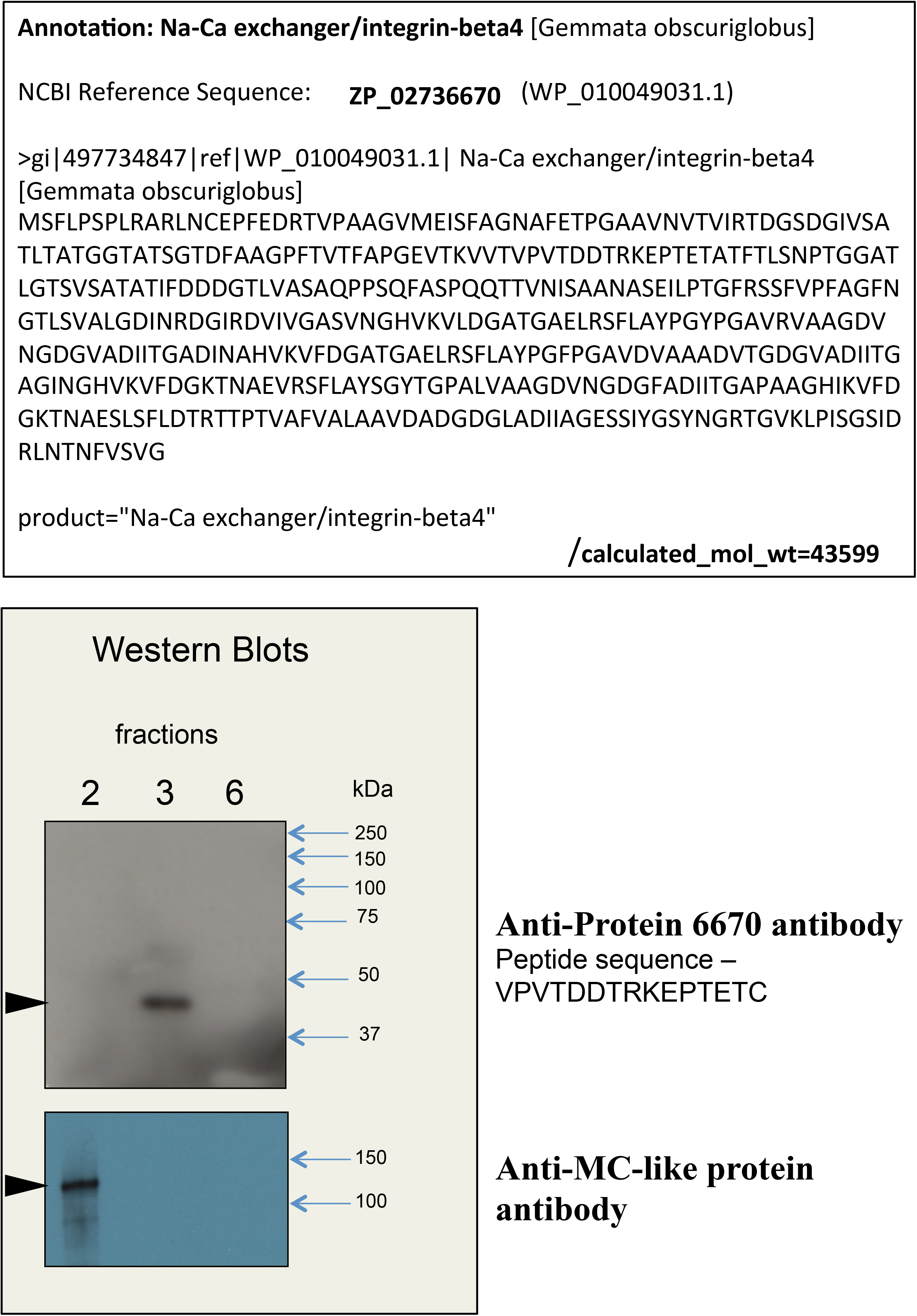
The antibodies 6670 binds specifically to the beta-propeller-containing protein from fraction 3. **(*A*)** Amino acid sequence of the protein annotated as *Na-Ca exchanger/integrin-beta4* (NCBI Reference Sequence: ZP_02736670, later renumbered as the synonymous WP_010049031.1) was used for generation of an antibody (antibody 6670). The protein was identified by mass-spectrometry analyses as a unique protein for Fraction 3 and the full sequence was retrieved from the NCBI Database. **(*B*)** *G.obscuriglobus* fractions were used for testing the antibody specificity. The antibody does not react with proteins from fractions 2 and 6, and only one band was detected in fraction 3 at ca. 40-45 kDa (arrowhead), which is consistent with the predicted MW for the *Na-Ca exchanger/integrin-beta4*. As a control for purity of fractionations the antibody against MC-like protein was tested. The antibody recognizes specifically a protein from fraction 2 with the approximate molecular mass of 120 kDa, which is in accordance with the calculated mass for this protein.

**S10 Fig.**
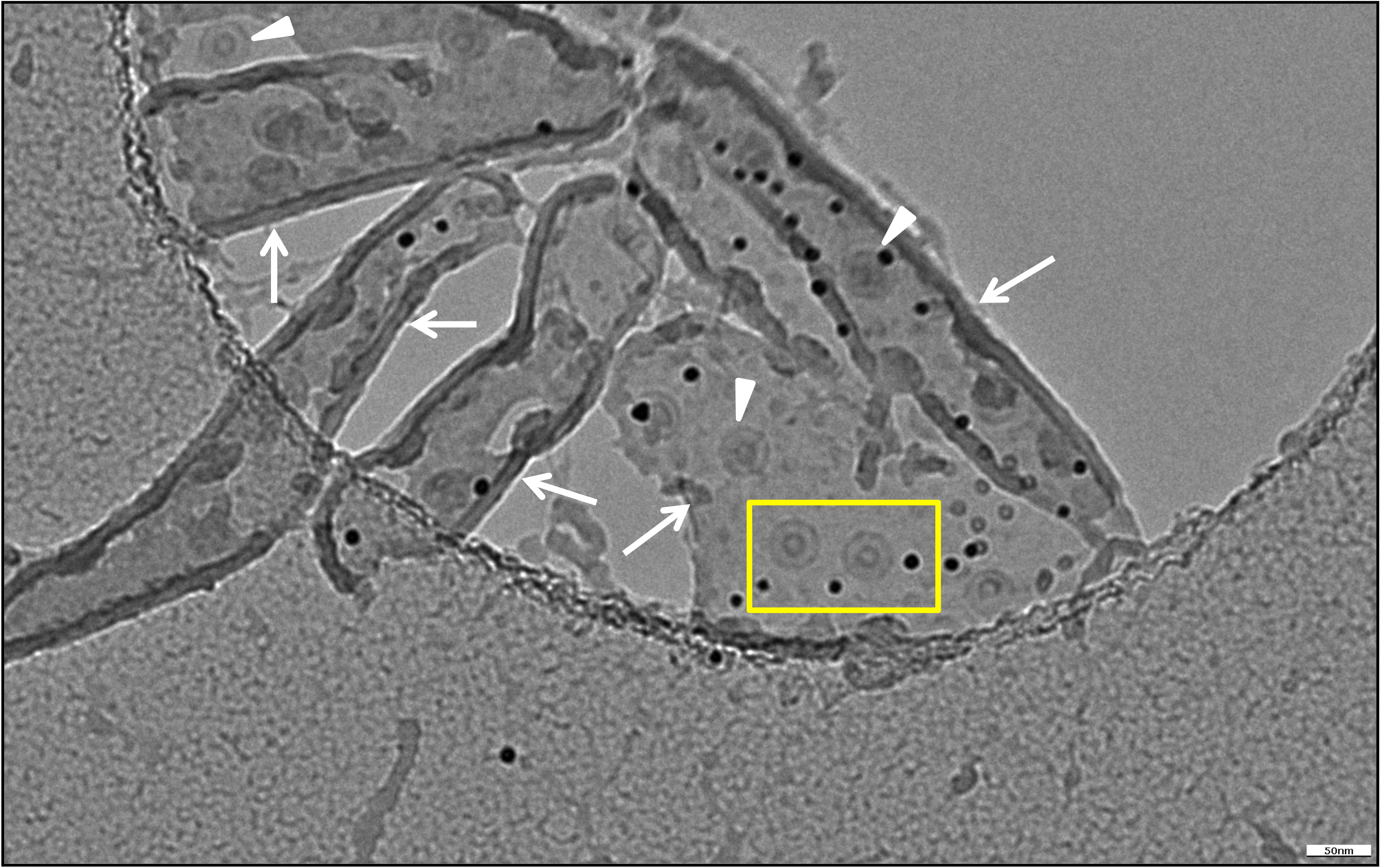
Cryo-EM of fraction 3 isolated membrane sheets. Cryo-EM image of frozen-hydrated sucrose-purified membranes from fraction 3 isolated from lysed cell preparation by density gradient centrifugation. Membrane sheets are indicated by arrows and large pore structures are marked by arrowheads. The two pores in the boxed region are seen in Figure 4C.

**S11 Fig.**
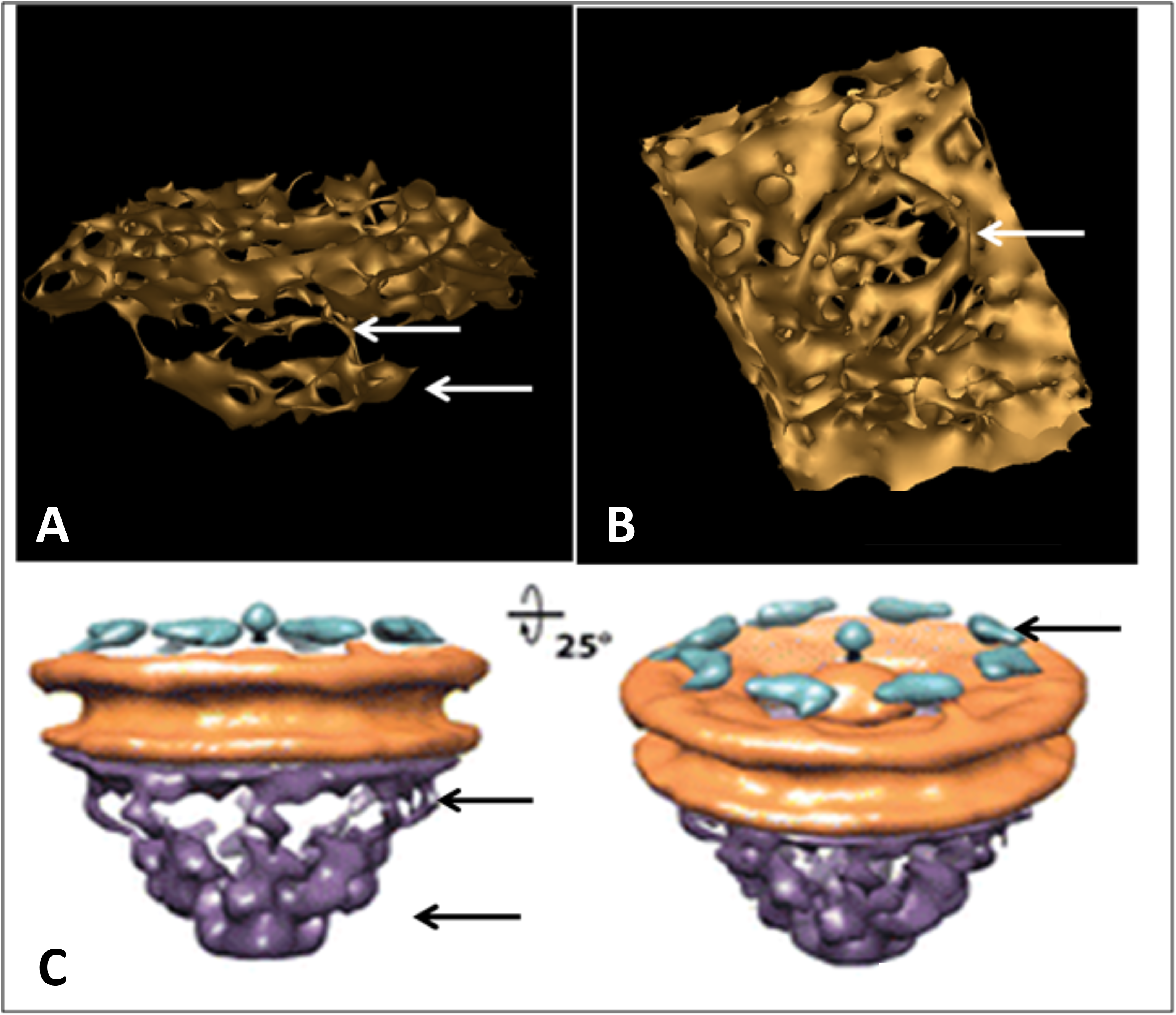
Comparison of *Gemmata* pores with the nuclear pores of eukaryotes. **(*A*)** and **(*B*)** Reconstruction of architecture of a single pore from two different angles. In panel **(*A*)**, a side view of the pore displays the basket structure with a series of struts (arrows) connecting with the main pore rings. In panel **(*B*)**, a top view shows the ring-like element (arrow) of the main part of the pore and a central plug structure is visible within the pore connected to the ring’s inner rim via spokes. The same major pore structural elements (plug, ring and spokes) are indicated in the eukaryote nuclear pore shown in Fig S11C and the pores are shown at similar angles (***C***) The image in (***C***) represents a cryo-electron tomographic reconstruction of the *Dictyostelium discoideum* nuclear pore complex published previously by Beck et al. [26].

**S12 Fig.**
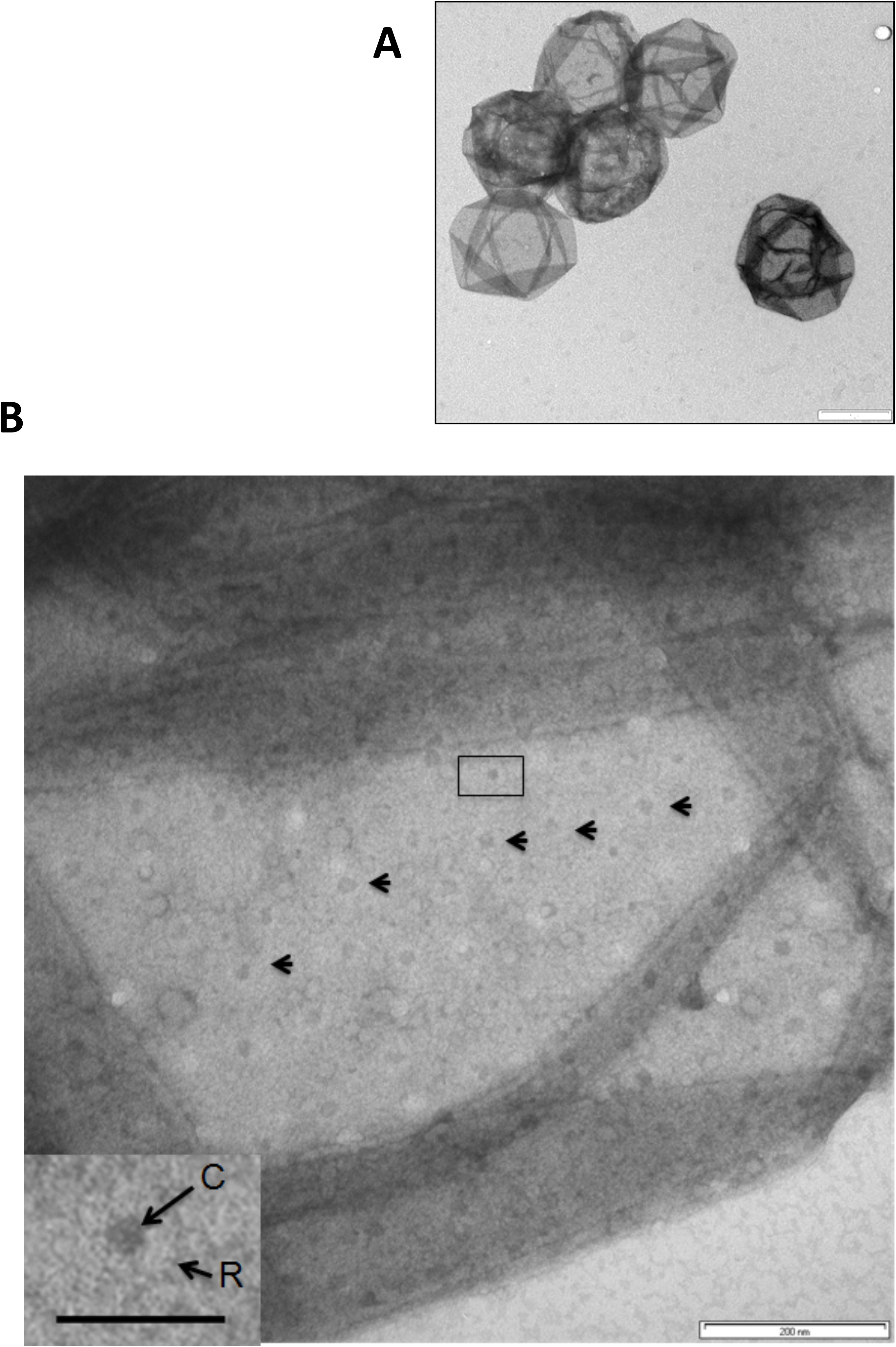
Crateriform structures on the surfaces of *G.obscuriglobus* cell walls. **(*A*)** TEM of cell walls of *G.obscuriglobus* isolated via boiling of bacteria in 10% SDS for 1hr. Bar, 2 µm. (***B***) One of the cell walls of *G.obscuriglobus* with clearly recognizable crateriform structures (arrowheads). The electron dense core regions are variable in shape. Bar, 200 nm. Inset: an enlarged view of the boxed area in A showing a single crateriform structure with central electron-dense core surrounded by a single electron-transparent ring. There is no indication of division of the ring region into an inner and outer ring. Bar, 50 nm.

**S13 Fig.**
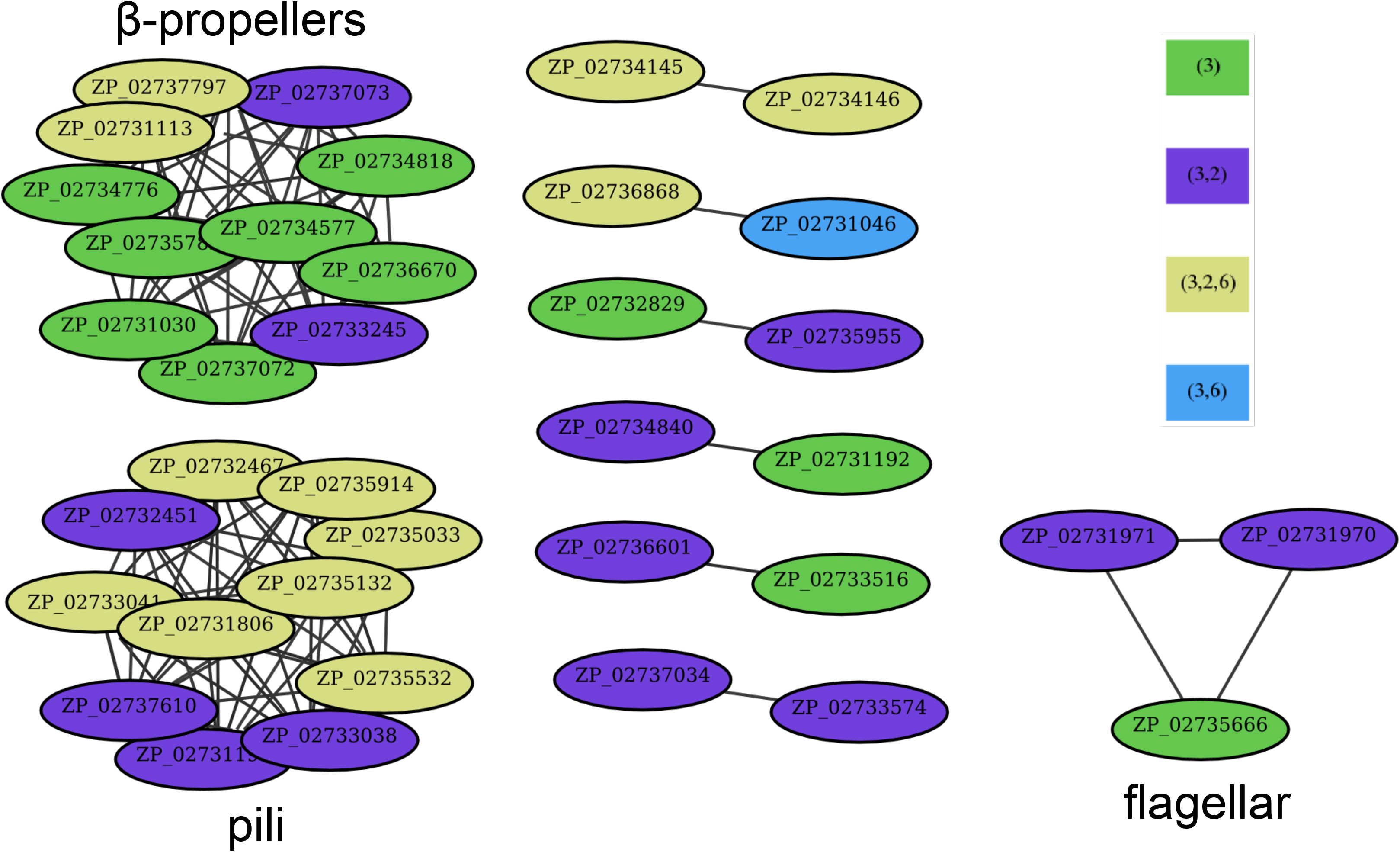
Clustering of membrane-related proteins. All 128 proteins associated with the membrane fractions were clustered using VisBLAST (E = 0.001, i = 2). Proteins are identified by Genbank accession numbers. Lines indicate detectable sequence similarity between proteins. Colour key indicates membrane fractions in which proteins were detected. The 91 singleton proteins that did not show any significant sequence similarity to the other proteins are listed in S3 Table. The cluster in the top left corner (cluster 1) corresponds to the cluster containing fraction 3 proteins from Fig S14. Structural modelling using Phyre2 indicates that constituents of this cluster carry beta-propeller folds (Fig 7C and S4 Table). The cluster on the bottom left is dominated by pili proteins (cluster 2) (see also Fig S18 and S5 Table).

**S14 Fig.**
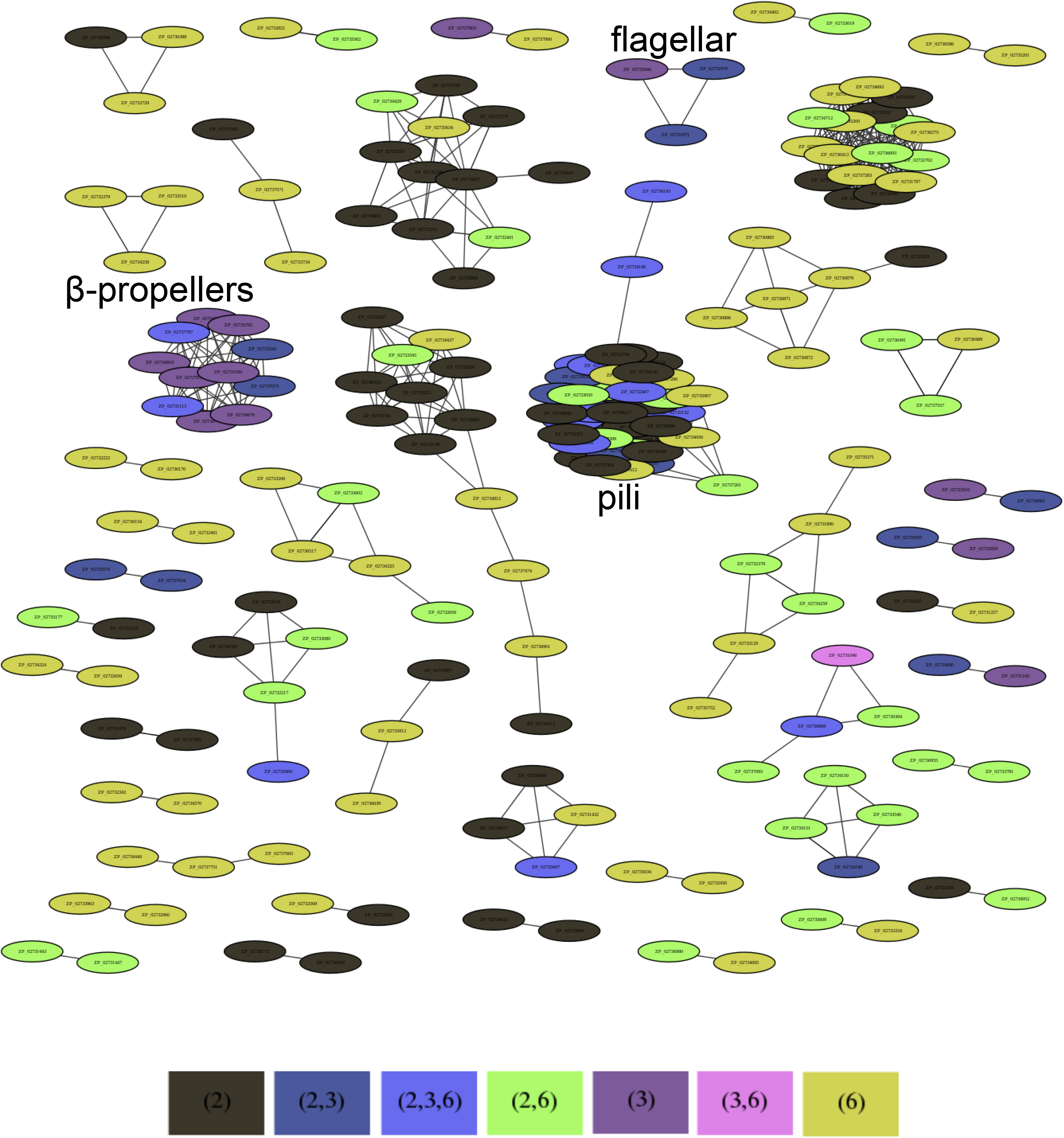
Clustering of all proteins identified through proteomics. All 512 identified in our proteomics analysis were clustered using VisBLAST (E = 0.001, i = 2). This revealed several large clusters, though only one of these contained proteins specific to fraction 3 (cluster 1). Proteins are identified by Genbank accession numbers. Lines indicate detectable sequence similarity between two proteins. Colour key indicates in which membrane fractions proteins were detected.

**S15 Fig.**
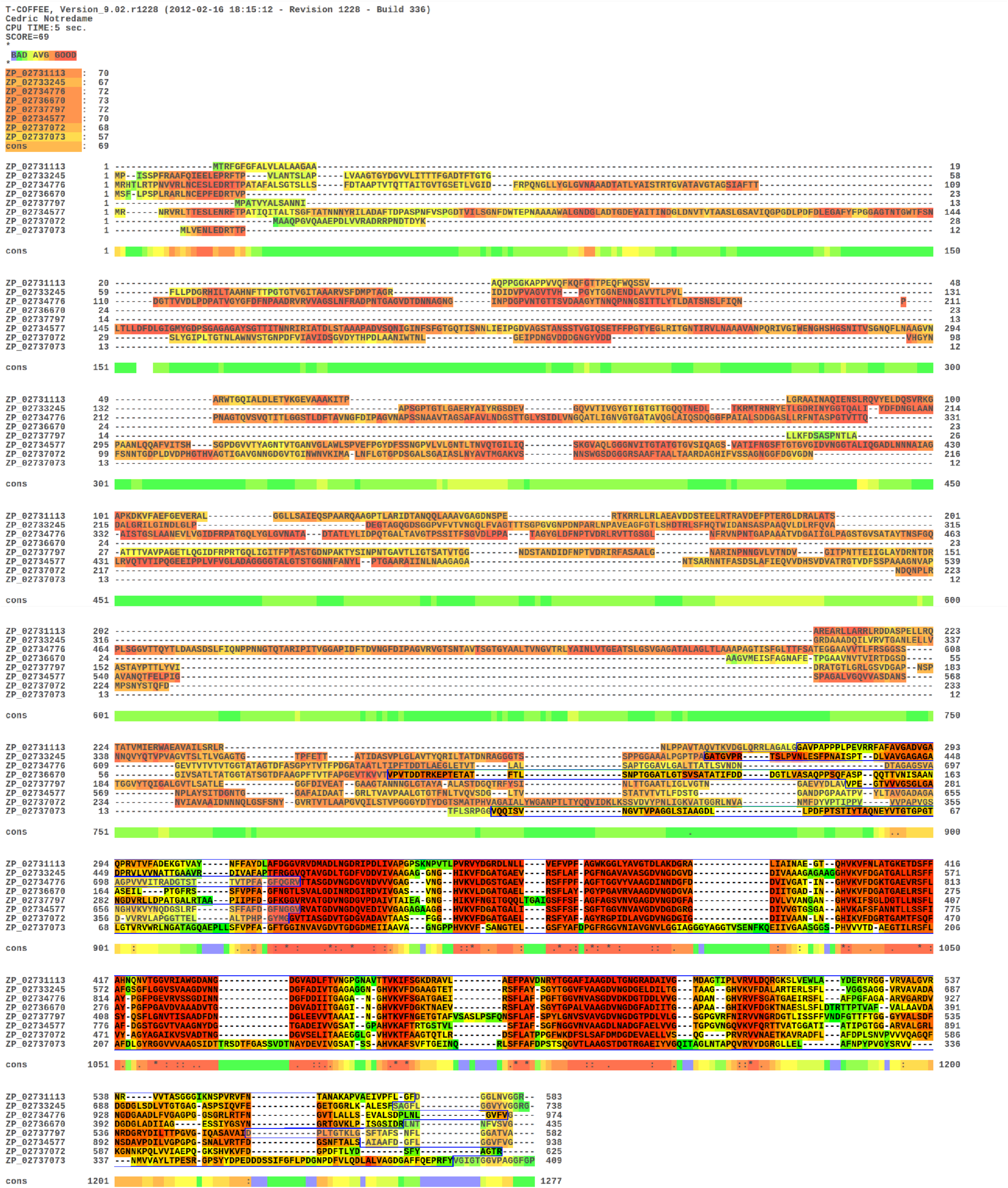
Multiple alignment of the proteins with significant beta-propeller structures. The multiple alignment of the 8 proteins from cluster 1 (top left cluster in Fig S13 and S4 Table) that gave significant structure predictions using Phyre2 is shown. The alignment was made using MAFFT, option L-ins-i, and was evaluated using the T-Coffee CORE server (see Supplementary Text for details). The modelled structures are indicated by the highlighted part of the alignment at the C-terminal end. Structural models (Fig 3) correspond to the conserved part of the sequences.

**S16 Fig.**
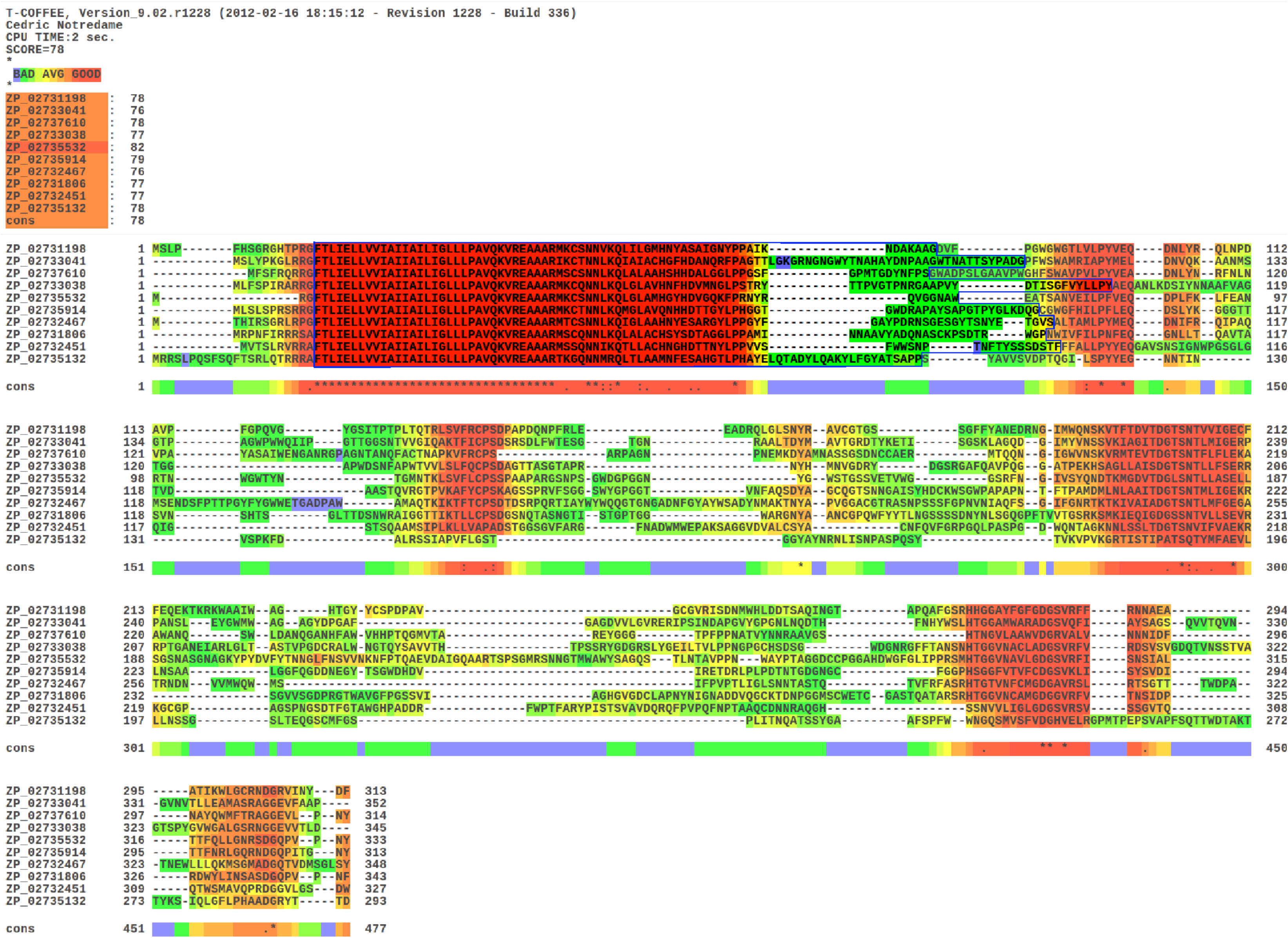
Multiple alignment of the proteins with significant structure predictions in cluster 2. The 10 pili proteins in cluster 2 (bottom left in Fig S13 and S5 Table) that gave significant structure predictions using Phyre2 were aligned using MAFFT, option L-ins-i, and the alignment was evaluated using the T-Coffee CORE server (see text for details). The predicted structures are all clearly located in the conserved subsequence in the N-terminal end of the alignment (see the highlighted part).

**S17 Fig.**
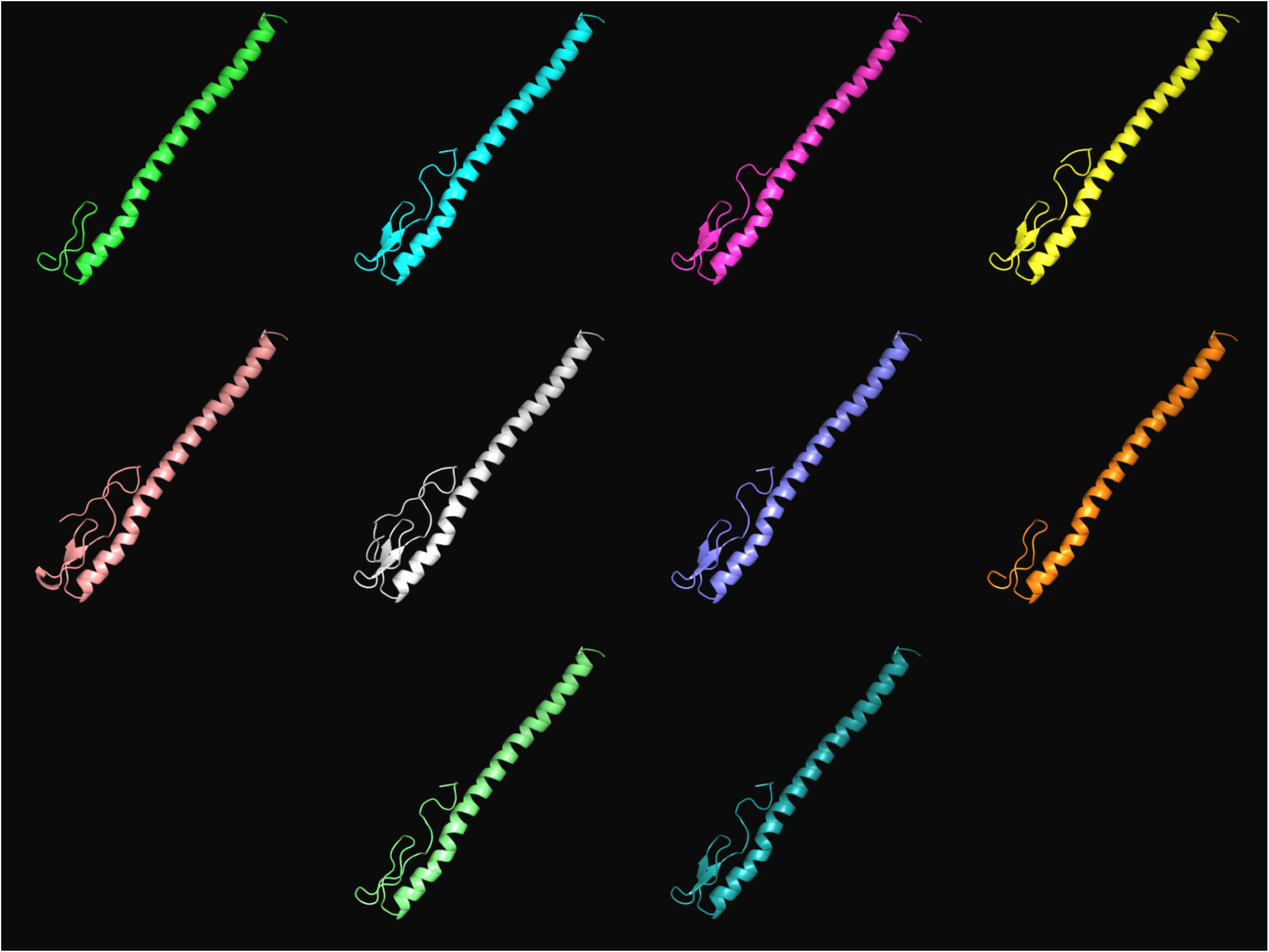
Structures of the pili-proteins from membrane fraction 3. Using PyMOL, the predicted structures for the 11 proteins from membrane fraction 3 that were clustered together and shared a pili-like structure were visualized. None of these are unique to fraction 3 (S5 Table). The structures are for (from left to right and top to bottom): ZP_02731198, ZP_02731806, ZP_02732451, ZP_02732467, ZP_02733038, ZP_02733041, ZP_02735033, ZP_02735132, ZP_02735532, ZP_02735914 and ZP_02737610.

**S18 Fig.**
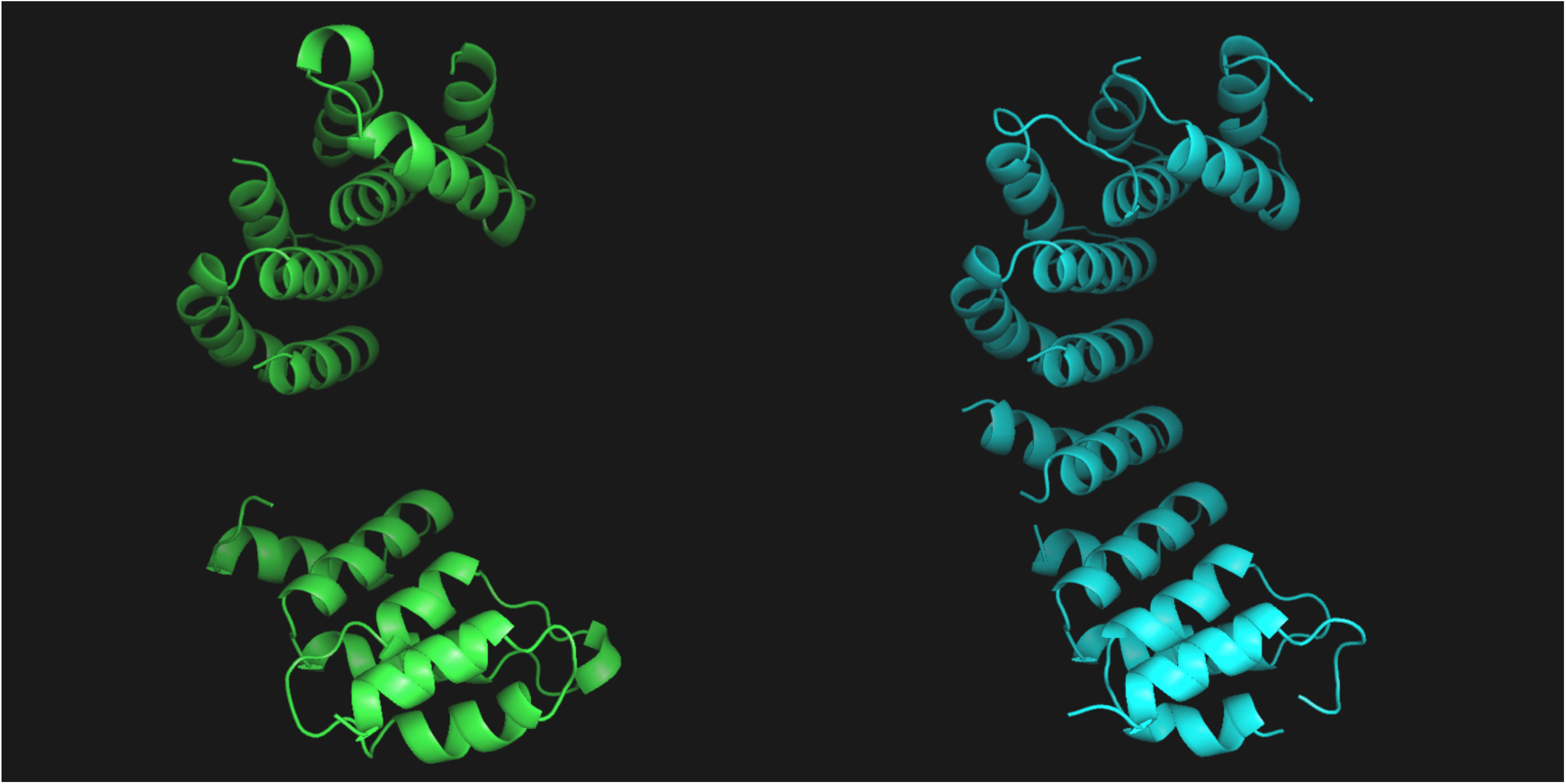
Structures of the two α-solenoids. Two proteins showed a potential α-solenoid structure with stacked α-helices. Left: ZP_02735673 (constituent of fraction 2 and fraction 3) and right: ZP_02736511, unique to pore-containing membrane fraction 3.

**S19 Fig.**
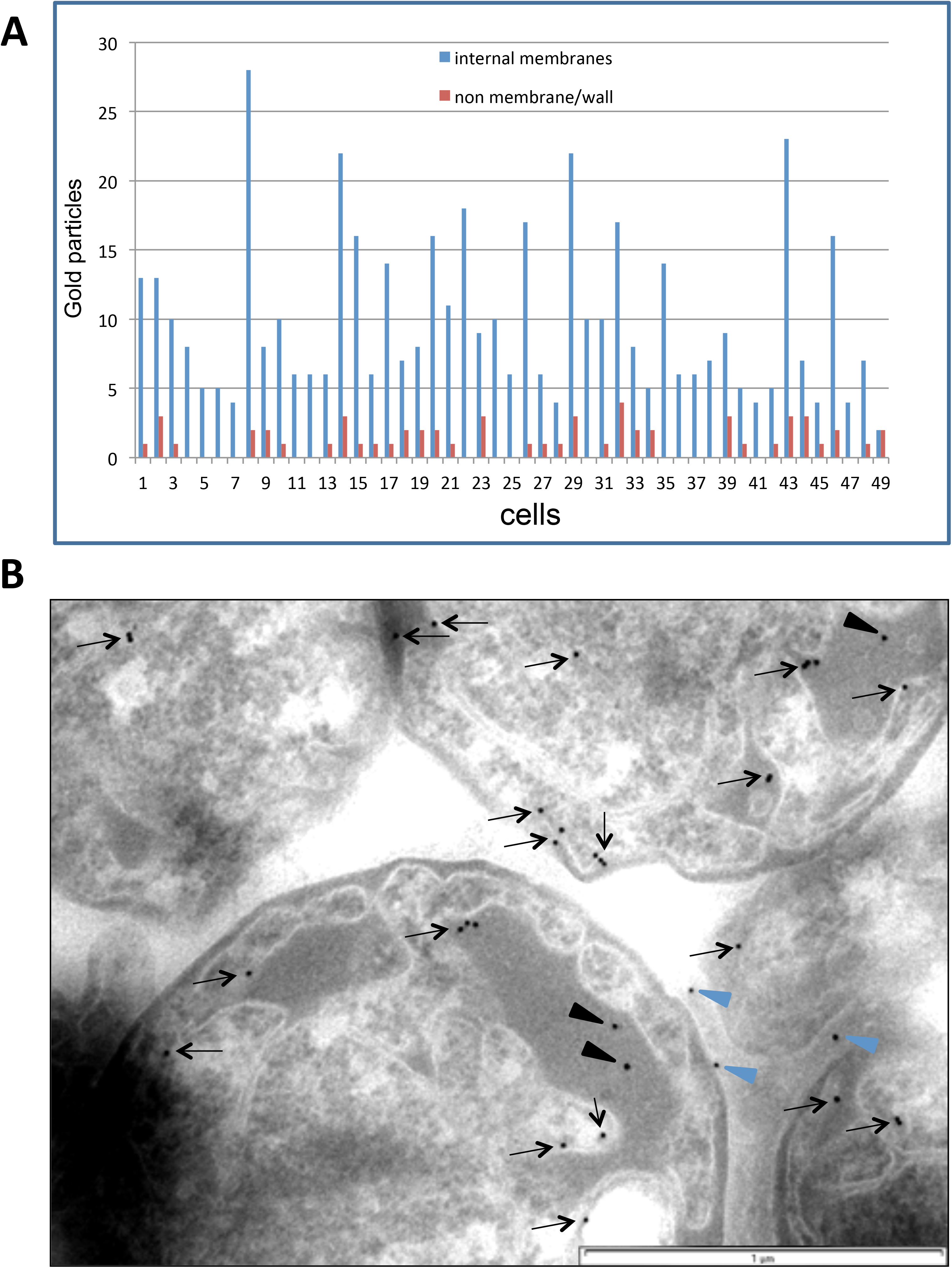
Immuno-gold particles distribution. (***A***) Distribution of gold particles within *Gemmata obscuriglobus* cells. The bars represent a number of particles associated with the intracytoplasmic membranes (blue bars) vs with no visible association with the membranes (red bars). A total of 50 cells were used for the counting; 549 particles were found as associated with the membranes and 60 as not associated. The particles were counted according to the description in (*B*). (***B***) An example of gold particle distribution within the cells of *Gemmata obscuriglobus*, labelled with 6670 antibody and then with 10 nm gold protein A. The majority of the particles were found associated with the intracytoplasmic membrane (black arrows). The particles were considered as membrane-associated if the distance between the membrane and the particle did not exceed 20 nm. Blue arrowheads indicate particles which were counted as a background or cell wall-associated, black arrowheads show non-membrane associated particles. Bar, 1 µm.

**S1 Table.**
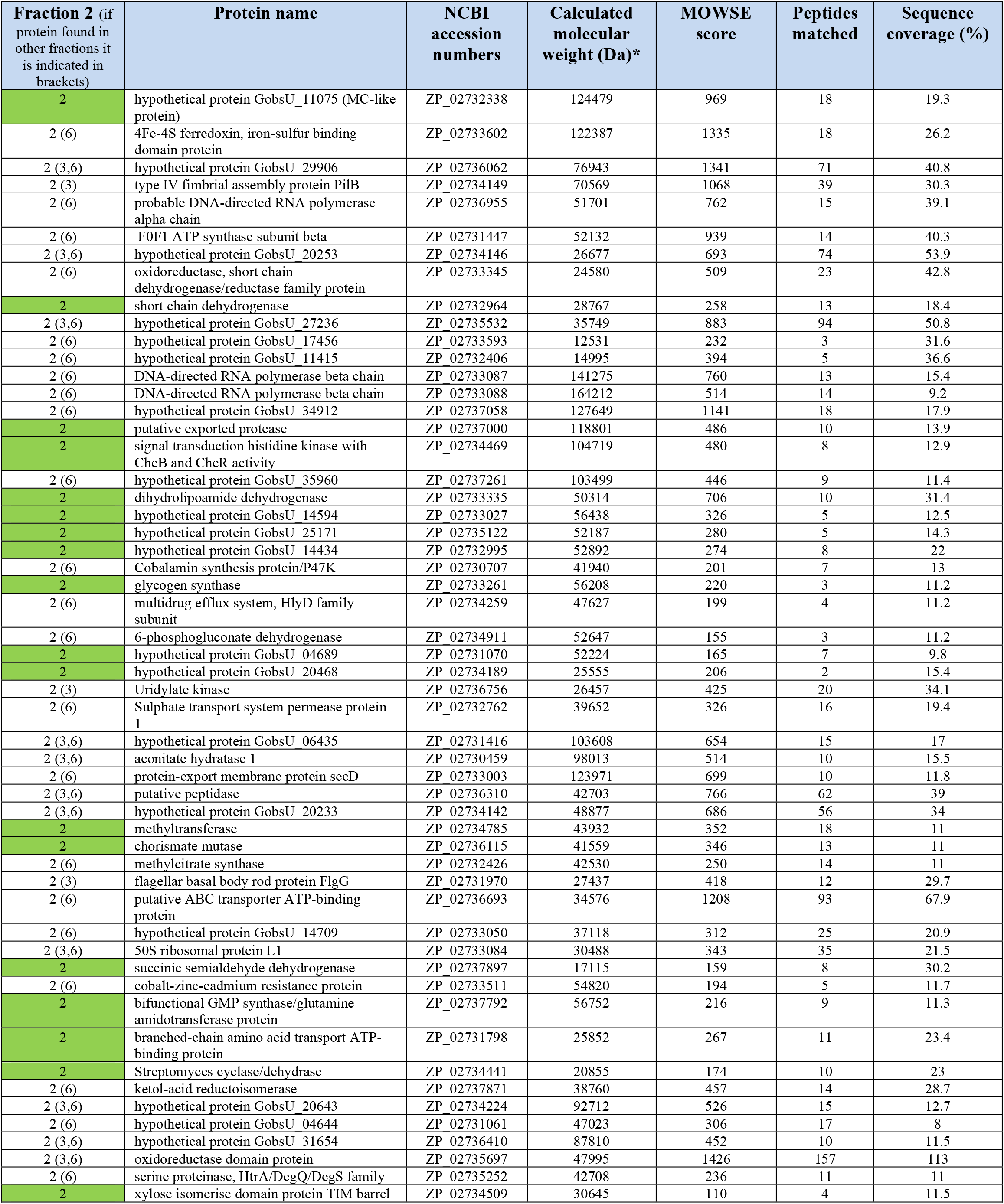
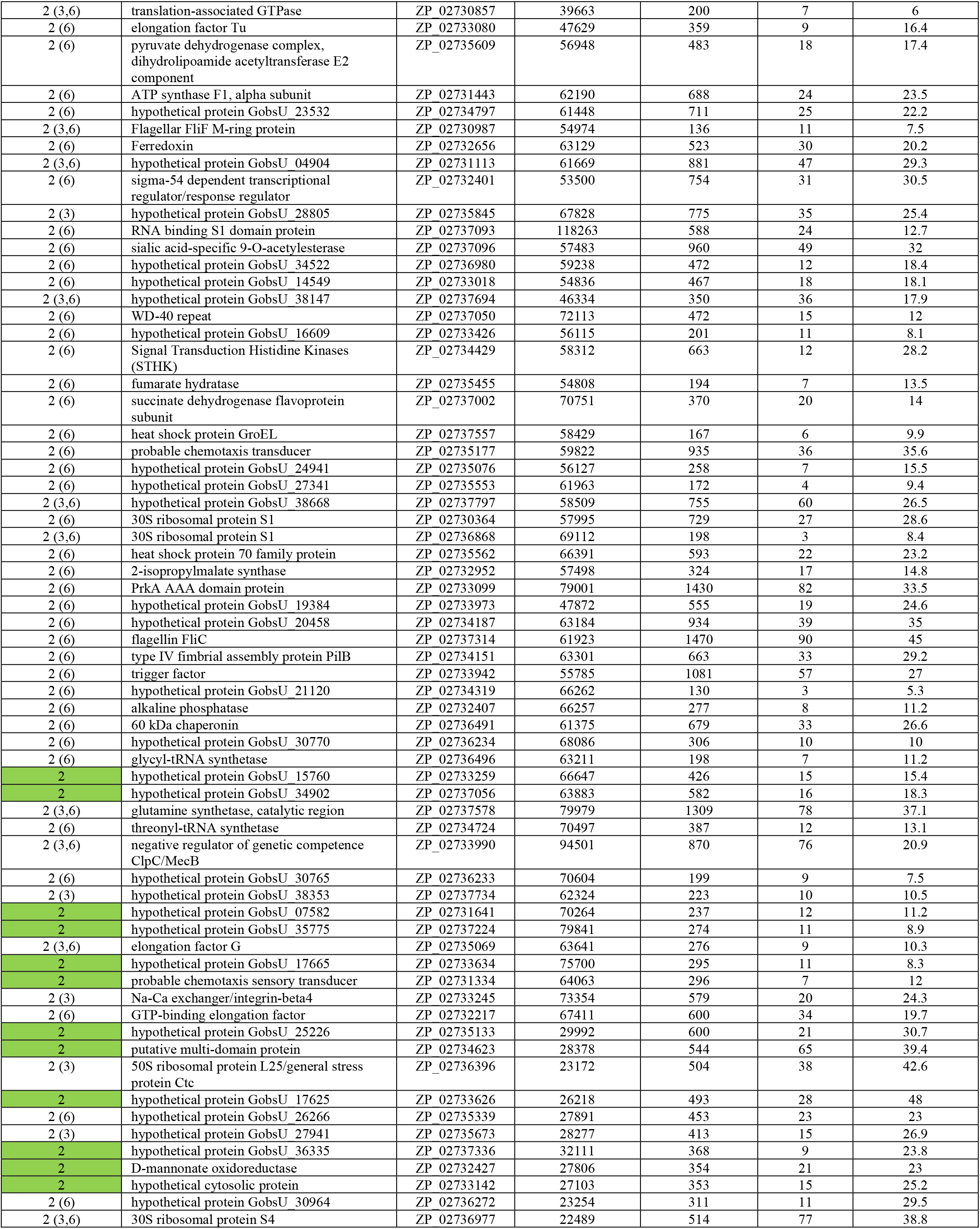
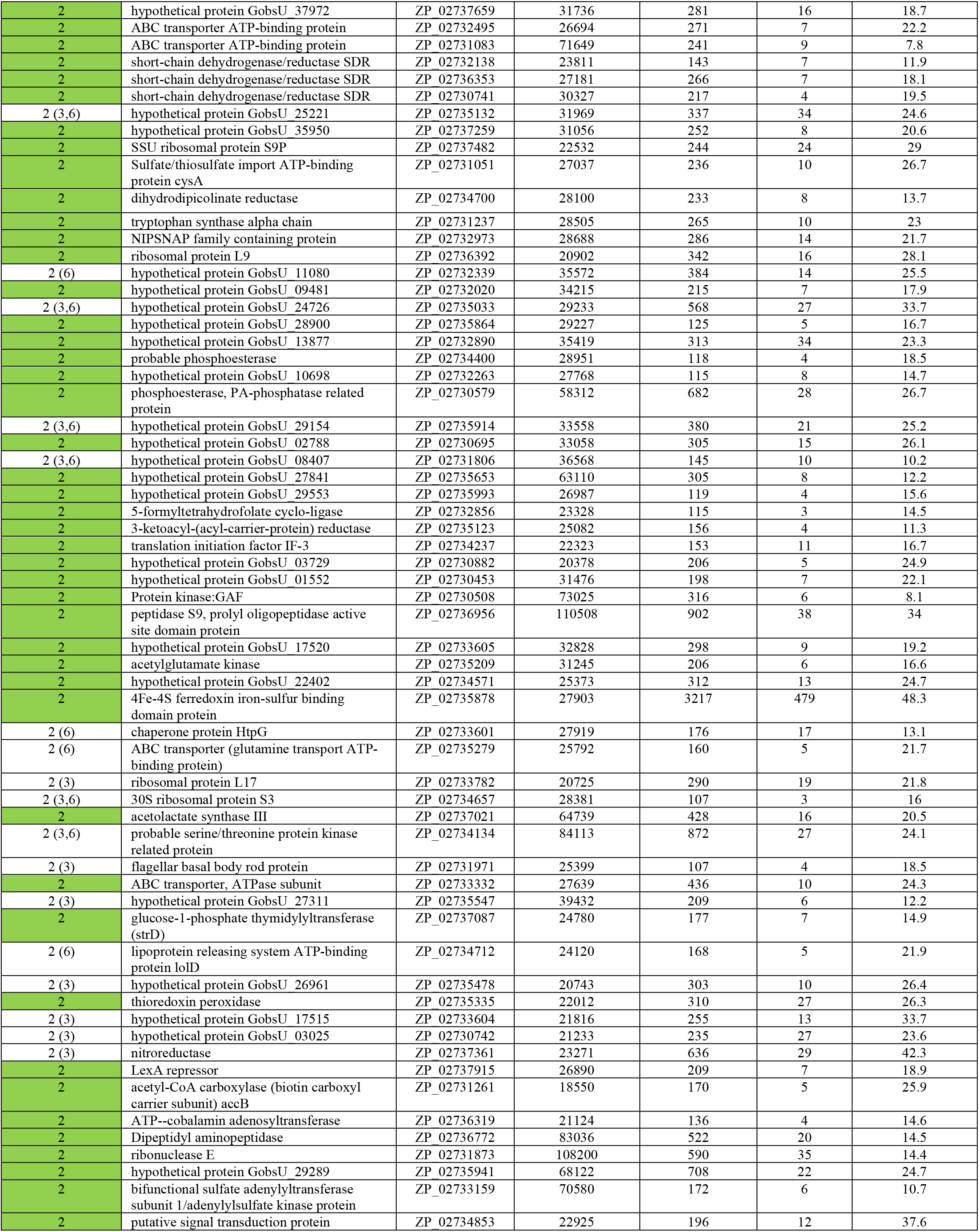
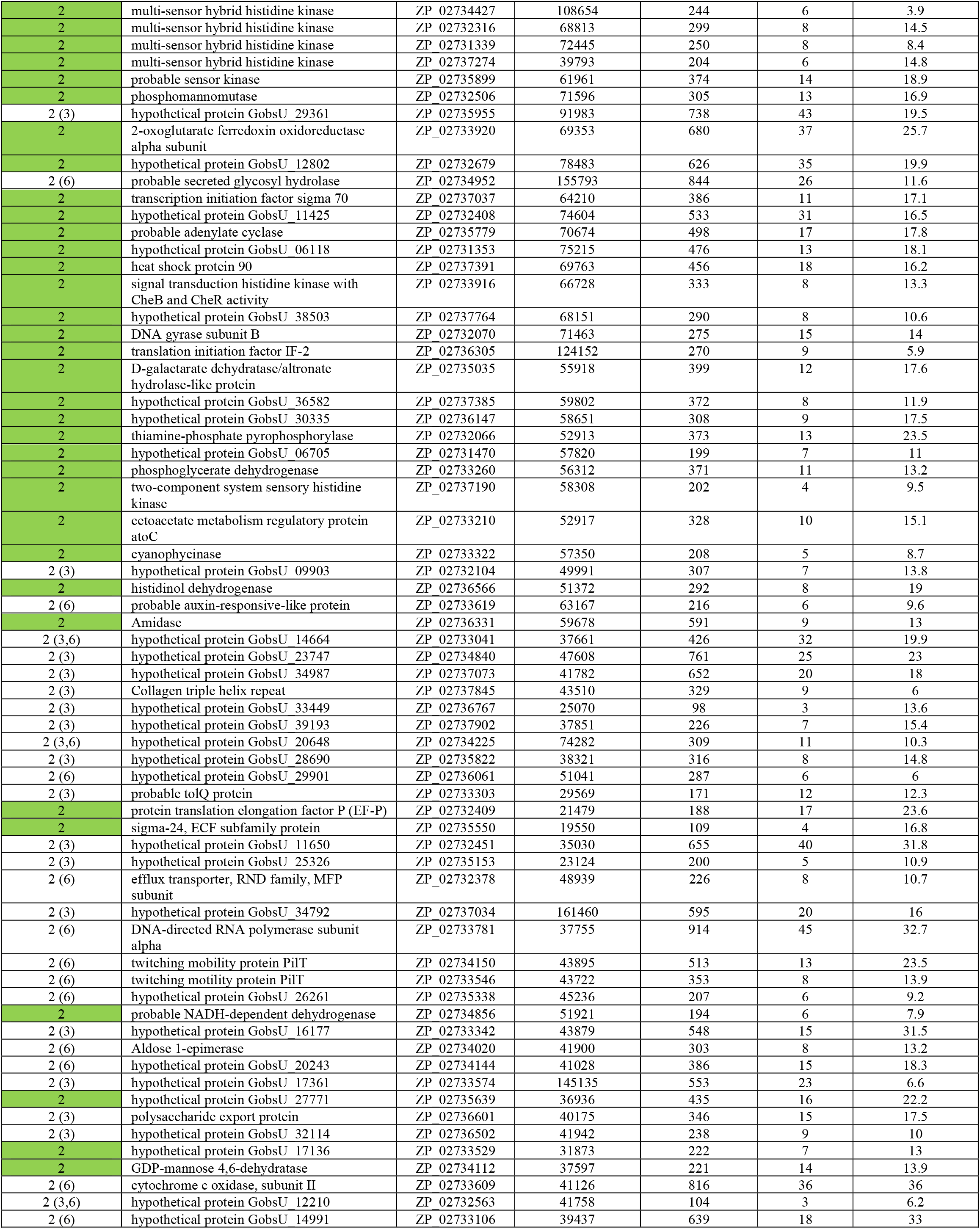
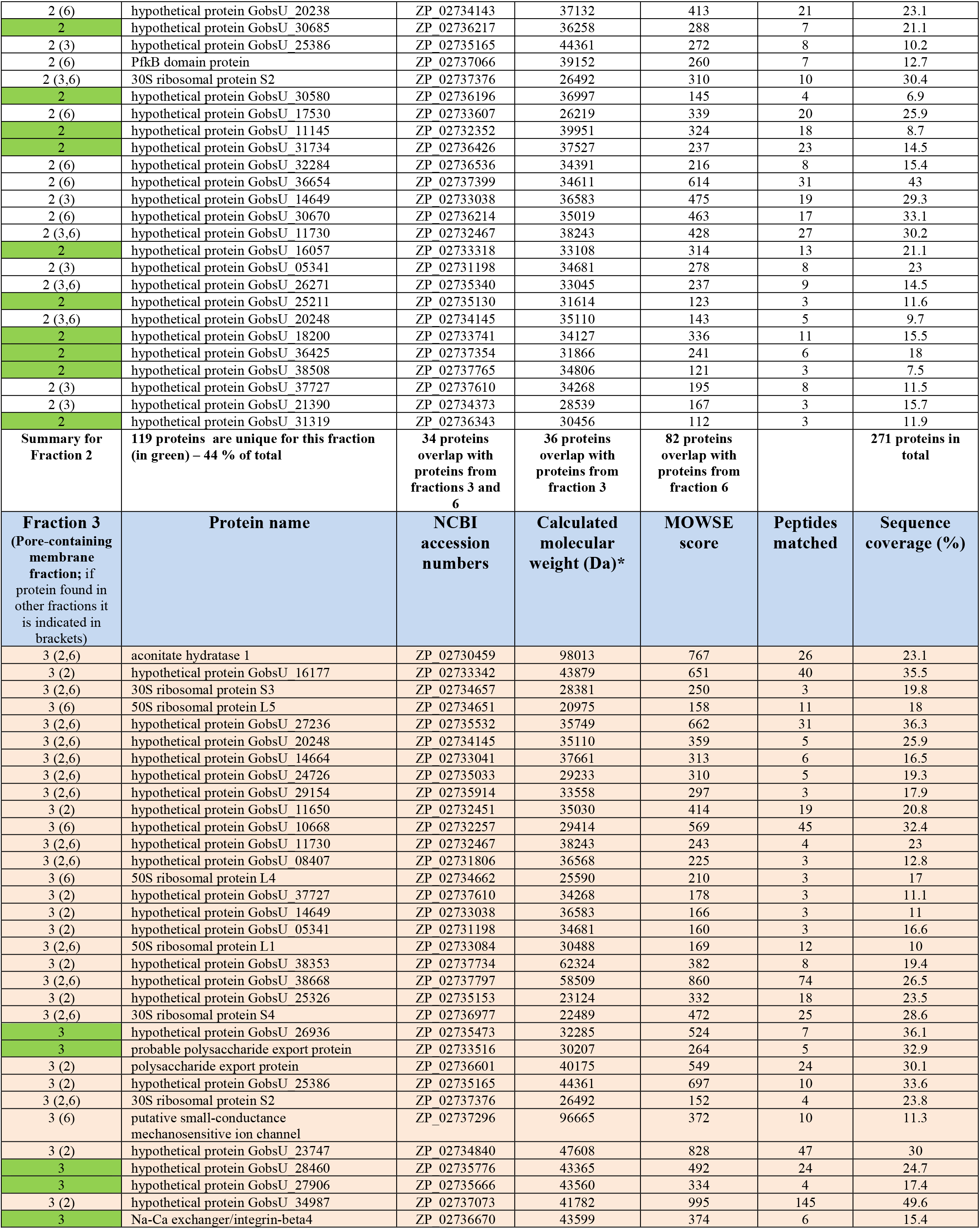
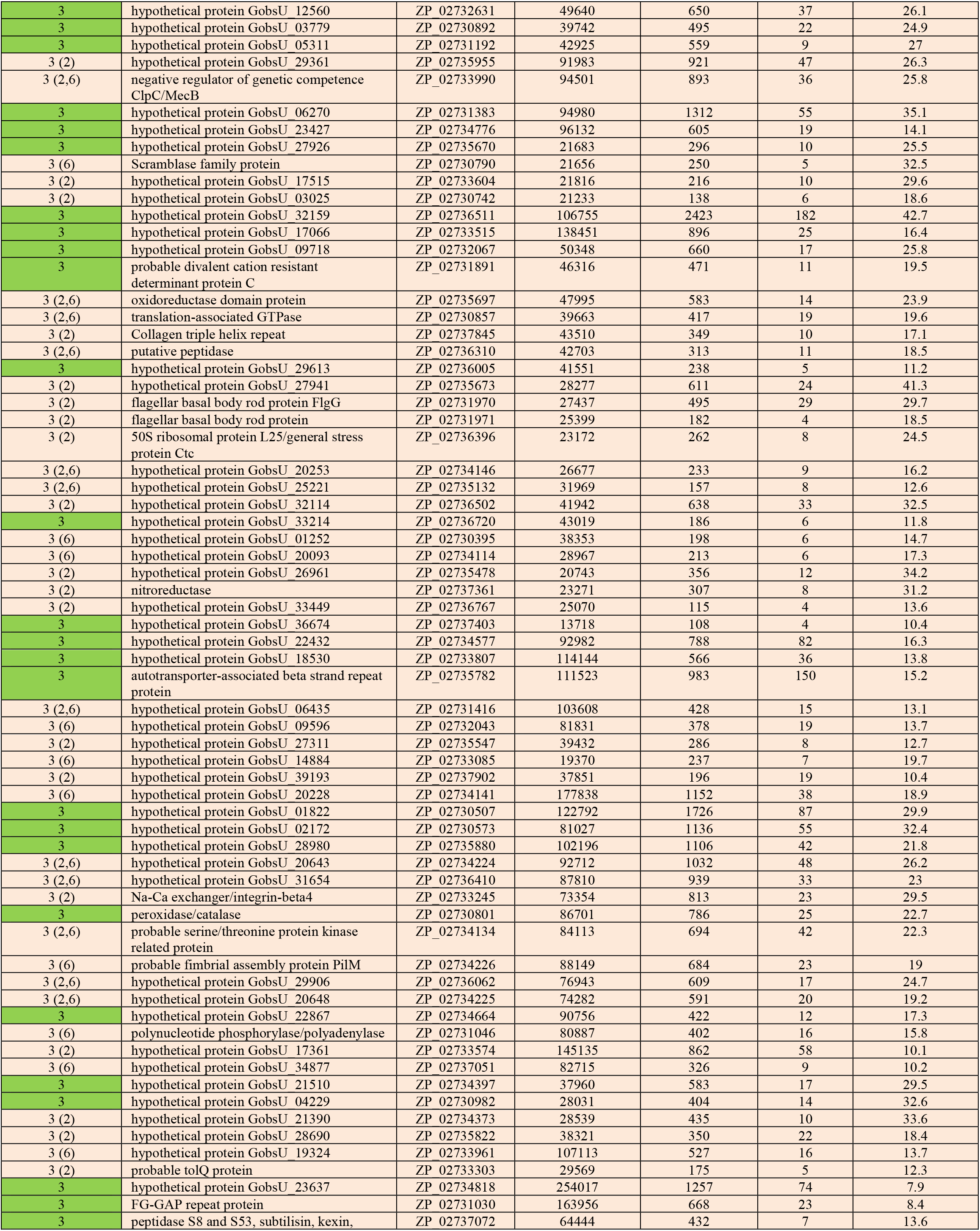
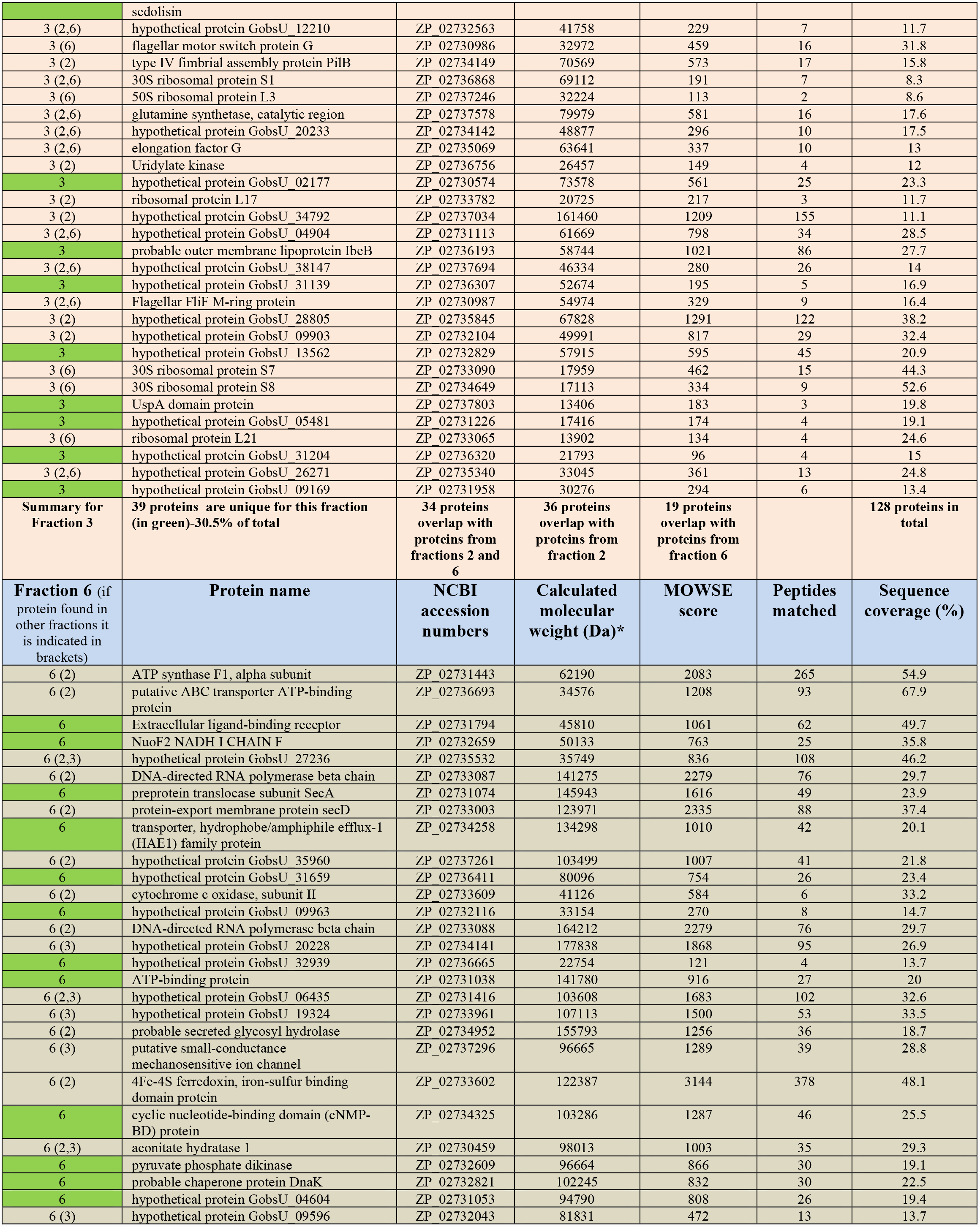
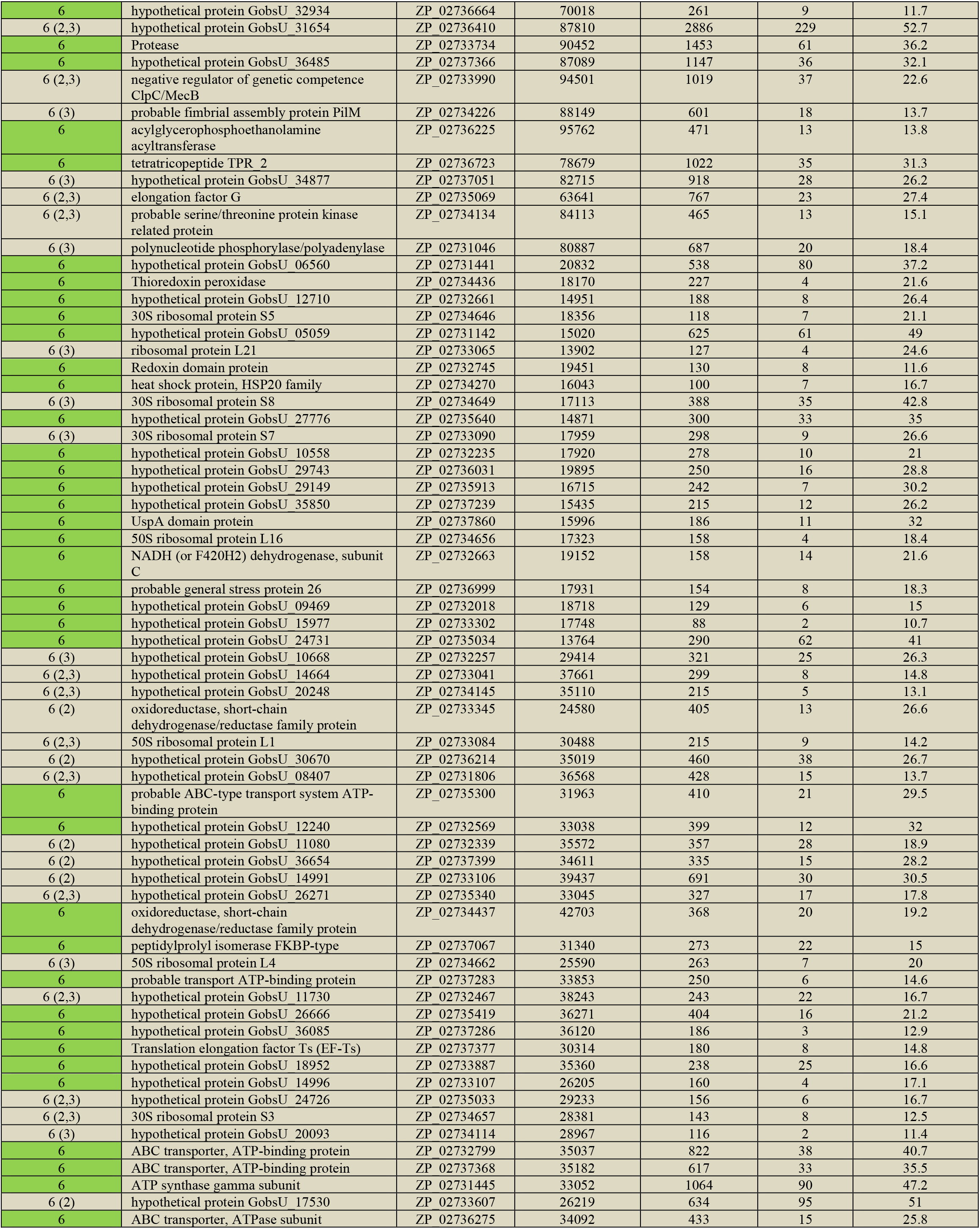
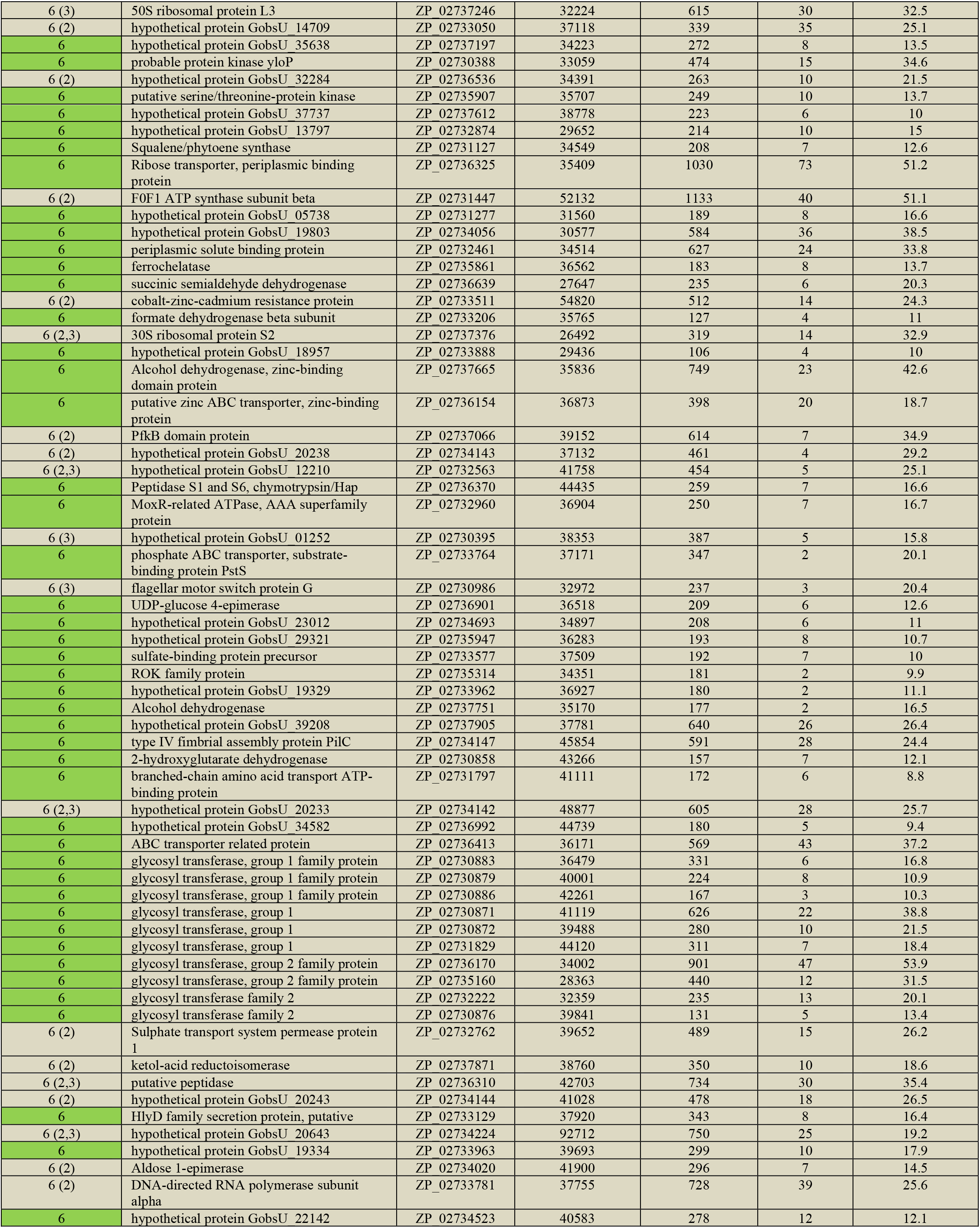
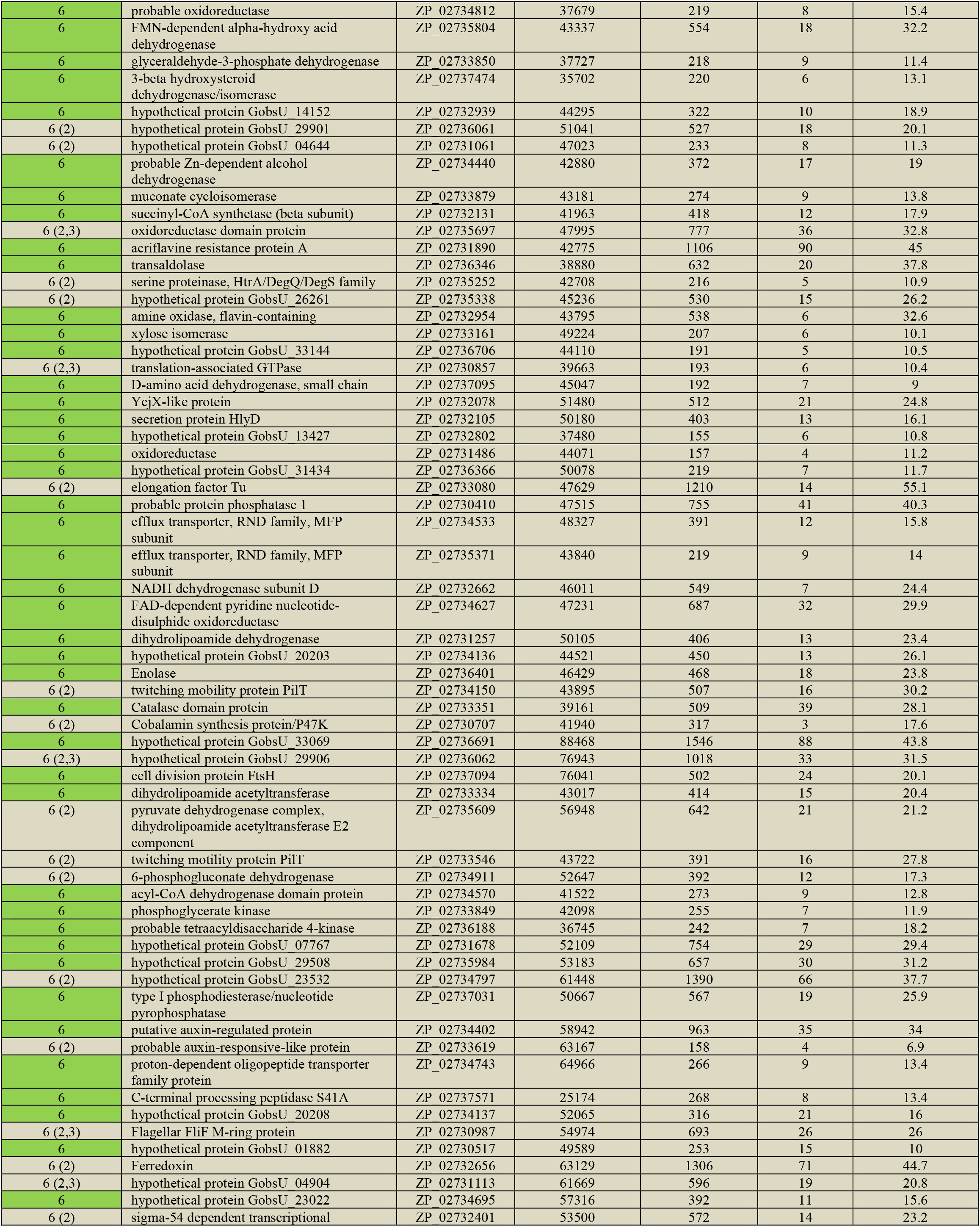
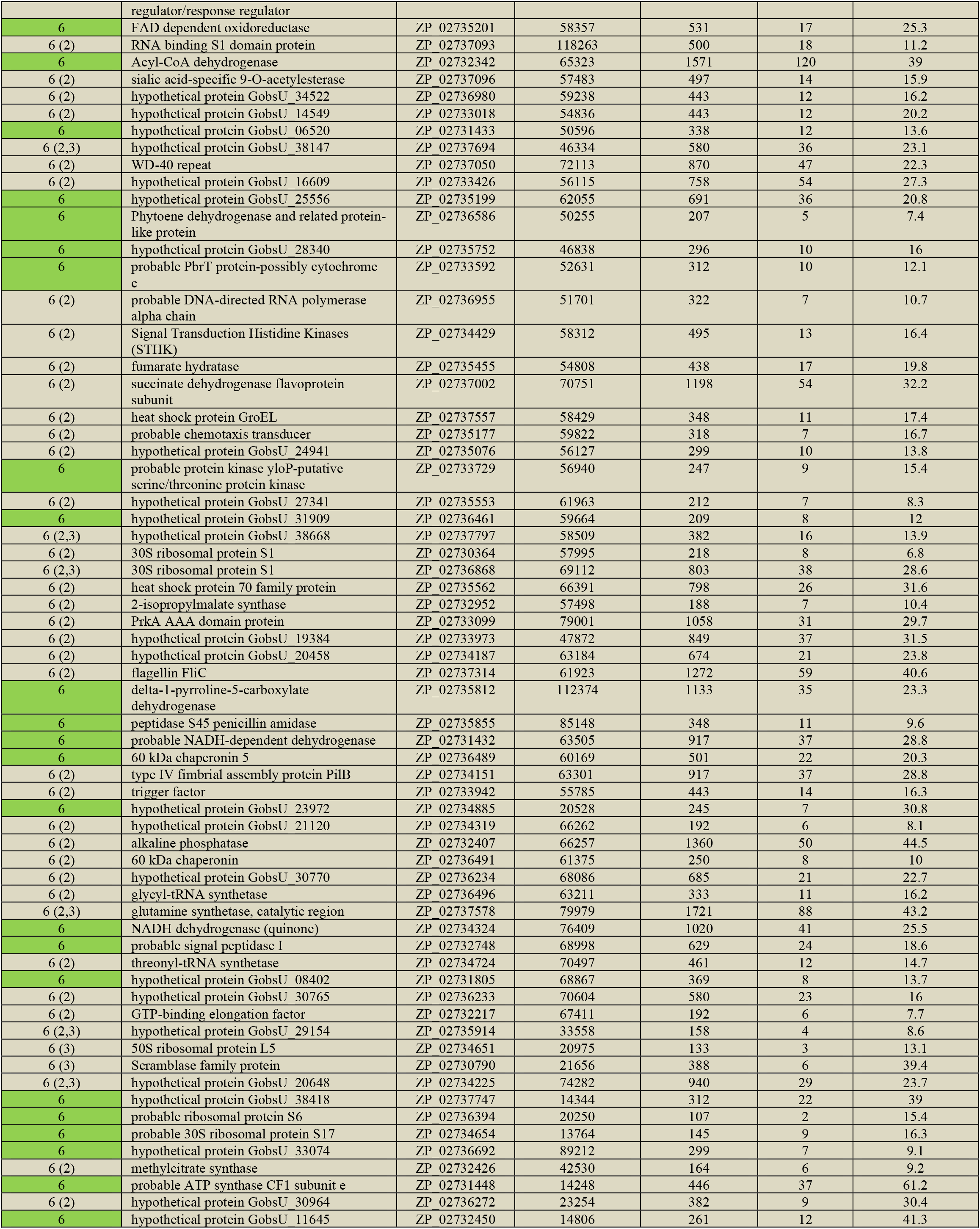
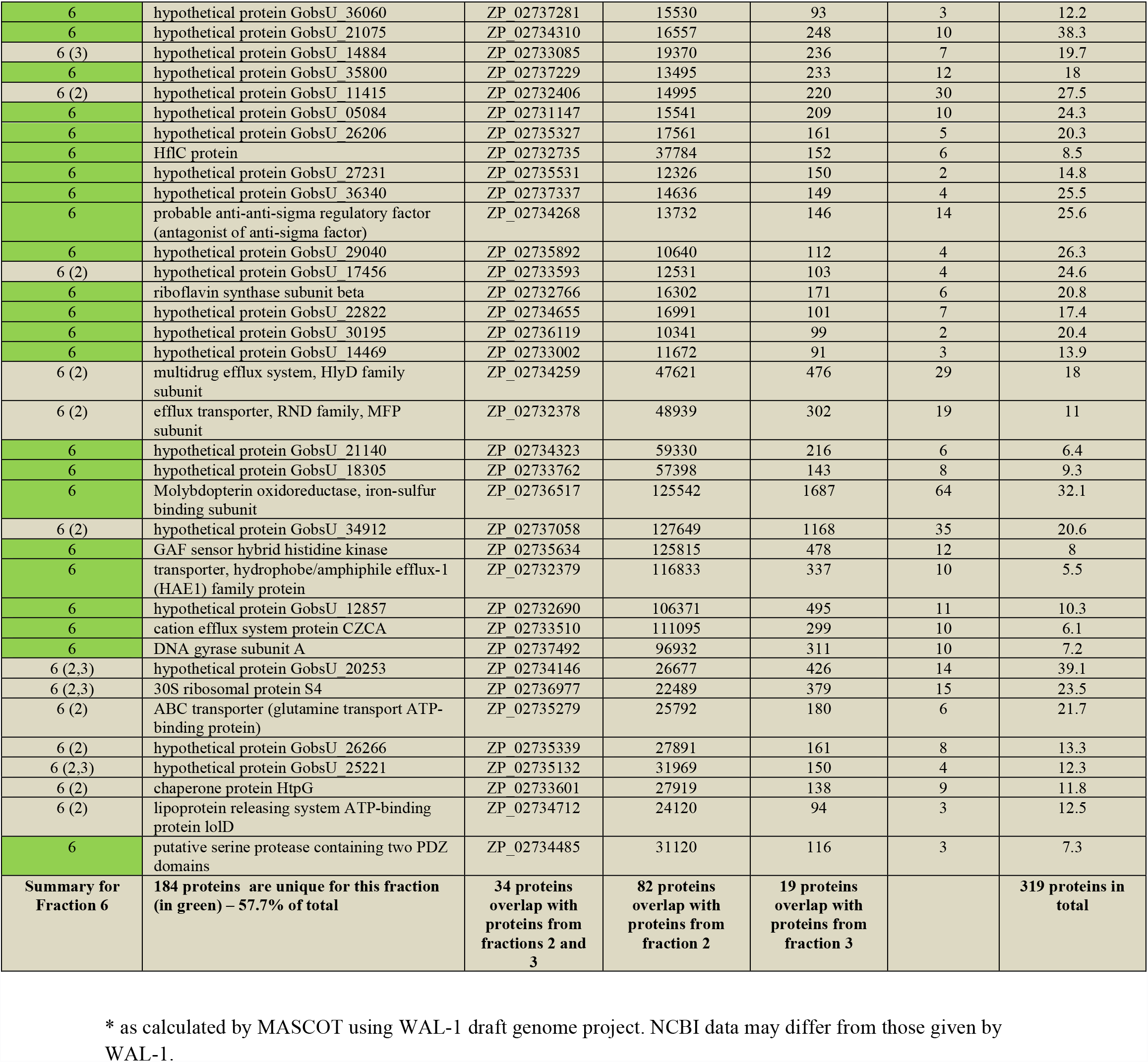
Proteins identified in membrane fractions by MALDI-TOF.

**S2 Table.** Summary of the membrane proteome analysis.

| **fraction** | **total**<br>**number of**<br>**proteins** | **proteins**<br>**with**<br>**predicted**<br>**function** | **hypothetical**<br>**proteins (unknown**<br>**function)** | **unique**<br>**proteins in**<br>**fraction** |
| --- | --- | --- | --- | --- |
| **2** | **271** | **150 (55%)** | **121 (45%)** | **119 (44%)** |
| **3 (pore**<br>**fraction)** | **128** | **44 (34%)** | **84 (66%)** | **39 (30.5%)** |
| **6** | **319** | **193 (61%)** | **126 (39%)** | **184 (58%)** |

**S3 Table.**
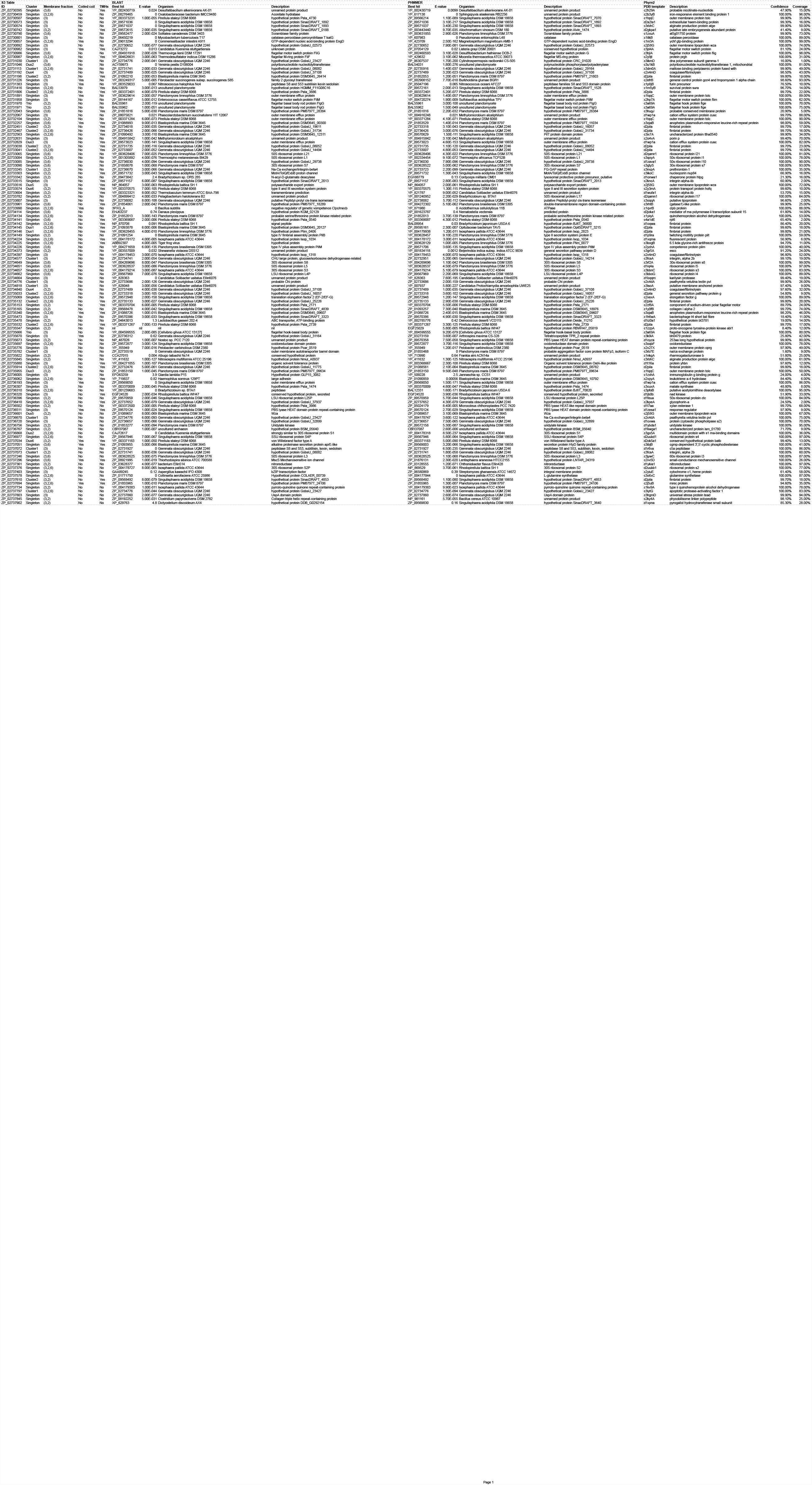

**S4 Table.** Results from structural analysis for the C-terminal region of cluster 1 (β-propeller) protein constituents^*^

| ID | Fraction | Confidence | Coverage | PDB template |
| --- | --- | --- | --- | --- |
| ZP_02731030 | (3) | 16.0% | 1% | 3IKM |
| ZP_02731113 | (2,3,6) | 99.9 | 52% | 2C4D |
| ZP_02733245 | (2,3) | 99.9% | 42% | 2C4D |
| ZP_02734577 | (3) | 98.6% | 25% | 2C4D |
| ZP_02734776 | (3) | 99.9% | 25% | 2C4D |
| ZP_02734818 | (3) | 40.1% | 5% | 3FCS |
| ZP_02735782 | (3) | 1.6% | 9% | 3RB7 |
| ZP_02736670 | (3) | 100% | 76% | 2C4D |
| ZP_02737072 | (3) | 99.8 | 39% | 2C4D |
| ZP_02737073 | (2,3) | 100% | 91% | 2C4D |
| ZP_02737797 | (2,3,6) | 99.9% | 49% | 2C4D |

\*Models generated from full sequences except ZP\_02734818 (see text). Best hits (shown) were chosen based on combined top confidence and coverage scores, and location in the alignable C-terminus (Figure supplement 13). Column 1, Genbank accessions. Columns 2 and 3, confidence and coverage scores as generated by Phyre2. Column 4, PDB template ID used by Phyre2 to generate models. Results below 95% confidence were not further analysed.

**S5 Table.** Results from structural analysis of cluster 2 (pili) protein constituents^*^

| ID | Fraction | Confidence | Coverage | PDB template |
| --- | --- | --- | --- | --- |
| ZP_02731198 | (2,3) | 99.7% | 21% | 1OQW |
| ZP_02731806 | (2,3,6) | 99.6% | 23% | 1OQW |
| ZP_02732451 | (2,3) | 99.7% | 24% | 1OQW |
| ZP_02732467 | (2,3,6) | 99.7% | 23% | 1OQW |
| ZP_02733038 | (2,3) | 99.6% | 25% | 1OQW |
| ZP_02733041 | (2,3,6) | 99.7% | 27% | 1OQW |
| ZP_02735033 | (2,3,6) | 54.8% | 8% | 2PIL |
| ZP_02735132 | (2,3,6) | 99.7% | 27% | 1OQW |
| ZP_02735532 | (2,3,6) | 99.6% | 20% | 1OQW |
| ZP_02735914 | (2,3,6) | 99.7% | 26% | 1OQW |
| ZP_02737610 | (2,3) | 99.7% | 26% | 1OQW |

\*Models generated from full sequences. Best hits (shown) were chosen based on combined top confidence and coverage scores. Column 1, Genbank accessions. Columns 2 and 3, confidence and coverage scores as generated by Phyre2. Column 4, PDB template ID used by Phyre2 to generate models.

**S6 Table.** Results from keyword analysis of Phyre output.

**outer membrane lipoprotein**
| ID | Fraction | Confidence | Coverage | PDB template |
| --- | --- | --- | --- | --- |
| ZP_02730892 | (3) | 100.00% | 95.00% | 2J58 |
| ZP_02731226 | (3) | 77.00% | 29.00% | 1JCC |
| **ZP_02733516** | (3) | 100.00% | 75.00% | 2J58 |
| **ZP_02736601** | (2,3) | 100.00% | 70.00% | 2J58 |

**outer membrane efflux protein**
| ID | Fraction | Confidence | Coverage | PDB template |
| --- | --- | --- | --- | --- |
| ZP_02730507 | (3) | 99.00% | 67.00% | 1WP1 |
| ZP_02731891 | (3) | 100.00% | 91.00% | 1WP1 |
| ZP_02732067 | (3) | 99.70% | 91.00% | 1WP1 |
| ZP_02732104 | (2,3) | 100.00% | 95.00% | 1WP1 |
| **ZP_02732829** | (3) | 100.00% | 88.00% | 1WP1 |
| **ZP_02735955** | (2,3) | 100.00% | 62.00% | 1WP1 |
| ZP_02736193 | (3) | 100.00% | 89.00% | 1WP1 |

**transmembrane beta barrel**
| ID | Fraction | Confidence | Coverage | PDB template |
| --- | --- | --- | --- | --- |
| ZP_02730574 | (3) | 99.90% | 58.00% | 3RBH |
| **ZP_02731192** | (3) | 98.30% | 53.00% | 1I78 |
| ZP_02732104 | (2,3) | 5.30% | 9.00% | 1QJ8 |
| ZP_02732631 | (3) | 99.40% | 78.00% | 2O4V |
| ZP_02734397 | (3) | 96.80% | 52.00% | 1I78 |
| **ZP_02734840** | (2,3) | 97.40% | 46.00% | 1I78 |
| ZP_02735776 | (3) | 97.90% | 60.00% | 2X27 |
| ZP_02735845 | (2,3) | 100.00% | 64.00% | 3RBH |
| ZP_02735880 | (3) | 97.60% | 11.00% | 1T16 |
| ZP_02735955 | (2,3) | 38.80% | 6.00% | 2MPR |

**porin**
| ID | Fraction | Confidence | Coverage | PDB template |
| --- | --- | --- | --- | --- |
| ZP_02730574 | (3) | 98.60% | 64.00% | 3SYB |
| **ZP_02731192** | (3) | 20.80% | 9.00% | 2O4V |
| ZP_02732631 | (3) | 99.40% | 78.00% | 2O4V |
| ZP_02734397 | (3) | 36.80% | 61.00% | 1T16 |
| **ZP_02734840** | (2,3) | 50.60% | 58.00% | 1T16 |
| ZP_02735776 | (3) | 76.20% | 34.00% | 2WJQ |
| ZP_02735845 | (2,3) | 100.00% | 64.00% | 2Y0K |
| ZP_02735880 | (3) | 97.60% | 11.00% | 1T16 |
| ZP_02735955 | (2,3) | 38.80% | 6.00% | 2MPR |

**tolc**
| ID | Fraction | Confidence | Coverage | PDB template |
| --- | --- | --- | --- | --- |
| ZP_02730507 | (3) | 99.90% | 61.00% | 1TQQ |
| ZP_02731891 | (3) | 100.00% | 89.00% | 1TQQ |
| ZP_02732067 | (3) | 98.40% | 82.00% | 1TQQ |
| ZP_02732104 | (2,3) | 100.00% | 99.00% | 1TQQ |
| **ZP_02732829** | (3) | 100.00% | 80.00% | 1TQQ |
| **ZP_02735955** | (2,3) | 100.00% | 60.00% | 1TQQ |
| ZP_02736193 | (3) | 100.00% | 77.00% | 1TQQ |

oprX
| ID | Fraction | Confidence | Coverage | PDB<br>template | Class |
| --- | --- | --- | --- | --- | --- |
| ZP_02730574 | (3) | 97.10% | 61.00% | 2ODJ | oprD |
| **ZP_02731192** | (3) | 96.50% | 48.00% | 2X27 | oprG |
| ZP_02732631 | (3) | 99.40% | 78.00% | 2O4V | oprP |
| ZP_02734397 | (3) | 91.00% | 40.00% | 2X27 | oprG |
| **ZP_02734840** | (2,3) | 58.20% | 39.00% | 2LHF | oprH |
| ZP_02735776 | (3) | 97.90% | 60.00% | 2X27 | oprG |
| ZP_02735845 | (2,3) | 99.90% | 64.00% | 2ODJ | oprD |
| ZP_02735880 | (3) | 78.90% | 5.00% | 2X27 | oprG |
| ZP_02737902 | (2,3) | 5.60% | 10.00% | 2LHF | oprH |

## Bioinformatics analyses

We performed a number of bioinformatics analyses on the 128 unique proteins identified through proteomics as belonging to the pore-containing membrane fraction (fraction 3). We first searched for similarity to known proteins in the non-redundant protein database (downloaded from NCBI) using BLASTP and PHMMER.

BLASTP (2) reported 112 proteins with significant homologs (E<0.001) (S3 Table). Of these, 33 are in the membrane fraction 3 (84.6% of all fraction 3 proteins). Of the significant hits, 25 top hits are to non-identical sequences in *G. obscuriglobus*, 18 are to *Planctomyces*, and 16 to *S. acidiphila*. One sequence found no hits at all (ZP_02735547). Almost half of all hits are to hypothetical (50), unnamed (5) or probable (2) proteins. Among hits with assigned function, 6 are flagellar, 12 ribosomal, 6 are membrane-related, and 4 are efflux proteins.

PHMMER (from the HMMER package, ver. 3.0(1)), returned significant hits for 115 proteins (E<0.001) (S3 Table), of which 21 are to *G. obscuriglobus*, 20 are to *Planctomyces*, and 21 are to *S. acidiphila*. The functional distribution is similar to the BLAST results: A large fraction are hypothetical (51), unnamed (7) and probable (3), while annotated functions include flagellar (6), ribosomal (13), efflux (4) and membrane (5) hits. Notably, PHMMER and BLASTP find the exact same top hits for 64/128 proteins. 17 are non-identical *G. obscuriglobus* proteins, 11 are from *Planctomyces*, and 14 from *S. acidiphila*.

As BLASTP and PHMMER screens both identified hits to other proteins coded in the *G. obscuriglobus* genome, we examined similarity among proteins from the pore-containing fraction. We performed all against all BLAST (2) followed by a Markov clustering on the results (3) as implemented in VisBLAST (4) with default parameters E<0.001 and i-value=2.0. This yielded two large clusters (both containing 11 proteins), and a number of small triplet and doublet clusters (Fig S13). The vast majority of proteins were singletons (91 proteins; S3 Table), and the clustering was robust at higher E-value cutoffs (up to E=10) and i-values (from 1.2 to 5). One of these large clusters contained 7 proteins that were unique to fraction 3 with the remaining four proteins being divided between fractions (2,3) and (2,3,6). The second large cluster contained no proteins that were unique to fraction 3. We performed the same clustering on the full set of all 512 proteins and the same clusters were recreated. VisBLAST was used to cluster proteins based on sequence similarity. An E-value cut-off of 0.001 was used together with an i-value of 2. Interestingly, the cluster containing 8 proteins unique to fraction 3 did not change at all in this larger analysis, which indicates that it is indeed membrane-specific. The other large cluster expanded with proteins belonging to fractions (2,6), (6) and especially fraction (2). Other large clusters were found within the set of 512 proteins, however none contained fraction 3 proteins (Fig S14).

Transmembrane helix structure potential was predicted for all 128 fraction 3 sequences using TMHMM2 (5). Of these, 42 showed significant signal of one or more transmembrane helices. Seven of these are unique to fraction 3 (S3 Table). We also looked for evidence of coiled coils using Paircoil2 (6). We report 15 proteins with significant signal of a coiled coil structure (S3 Table).

We next examined fold architecture of proteins in the membrane pore fraction using Phyre2 (7). This approach uses homology modelling to infer the structure of an amino acid sequence based on resemblance to known structures. 127 of the sequences were modelled in full, but one protein (ZP_02734818, 2558 aa) was analyzed in pieces because of its large size. The Phyre2 result for this protein is thus based on the best scoring subsequence. In the S3 Table we report the top Phyre2 structural hit for each protein. These hits are extracted automatically from the output, and we note that they may not represent the sole best hit (multiple hits often have the same highest confidence score). Based on PDB descriptions, these unfiltered results include membrane proteins (14), flagellar proteins (7) and ribosomal proteins (8), along with a number of other bacterial membrane/transport-related hits.

Comparing all structural predictions with our clustering analysis reveals a number of interesting patterns. Most significantly, cluster 1 consists of proteins modelled by Phyre2 as β-propeller-containing. This is notable given the presence of the β-propeller architecture in protein constituents of the eukaryotic nuclear pore complex (8). Of the 11 proteins in this β-propeller cluster, seven are unique to the pore-containing membrane fraction (3) (Fig S14 and S3 Table). The second large cluster (cluster 2) is dominated by pilins, and the proteins in this cluster mainly come from membrane fraction (3, 2, 6). Approximately half the structural predictions for singleton proteins showed significant structural similarity to porins and membrane proteins, ribosomal subunits, and flagellar proteins (S3 Table). The predicted triplet cluster contains flagellar proteins.

A closer investigation of the structure predictions for the 11 members of the cluster 1 showed that eight yield at least one structure prediction with a confidence >95%. In most cases, multiple predictions are made covering all parts of the sequences.

As constituency in a cluster does not establish whether all constituents share a common region of sequence similarity, we performed multiple sequence alignments across all members of both cluster 1 and cluster 2 (using MAFFT, option L-ins-i, (9). For cluster 1, all sequences displayed similarity in the C-terminal region. Fig S15 shows the alignment for the eight sequences for which we also obtained high confidence (>95%) structures with Phyre. We evaluated the full alignment using the T-Coffee CORE program (10), which shows a moderately robust 8-way alignment with a CORE-score of 69 (where 100 is perfect alignment). It is clear that the conservation is most pronounced in the C-terminal end of the sequences from around position 850 in the alignment. Indeed, if only the C-terminal part of the alignment is analysed using T-Coffee, the score increases to 81. This corresponds well with the observation that the majority of hits retrieved when searching the non-redundant protein database using both BLAST and PHMMER are also against the C-terminal ends.

With the aim of better characterising the commonalities of cluster 1, we focused on the structural predictions associated with the common C-terminal region. Note that there is some disagreement between the top hits shown in S3 Table (which was automatically generated) and those derived from the conserved C-terminal region (S4 Table). If both coverage and confidence scores generated by Phyre 2 are considered, the structures associated with the C-terminal region (S4 Table) emerge as the best hits. Some cluster 1 proteins also yield significant predictions for their N-terminal ends, but these are not in conflict with results from their respective C-termini, indicating these may be multi-domain proteins. In all 8 cases where a significant (confidence >95%) structure model is obtained, the C-terminal predictions are for β-propeller structures that overlap with the conserved C-terminal region of the sequences (Fig S15). Furthermore, Phyre2 modeled all 8 proteins to the same PDB template (2C4D), which we interpret as independent verification that these proteins share a common structural fold. These results are not due to extensive sequence similarity, as the overall sequence identity between the queries and the PDB template ranges from 13% to 19%.

Results from our structural analysis of cluster 1 are given in S4 Table, and structural models are depicted in Fig 7C (all structures are visualized using The PyMOL Molecular Graphics System, Version 1.4.1 Schrödinger, LLC).

For the second large cluster (Fig S14), we performed the same type of analysis. All can be aligned (Fig S16), 10 of 11 sequences have significant (>95% confidence) structure predictions, and in all cases the best hit in terms of both confidence and coverage (S5 Table) was modeled against PDB file 1OQW. The predicted structures are all very similar and consist of a single α-helix.

Two proteins (ZP_02735673 and ZP_02736511) show possible α-solenoid structures with stacked α-helices (Fig S18). Both are singletons in the cluster analysis, and both are present in membrane fraction 3 (the first is in both fractions 3 and 2, and the latter is unique to fraction 3). The left-hand structure in Fig S18 (ZP_02735673) models against alpha-solenoid structures in pdb with high (>99%) confidence, with models spanning >95% of the sequence. The model shown is based on 1OYZ, a hypothetical protein from *E. coli*, which is classified in SCOP as a member of the ARM repeat superfamily. Within the top 10 hits are structures that derive from Bacteria, Archaea and Eukaryotes, including clathrin adaptor core proteins (2VGL, 1W63). The right-hand structure in Fig S18 (ZP_02736511) contains two high confidence domain models. The N-terminal region models to the same alpha-solenoid structure as seen in the left-hand structure in Fig S18. This spans 30% of the protein sequence. In the adjacent central region, Phyre2 models a response regulator (top hit: 1ZES) with high (>99%) confidence.

A possible FG repeat-containing protein (ZP_02734840) was found in the pore-containing membrane fraction and in the total nuclear membrane proteome. With 5 FGs in the first 200 residues, this conforms to a recent definition of FG-repeat nucleoporin (11) but the C-terminal half of the protein models as a transmembrane beta-barrel protein.

We also performed a more general screen for bacterial transmembrane proteins among our structural predictions. To do this, we screened results for predicted structures of the following type: “outer membrane lipoprotein”, “outer membrane efflux protein”, “transmembrane beta barrel”, “porin”, “tolc” and “oprd/g/h/p/”. The same protein might have hits in more than one of these categories. For transmembrane beta barrels, we chose the best hit to a beta barrel spanning the membrane even if the term “transmembrane” was not used to describe that particular hit (however, for all proteins in this category at least one hit is called “transmembrane”). For all other categories we chose the best hit containing the specified keyword(s). S6 Table summarises these hits with their ID, membrane fraction, confidence score and coverage (as reported by Phyre2), and the PDB template used. For the oprx group we also list the specific class.

